# Decoupling glycation from mortality: glucose, but not methylglyoxal, reduces survival in zebra finches

**DOI:** 10.64898/2026.05.04.722681

**Authors:** Adrián Moreno Borrallo, Sarahi Jaramillo Ortiz, Christine Schaeffer-Reiss, Julie Zumsteg, Claire Villette, Dimitri Heintz, Astolfo Mata Betancourt, Jean-Patrice Robin, Anaïs Allak, François Criscuolo, Fabrice Bertile

## Abstract

Birds provide a unique model for ageing research, with greater longevity and slower senescence compared to mammals of similar size, despite a higher mass-adjusted metabolic rate and blood glucose levels than other vertebrate groups. While the effects of glucose, glycation, and advanced glycation end-products (AGEs) on ageing are well-documented in biomedical research, their impact on avian physiology and ageing remains poorly understood. Although birds may possess adaptations mitigating the potential detrimental effects of glucose, elevated glucose still predicts reduced lifespan, and protein glycation varies with age and can influence fitness-related traits, implying that glycation or AGEs accumulation may have relevant effects on avian longevity. In this study, we experimentally investigated how one year of dietary supplementation with glucose or methylglyoxal affects survival and physiology (metabolic rate, flying performance, and beak coloration) in captive zebra finches (*Taeniopygia guttata*). We reveal a significant increase in mortality and an elevated albumin glycation rate in glucose-supplemented birds. Plasma AGEs increased in both supplemented groups, but this and glycation did not help explain the increased mortality in glucose-supplemented birds. Additionally, we observed an age-related constraint on seasonal metabolic adjustment and an age-related decline in beak coloration. This secondary sexual trait was affected by both supplementation treatments. Furthermore, flight performance was lower in supplemented birds, particularly in the glucose group, during the peak mortality period, suggesting a broader deterioration in health. Thus, although glucose supplementation was more deleterious than methylglyoxal, the underlying mechanisms responsible for the increase in mortality remain to be elucidated in future studies.

## Introduction

Birds represent a fascinating model for aging research. They exhibit high mass-specific metabolic rates and blood glucose levels— among the highest observed in vertebrates (Polakof et al., 2011). According to the *pace of life syndrome* hypothesis (Wolf et al., 2007; Réale et al., 2010), such traits should impose physiological constraints predicting shorter lifespans and accelerated aging. However, birds defy this expectation, combining exceptional longevity with lower senescence rates than similarly sized mammals (Lindstedt and Calder, 1976; Speakman, 2005; Jones et al., 2008; Ricklefs, 2010). The pace of life syndrome hypothesis also posits that physiological, life-history, and behavioural traits co-evolve along a "fast-slow" continuum (Wikelski and Ricklefs, 2001; Ricklefs and Wikelski, 2002). Yet, recent comparative data challenge this prediction in birds: for example, plasma glucose levels and lifespan are not negatively correlated across species (Tomasek et al., 2019; Moreno-Borrallo et al., 2025). Moreover, the pace of life syndrome hypothesis has received mixed support (Royauté et al. 2018), suggesting that expected relationships among traits may be obscured by complex adaptations, diverse lifestyles and environmental contexts, particularly when comparing distantly related taxa (Montiglio et al. 2018).

A key mechanism potentially linking high blood glucose to aging is glycation — the non-enzymatic reaction of sugars with molecules such as proteins or DNA (e.g. Brownlee 2001). This process can impair protein structure and function (Rondeau and Bourdon 2011 for albumin), leading to the formation of Advanced Glycation End products (AGEs), a heterogeneous group of compounds that accumulate in tissues and contribute to diseases (Chaudhuri et al. 2018). AGE formation is amplified by oxidative stress, and they also promote proinflammatory feedback loops that increase oxidative stress, thus further increasing AGEs formation (Khalid et al. 2022; Twarda-Clapa et al. 2022 for a review). While these processes are well studied in humans and the conventional models used in biomedical research (Ulrich and Cerami 2001; Suji and Sivakami 2004), their effects in non-model species such as birds remain poorly understood (Duval et al. 2024).

Paradoxically, although birds exhibit blood glucose levels that would be pathological in mammals, diabetes is rarely reported in avian species, and when it does occur, it is usually secondary to other pathologies (Van de Weyer and Tahas 2024). This observation has led to the hypothesis that birds may possess adaptive mechanisms mitigating the deleterious effects of glucose, such as reduced glycation or AGE formation, or increased resistance to their deleterious impacts (Szwergold and Miller 2014a; 2014b). However, recent studies in zebra finches challenge this idea: as in mammals, higher glucose levels negatively predict lifespan, and reduced glucose tolerance is associated with increased mortality in older individuals (Montoya 2018; Montoya et al. 2022), though the underlying mechanisms remain unknown.

Few studies have examined protein glycation as a potential mediator of glucose effects on ageing (i.e., the decline in general performances with time) and other life-history traits in birds. While haemoglobin glycation rate is lower in birds than in humans (e.g. Rendell et al. 1985; Brun et al. 2022) – likely due to reduced glucose uptake in red blood cells (Bell 1957) – albumin glycation rate tends to be higher. Correlational evidence shows that haemoglobin glycation rate varies with age in cross-sectional analyses, positively predicts mortality (Récapet et al. 2016), and is associated with the number of fledglings produced in collared flycatchers (Andersson and Gustafsson 1995) and with growth rate in kestrels (Ardia 2006). In captive zebra finches, higher haemoglobin glycation rate has been reported in females after raising more chicks (Borger 2024), suggesting that blood protein glycation may mediate trade-offs relevant to ageing, such as the costs of reproduction or growth.

Regarding AGEs, several studies have reported positive relationships between specific compounds (notably pentosidine) and age in avian tissues (primarily skin; Iqbal et al. 1999; Klandorf et al. 1999; Fallon et al. 2006a; Fallon et al. 2006b; Cooey and Virginia 2008; Cooey et al. 2010; Dorr et al. 2017), while others have not (Le Souëf 2012; Campion 2015; Rattiste et al. 2015; Labbé et al. 2019). Moreover, the mechanisms linking AGEs accumulation to age-related decline in performance and fitness remain poorly understood. The presence of the receptor of AGE (RAGE), which mediates proinflammatory responses in mammals (Hofmann et al. 1999; Schmidt, et al. 2001), in avian vasculature is doubtful (Eythrib 2013; but see Cousens and Braun 2010), suggesting a potential resistance of birds to the proinflammatory effects of AGEs. Indeed, the gene (*Ager*) is thought to have emerged in mammals and to be absent from birds (Sessa et al. 2014). However, evidence exists for AGEs-induced proinflammatory responses via peripheral blood leukocytes in chicken (Wein et al. 2020), although further studies also suggest that AGEs-induced proinflammatory effects are much more limited and tissue-restricted than in mammals (Luo et al. 2025).

To disentangle the effects of glucose on glycation and AGE formation, and their consequences for ageing patterns in birds, experimental studies are required. Yet, few have investigated how experimentally increasing these parameters affects individual fitness and age-related processes. Most dietary glucose supplementation experiments have been conducted in poultry science, focusing on agriculturally relevant traits, such as changes in body composition and weight gain (e.g. Brambila and Hill 1967; Shapira et al. 1978; Jiang et al. 2008), but with little attention to ageing-related outcomes. Furthermore, most studies have reported no effect of dietary supplements on blood glucose levels (e.g. Basile et al. 2022).

In this study, we experimentally examined how one year of dietary supplementation with glucose or methylglyoxal — a highly reactive dicarbonyl compound considered by some authors as one of the major pro-glycation agents, and perhaps the most researched one (Rabbani and Thornalley 2014; Sibbersen et al. 2018; Nigro et al. 2019), known to promote the formation of AGEs, such as CEL (carboxyethyl lysine) or hydroimidazolones (e.g. MG-H1) — affects survival and age-related mechanisms in captive male and female zebra finches of known age. Methylglyoxal is naturally produced in living organisms via retro-aldol condensation and auto-oxidation of monosaccharides or early glycation products, degradation of trioses phosphate (from glycolysis), and oxidation of lipids and other compounds (Thornalley et al. 1999; Lange et al. 2012). Increased glucose intake would therefore enhance endogenous production of reactive dicarbonyls, such as both methylglyoxal and glyoxal (Aragno and Mastrocola 2017; Xiaodi Zhang et al. 2023). Our aim was thus to test whether glucose supplementation promotes ageing through increased glycation and/or AGEs formation, specifically measuring CML (Carboxymethyl lysine) and CEL, two of the most commonly assessed AGEs molecules (see **Figure 1**). Glucose supplementation was expected to promote the formation of both CML (via glyoxal) and CEL (via methylglyoxal), while methylglyoxal supplementation was expected to promote only CEL formation. Methylglyoxal supplementation was also used in parallel to increase AGE levels independently of glucose, enabling us to separate AGEs-specific effects from other physiological effects of glucose.

**Figure 1.**
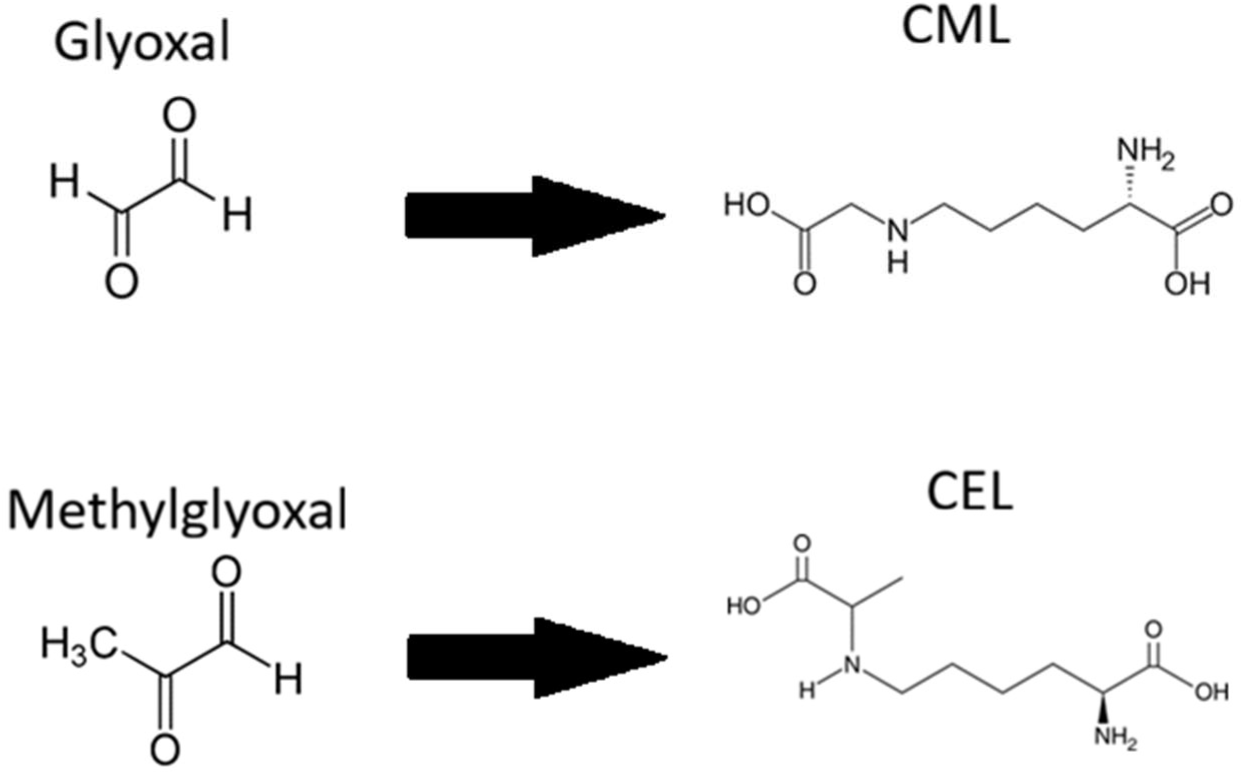
Different compounds for which we assessed free plasma concentration (see **Methods**). Both glyoxal and methylglyoxal are reactive dicarbonyl compounds originated (mostly) from glucose degradation or metabolism that lead to the formation of AGEs like CML and CEL, respectively. These compounds are metabolically inactive and can only be eliminated by renal filtration at a limited rate.

To evaluate the impact of glucose and methylglyoxal supplementation on health and aging, we monitored changes in multiple physiological and behavioural parameters. We hypothesized that both treatments would increase markers of cellular and tissue damage while impairing flight performance, body mass, and overall body condition. We specifically assessed resting metabolic rate, a well-established marker of physiological ageing in zebra finches that declines with age (Moe et al. 2009; Briga and Verhulst 2021). We predicted that glucose supplementation would initially elevate metabolism and the respiratory exchange ratio (RER) due to a greater carbohydrate use, followed by a more rapid metabolic decline compared to the control group. In contrast, methylglyoxal supplementation was expected to induce metabolic decline without the initial transient increase.

Although proper reproduction did not occur during the experiment, we measured beak coloration (a sexually selected secondary sexual trait; Simons and Verhulst 2011), as an additional indicator of individual quality. Beak coloration may indeed indirectly reflect reproductive senescence (Simons et al. 2012), providing further insight into age-related changes. To explore potential mechanisms underlying treatment effects, we also analysed the body composition of individuals that died during the experiment, testing for fat and water content. Increased adiposity is a phenomenon previously reported in glucose-supplemented chickens (Shapira et al. 1978; Jiang et al. 2008). Excess fat deposition could lead to hepatic steatosis, as observed in non-alcoholic fatty liver disease (Chao et al., 2019; Powell et al., 2021), or ectopic lipid accumulation in muscles, which may impair metabolic function in a manner analogous to diabetes and metabolic syndrome (Grundy, 2016). Such effects could be mediated by increased sugar intake in the glucose-supplemented group or by enhanced fatty acid synthase (FAS) activity due to methylglyoxal (Garrido et al. 2015; Moraru et al. 2018). Finally, water content assessment serves us for determining potential dehydration effects resulting from treatment-related differences in drinking behaviour (see **Pilot study section on ESM1 & 2**). However, as desert birds, zebra finches are relatively resistant to water deprivation and can face dehydration through different mechanisms, including reductions in skin water permeability, maintenance of high body water content, efficient production of metabolic water from fat reserves and enhanced evaporative heat loss efficiency (e.g. Menon et al. 1989; Rutkowska et al. 2016; Wojciechowski et al. 2021).

## Materials and methods

### Study species

Zebra finches (*Taeniopygia guttata*; Vieillot, 1817) are passerine birds of the family Estrildidae, originating from Australia and the Lesser Sunda Islands in Indonesia. The Australian subspecies is the source of all the domestic strains currently used in research laboratories. Zebra finches are relatively small (∼15 g), colourful, highly gregarious granivorous bird, and the Australian subspecies is also known as a desert dweller. Owing in part to their ease of captive breeding, zebra finches have long been used as a model species in avian research (Griffith and Buchanan 2010). They are particularly relevant for studies of coloration and sexual selection, given their sexual dimorphism (Morris 1954), and ecophysiology, (e.g. Veasey et al., 2001; Bertrand et al., 2006; Criscuolo et al., 2008; Tschirren et al., 2009; Criscuolo et al., 2011; Crino et al., 2014; Reichert et al., 2014; Reichert et al., 2015; Briga, 2016). Their sequenced genome (Warren et al. 2010) further facilitates their use in omics approaches. Several characteristics also facilitate their captive rearing, including their gregarious nature, which allows housing them in large aviary groups, as well as their opportunistic breeding habits and early sexual maturity (see Zann 1996). It remains relevant to mention, though, that some concern has been raised about the use of zebra finches as a model species, particularly given some of the same peculiarities in terms of life-history characteristics that makes them easy to use in captive studies, like their very rapid development and high fecundity, despite their life-span not being apparently shortened (Griffith et al. 2021).

Finally, the availability of a permanently maintained captive population at the DEPE (Department of Ecology Physiology and Ethology) in Strasbourg, motivated their use as a model species in our study. In addition to individuals already present in the DEPE’s animal facility, the study population was completed in 2021 and 2022 by birds originating from several sources. Most originated from the Max Plank Institute for Ornithology (including many of the ones already present at the DEPE), currently part of the Max Planck Institute for Biological Intelligence (Pöcking-Seewiesen, Bavaria, Germany), with additional individuals from Bielefeld University (Bielefeld, Germany), and a small number from pet shops. The birds from the Max Plank Institute originated from several breeds, predominantly Cracow, Seewiesen and Davis (see Forstmeier et al. 2007). The remaining of our birds were either from the Bielefeld breed or of unknown breed. Breed and source were not significantly biased across treatment groups (tested by Fisher’s exact test for count data; P = 0.552). Further details are provided in **ESM 3** (**Table ESM3.3**). Although domestication has led to some loss of genetic diversity in zebra finches, no major bottlenecks have occurred, and domesticated zebra finches still retain relatively high levels of genetic diversity compared with some other domesticated birds (Forstmeier et al. 2007). The inclusion of birds from different breeds and sources therefore contributed to a broader genetic background in our experimental population, which, together with the seminatural conditions (see below), may enhance the generalizability of our findings. We further tested if breeds influenced the measured variables using GLMMs including breed as a fixed factor and bird ID as a random factor, followed by pairwise comparisons using **`emmeans()`**. Results of these models are briefly discussed in **ESM1**.

### General protocol

A sample of 90 adult zebra finches (*Taeniopygia guttata*; Vieillot, 1817; see **study species section**) identified with numeric or alphanumeric codes on leg rings, was randomly assigned to three experimental groups: (i) control, (ii) glucose supplementation and (iii) methylglyoxal supplementation. Each group consisted of 15 males and 15 females (sex determined on their sexual dimorphic plumage; see **study species section**). Most individuals (86 out of 90) were of known age and belonged to two main cohorts: birds approximately 4 years old and birds approximately 6 years old (see detailed ages on **Table ESM3.2**), with a smaller number of younger birds (2-4 years old). Although a maximum lifespan of 14.5 years has been reported for zebra finches (Tacutu et al., 2018), typical lifespans are considerably lower, with these birds generally living around 5 years in the wild (Zahn, 1996) and usually less than 9 years in captivity (Briga et al., 2019; Jimeno et al., 2018; Montoya et al., 2018). Our birds, aged 2–6 years, represent a young to middle-aged cohort where early signs of senescence are detectable (Briga et al., 2019; Moe et al., 2008). This age range is appropriate for capturing the onset of aging-related declines under glucose and methylglyoxal supplementation. Thus, most of our birds, aged 2–6 years, can be considered as ranging from young adults to early senescent individuals. Consistent with this classification, previous studies addressing age-related variation in zebra finches have used birds of about 4 years (our ’young’ cohort) as an old cohort (Salmón et al., 2022), supporting the use of even older birds to study aging-related trait variations and the onset of senescence. Ages were distributed as equally possible across the treatment groups. They were placed into outdoors aviaries (380 x 215 x 240 cm; depth, width, height) in the IPHC animal facilities (Strasbourg, France), allowing them to perform exercise, avoiding some of the potential deleterious effects due to excess energy intake from supplemented diets. The photoperiod, temperature and humidity varied naturally, and were recorded by a thermometer/hygrometer placed in each aviary to register current, minimum and maximum temperature and humidity, taking measurements once a day in the afternoon (data shown in **ESM4**). Separation of sexes within the aviaries was not possible due to space limitation constraints. However, housing both sexes together makes our bird population closer to natural conditions. Food, consisting into a commercial seed mix (Deli Nature® for tropical finches), water (tap water with the adequate treatment, see below), cuttlebone and grate were provided *ad libitum*. Both food and water were provided in one feeder each, placed on the floor, so that all the birds could physically access to them easily, regardless of their health status. The glucose and methylglyoxal treatments were provided in the drinking water at 50 g L^-1^ for glucose (Sigma Aldrich G8270) and 8.33 g L^-1^ for methylglyoxal (prepared by diluting 7 mL of Sigma Aldrich M0252 40% methylglyoxal solution in water into 993 mL of tap water). The control group received plain water only. Treatment concentrations were determined based on the results from of a pilot study performed in February-March 2022, before the start of the main study (see **ESM 1 & 2**). The final dosages were selected based on the minimum amount concentration needed required to induce a significant change in plasma methylglyoxal concentrations. For glucose, as none of the tested supplementation levels altered plasma glucose concentrations, we selected the minimum lowest concentration tested to avoid imposing an unnecessary stress on the birds. The pilot study also allowed us to determine the final dilutions, given the observed variation in drinking rates as a function of the methylglyoxal concentration in the supplemented water. Water feeders were refilled once a week, or earlier when necessary. Further details about housing conditions are explained in **ESM2**. Whenever an individual died during the duration of the experiment, the body was collected, weighed and stored at -20 °C until dissection could be performed for body composition analyses. The date of death was recorded for survival analyses to be performed, along with comments on the possible causes or related circumstances. Some autopsies were performed to better inform the cause of death.

### Measurements and schedules

The experiment began on the 21^st^ of September 2022 and finished the 25^th^ of September 2023, with a starting population of 90 zebra finches, of which 4 were reused from the pilot study and 48 died over the course of the study. Measurement sessions were conducted every three months (i.e. August and November 2022 and February, May and August 2023; see **Figure 2.A**). During each session, several parameters related to birds’ senescence were measured, including body mass and body condition (fat and muscle scores), resting metabolic rate (RMR) and respiratory exchange ratio (RER) using indirect calorimetry (i.e. respirometry), beak coloration and flying performance. Whole blood glucose was assessed at every session except November. Plasma glucose, plasma albumin glycation rate, and plasma AGEs concentrations were assessed every six months (i.e. August 2022, February and August 2023). A detailed description of the methods employed for the determination of all of these variable is provided in **ESM2**.RMR flexibility, defined as the ability to change RMR depending on the environment, especially across seasons (e.g. McKechnie 2008; Norin and Metcalfe 2019; Swanson et al. 2022), was also measured, using the AUC (Area Under the Curve) of all the RMR values obtained during the whole experiment for each bird, and the amplitude of the change in RMR values across measurement sessions (see further details in **ESM2**).The less intrusive procedures, such as respirometry, flying test, weighing, coloration and body condition assessment, were performed during the first two weeks of every measurement session, while blood was sampled twice in the subsequent weeks (∼100 µL each week; see **ESM2**). Each measurement session, lasted a maximum of four consecutive weeks **Figure 2.B**), reduced to three when possible, due to a lower amount of remaining birds (i.e. birds deaths occurring along the experiment would reduce sample size).

**Figure 2.**
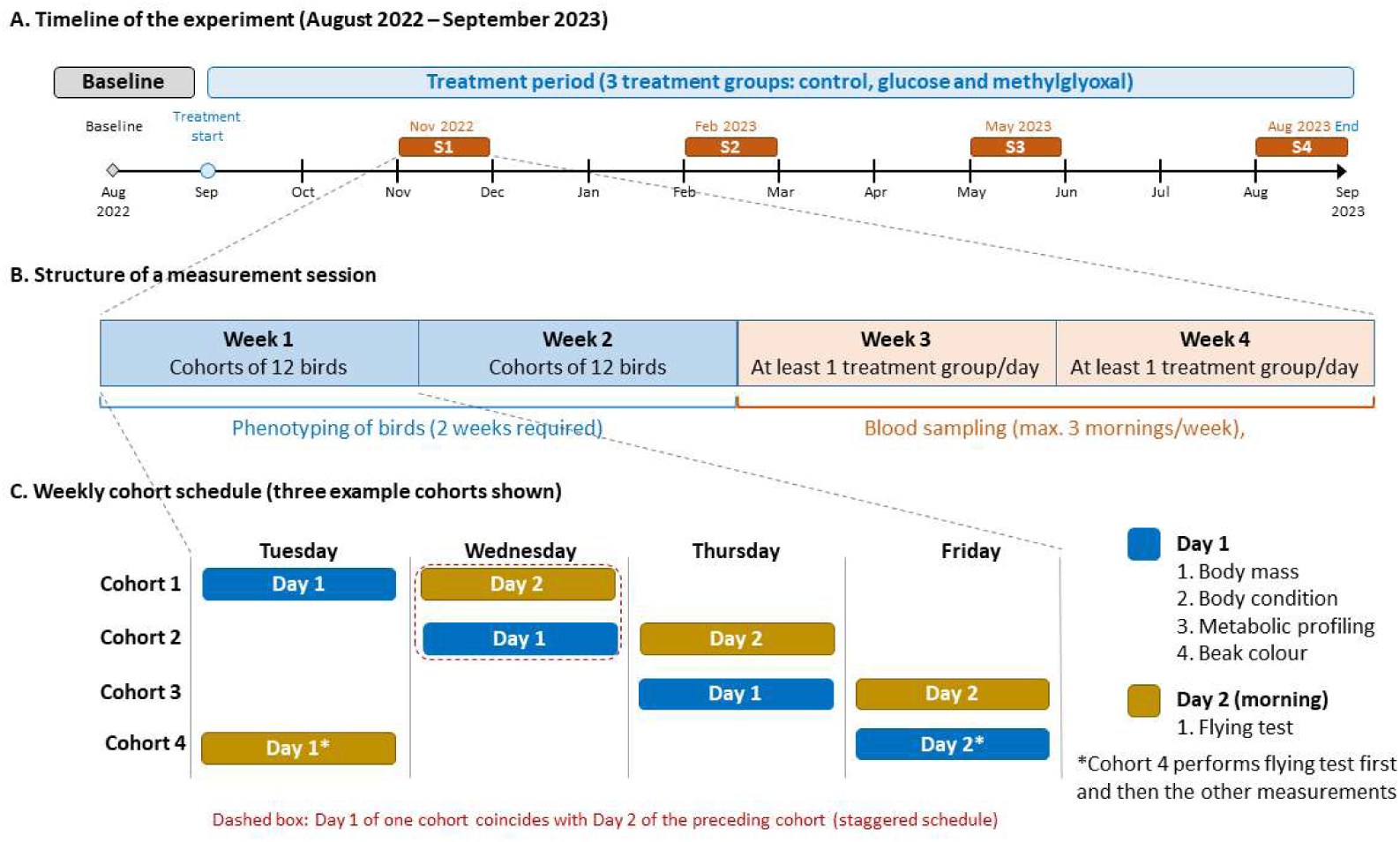
Experimental timeline and measurement schedule. (**A**) Overview of the experiment. Baseline measurements were taken in August 2022, treatment was initiated in September 2022, and the experiment ended in September 2023. Four measurement sessions (S1–S4) were conducted in November 2022, February 2023, May 2023, and August 2023. (**B**) Structure of a measurement session. Each session lasted up to four weeks. During Weeks 1 and 2, birds were assessed in mixed-treatment cohorts of 12, from Tuesday to Friday; two weeks were required to assess all birds. During Weeks 3 and 4, all these birds underwent one blood sampling per week (in the morning) and were then returned to their aviaries, with at least one full treatment group (i.e. control, glucose or methylglyoxal) each day. (**C**) Weekly cohort schedule for Weeks 1 and 2 (e.g. four cohorts). On Day 1, body mass and body condition were measured, followed by metabolic profiling and beak colour assessment. On Day 2 (the following morning), birds underwent a flight test and then they were returned to their aviaries. A new cohort was assessed each day, with the schedule staggered so that Day 1 for one cohort coincided with Day 2 for the preceding cohort (dashed box). One of the cohorts performed the flying test on the first day, and all the other measurements on the last day of the week, filling the schedule. This schedule was reduced when fewer birds remained alive.

At baseline (August 2022) there were 90 birds, and completing all measurements took longer than the usual four weeks for other measurement sessions. During this period, a few individuals died before the entire measurement process was completed. These individuals were replaced by new birds to maintain the initial number of 90 birds aimed for at the start of the experiment. Consequently, some baseline measurements for these replacement birds were performed a few weeks later than the others.

Birds were captured from their aviaries the day before each measurement, and placed in small cages (40.5 cm x 54.5 cm x 29.5 cm), with a maximum of 8 birds per cage. They had ad libitum access to water and food, the latter to be removed on the next morning.They were placed in a room maintained at 20°C with a photoperiod matching the outdoor conditions from the previous day, in which all procedures were carried out. All these procedures were followed (i) to control the birds’ feeding state (postprandial status for respirometry; see below for blood sampling), (ii) to keep environmental conditions consistent across seasons prior to the measurements, (iii) to mitigate handling and thermal stress when entering the temperature cabinet for respirometry (set at 35 °C; see **ESM3**), (iv) for organizational reasons, as the birds could not be captured the same day of the sampling, and (v) to reduce the stress by distancing capture from data collection. Birds underwent procedures in the following order: first respirometry, followed immediately by coloration assessment (both in the afternoon/evening), and the flying test on the following morning. Birds were then released back into the aviaries.

To implement this, birds were selected at random using R **sample()** function, to avoid any capture bias, and 24 birds were captured at the beginning of the week, with a flying test carried out on half of them in the following morning. Flight performance, a proxy for individual maintenance and directly linked to foraging and predator escape capacity (Lindhe Norberg, 2002), was measured using vertical take-off as previously described in Criscuolo et al. (2011) (see further details in **ESM2 - Flying test section**). These twelve birds were then released back to their aviaries, and the other twelve underwent respirometry assessment in the afternoon, divided into three rounds of four birds each, from 14:00 to 18:00 (food was thus removed at 9:00; see **ESM2 - Respirometry** section). Muscle and fat scores were assessed before respirometry, while body mass and cloacal temperature were recorded twice, immediately before and after respirometry. Coloration was measured directly after respirometry for each bird (see **ESM2 - Coloration** section). These twelve birds were then kept overnight under the same indoor conditions for flying test the following morning, except at the end of the week, and another set of twelve birds was caught each day for the next afternoon (see **Figure 2.C**).

For blood sampling, birds were kept indoors from the day before sampling (about 6 birds housed per cage) without access to food since the evening prior to sampling, ensuring an overnight fast. At least one full treatment cage was captured each day, meaning all birds from a single treatment group (control, glucose, or methylglyoxal) were captured together, and birds were released back to their aviaries a few hours after the sampling. When the total number of birds still alive was sufficiently small to allow blood sampling from more than one treatment on the same day, we captured birds from different treatment groups simultaneously. This latter approach both reduced potential effects of day-to-day variation in sampling conditions between groups and helped reducing workload. Details of all measurement procedures, including those used in the pilot study, are provided in **ESM2**, including the potential stress effect of our handling procedure on our blood variables.

Due to limitations in the number of samples that could be processed, albumin glycation rate and plasma glucose levels were measured only in a subsample of birds (n=124 samples). The same applied for AGE measurements, which required larger plasma volumes (n=105 samples; see **ESM2**). Both glyoxal and methylglyoxal molecules had to be derived previous to the determination of their plasma levels by targeted metabolomics (see **ESM2**), and thus all results regarding these metabolites presented in this manuscript are based on concentrations of the derived molecules. Owing to technical issues with the respirometry device, baseline metabolic measurements (August 2022) could not be included in the study.

This study adhered all the legal and ethical regulations and was authorized by the French Ministry of Secondary Education and Research, APAFIS #32475-2021071910541808 v5.

### Statistics

All statistical analyses were performed using R version ≥4.3.3 in RStudio (R Core Team, 2023). For all models, a significance threshold of α ≤ 0.05 was used, while 0.05 < α ≤ 0.1 was interpreted as a trend or marginal effect.

#### Linear Mixed Models (LMMs)

We used linear mixed models (LMMs) to assess the effects of treatment (Control, Glucose, or Methylglyoxal), month of measurement (Baseline, November, February, May, August), (horizontal) age, and sex—including all relevant interactions—on the measured variables (see **Table ESM2.Box 1**). Only horizontal age was included (i.e. age at the beginning of the experiment), as the short duration of the experiment, together with the marked seasonality, potentially made insufficient or even distorted the observation of any longitudinal age-related variations. Models were fitted using the **`lmer()`** function from the **`lme4`** package (Bates et al., 2015). Model selection was performed using the **`dredge()`** function from the **`MuMIn`** package (Burnham and Anderson, 2002), based on log-likelihood (logLik), AICc, ΔAICc, and model weights. Fixed effects such as sex, age, and their interactions (except for the treatment*month interaction) were removed if they did not improve model fit according to information criteria. Sex was retained a priori in models predicting beak coloration due to its biological relevance as a secondary sexual trait. Individual identity was always included as a random effect to account for repeated measures.

#### Covariates and Transformations

To improve model interpretability, we included the following covariates in specific models. Plasma glucose for testing albumin glycation, glyoxal and methylglyoxal levels (log_10_ transformed) when modelling CML and CEL, average body mass (from before and after the respirometry measurement) when predicting RMR, dry mass when predicting fat mass, fresh mass when predicting water content (all these latter four variables log_10_ transformed) and structural size (tarsus and head-beak length in cm) for body mass models. All quantitative predictors were centred to facilitate interpretation of intercepts (Schielzeth, 2010). Response variables were log_10_-transformed when necessary to achieve normality of residuals or to account for allometric relationships (e.g., RMR vs. body mass). We tested the potential effects of body mass in the models for dicarbonyls and AGEs, but it was not retained by the model selection process. A list with the structure of the final models tested for each response variable is provided in **ESM2**.

#### Post-hoc Analyses

Post-hoc comparisons between factor levels were conducted using **`emmeans()`**, and **`emtrends()`** was used to estimate group-specific slopes and their standard errors. Reported results and significance levels in figures are based on comparisons of the same treatments between and within months. Additional models performed but not presented in the main text are described in **ESM2**.

#### Model Diagnostics

Shapiro-Wilks test and Q-Q-plots were used to assess the normality of model residuals. Within-individual repeatability (i.e. repeatability across time points) was estimated using the **`rptGaussian()`** function from the **`rptR`** package (Nakagawa and Schielzeth 2010). Histograms of residuals were also examined when normality tests indicated deviations.

#### Flying test analysis

Since mean and maximum flying speed had a semi-continuous distribution (many zeros for non-flying birds and normally distributed values for flying birds), we used two-part models: (i) a generalized linear model (GLM) with a binomial distribution and logit link to assess the probability of not flying (excluding random effects); (ii) a linear mixed model (LMM) to analyse flying speed among birds that flew, including bird ID as a random effect. Model selection for both parts was performed using **`dredge()`**, as described above.

#### Survival analyses

Kaplan-Meier analyses were performed on the entire population to examine the effect of treatment on survival curves. Additionally, Cox proportional hazards models (using **`coxph()`** from the survival package; Therneau and Grambsch, 2000; Therneau, 2024) were fitted to assess the effects of treatment, sex, and time-varying covariates (e.g., body mass, glucose levels, glycation markers) on survival.

For models with time-varying covariates (i.e. variables measured or recorded at multiple time points; see Zhang et al., 2018), we used values measured immediately before each survival interval (i.e., August and November 2023, February, May and August 2024). Age was calculated as: (i) the initial age at the experiment’ start, and (ii) subsequent ages as initial age + (days elapsed/365). The same type of model (including treatment, sex, body mass and age) was repeated by adding one of the following explanatory variables: whole blood glucose, plasma glucose, albumin glycation, plasma glyoxal, methylglyoxal, CML, CEL, mean and maximum flying speeds (see **ESM2** for a more detailed explanation of the variables). was All quantitative variables were centred to facilitate the interpretation of the parameter estimates (Schielzeth 2010), particularly given the potential confounding effect of significant interaction terms.

A full model including treatment, sex, body mass, age, and their interactions was initially fitted. Model selection was performed using **`dredge()`** to identify the best-supported model. Proportional hazards assumption was checked using Schoenfeld residuals and **`cox.zph()`**. If non-proportionality was detected, time-interaction terms were included.

## Results

### Effects of dietary supplementation on blood glucose, glycated albumin rates and AGE levels

While the supplementation treatments did not affect plasma glucose levels in the subsample of individuals for which albumin glycation was assessed (baseline, February and August 2023; see **ESM2**), they had uneven effects on whole blood glucose levels (**Figure 3.A.; see ESM1** for data on pairwise contrasts between groups). Whole blood glucose levels indeed increased significantly in May for both the control and methylglyoxal groups compared with their respective baseline and February values (see **Table ESM1.1** for de-transformed group means). Subsequently, the control group went back down in August, being significantly lower than in May, while the methylglyoxal group remained higher in August than its baseline. Indeed, in August, both the control and glucose groups showed significantly lower blood glucose levels than the methylglyoxal group. No effects of age were detected. The repeatability of whole blood glucose at the individual level was moderate but significant (R = 0.198; SE = 0.076; CI_95_[0.054, 0.351]; P = 0.0008), whereas plasma glucose levels showed no repeatability (R = 0; SE = 0.067; CI_95_[0, 0.229]; P = 0.5).

**Figure 3.**
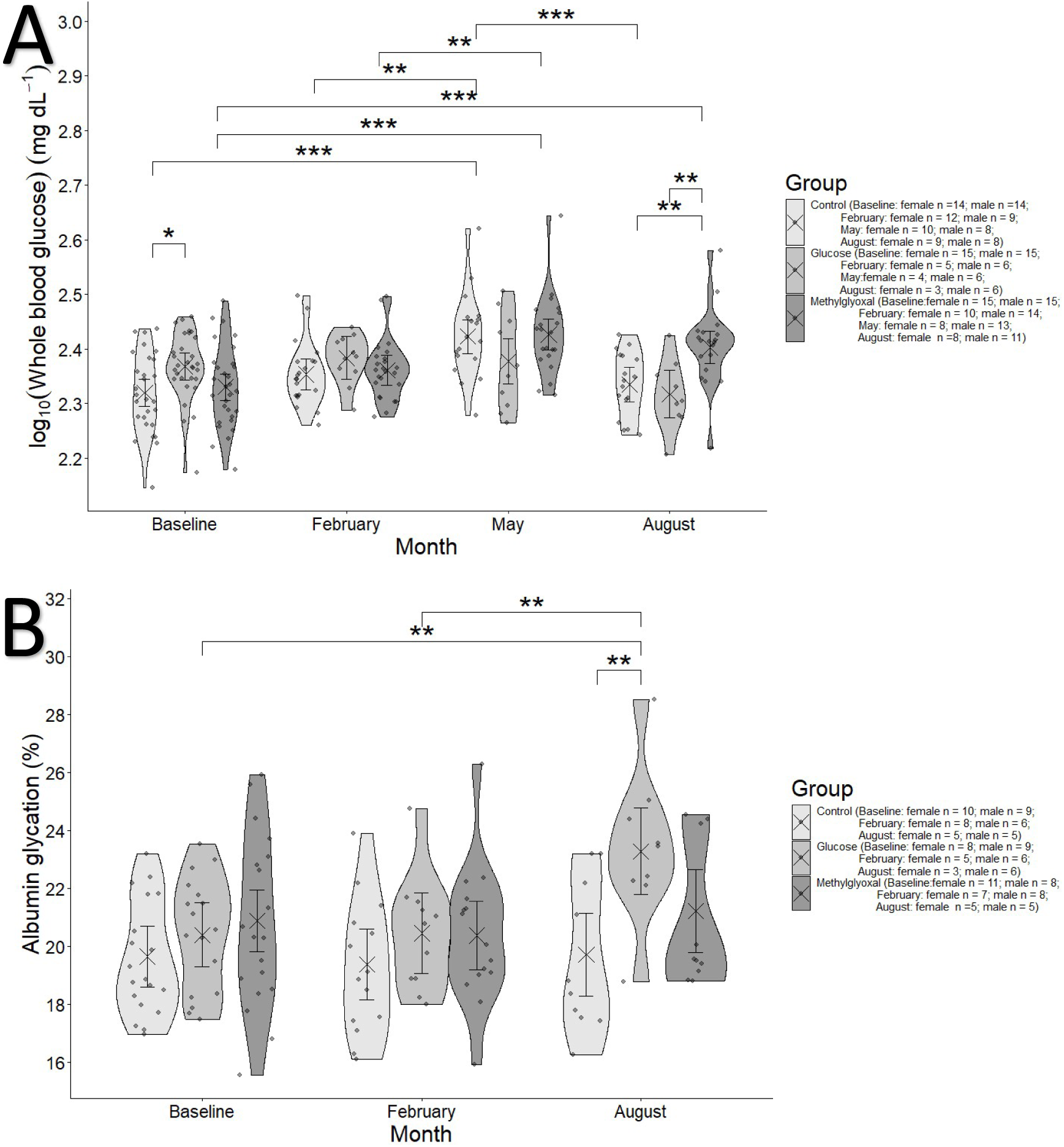
**A.** Whole blood glucose in mg dL^-1^ in the different treatment groups over the course of the experiment. Due to organizational issues, November was missing from the measurements. Crosses and error bars represent model-estimated marginal means ±95% CI. Significance annotations are based on pairwise contrasts performed separately within months and within treatments. **B.** Albumin glycation rate as a percentage of total plasma albumin in the different treatment groups over the course of the experiment. Crosses and error bars represent model-estimated marginal means ±95% CI. Significance annotations are based on pairwise contrasts performed separately within months and within treatments.

A significant effect of glucose supplementation was found on albumin glycation rate (**Figure 3.B**). At the end of the experiment (August 2023), glucose-supplemented birds showed higher rates of glycated albumin than control birds (Glucose - Control: see **Table ESM1.2**; Difference ± SE: 3.567 ± 1.05; t = 3.405, P = 0.0025), as well as higher rates than their own baseline levels (August - Baseline: Difference ± SE: 2.887 ± 0.859; t=3.362; P = 0.003) and February levels (August - February: Difference ± SE: 2.824 ± 0.891; t=3.169; P = 0.006). Glycated albumin rates were moderately repeatable (R = 0.302; SE= 0.124; CI_95_[0.049, 0.534]; P = 0.003).

For plasma methylglyoxal levels (log_10_ transformed), significant differences were detected in specific pairwise comparisons. In August, the control group exhibited significantly lower methylglyoxal levels than at baseline (August - Baseline: Difference ± SE: -0.22 ± 0.048; t = -4.602, P <0.0001; **Figure 4.A**). Additionally, glucose-supplemented birds in February showed lower methylglyoxal levels than at baseline (February - Baseline: Difference ± SE: -0.19 ± 0.076; t = - 2.496, P = 0.037) and the methylglyoxal group on the same month (Glucose - Methylglyoxal: Difference ± SE: -0.2 ± 0.074; t = -2.696; P = 0.022). Glucose-supplemented birds in February also showed lower methylglyoxal levels than at August (February - August: Difference ± SE: -0.125 ± 0.051; t = 2.457, P = 0.042-0.19 ± 0.076; t = -2.496, P = 0.037). Methylglyoxal levels showed no evidence of repeatability (R = 0; SE = 0.071; CI_95_[0, 0.229]; P = 1).

**Figure 4.**
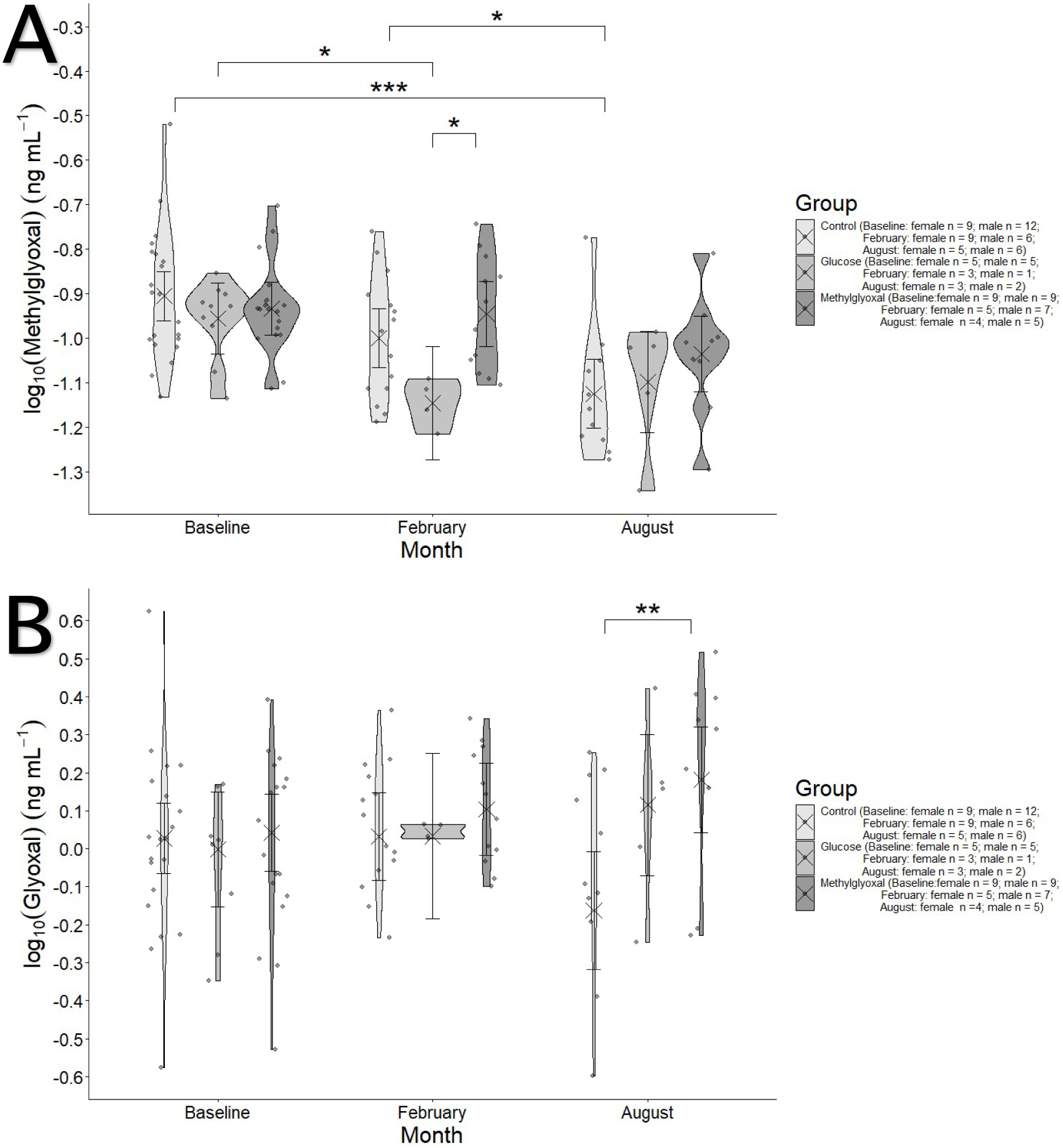
Plasma (**A**) methylglyoxal and (**B**) glyoxal levels in ng mL^-1^ (see **ESM2**) in the different treatment groups over the course of the experiment. Crosses and error bars represent model-estimated marginal means ±95% CI. Significance annotations are based on pairwise contrasts performed separately within months and within treatments.

Glyoxal levels were significantly higher in the methylglyoxal than the control group in August (Difference ± SE: -0.345 ± 0.101; t = -3.402, P = 0.003; **Figure 4.B**). A sex effect on plasma glyoxal was also found (**Figure ESM1.1**), with males showing lower levels in the control than in the methylglyoxal group (estimate ± SE: -0.239 ± 0.071; t = -3.381, P = 0.017). Plasma glyoxal levels (log_10_ transformed) showed a significant effect of age (estimate ± SE: 0.058 ± 0.028, t = 2.086, P = 0.04), although the estimation of the monthly slopes by **`emtrends()`** showed only the August slope to be significant (slope ± SE: 0.099 ± 0.043; CI_95_[0.014, 0.185]). Repeatability was low and non-significant (R = 0.11; SE = 0.112; CI_95_[0, 0.377]; P = 0.28).

Both glucose and methylglyoxal-supplemented birds exhibited significantly higher CML levels in February (Glucose: difference ± SE: -0.016 ± 0.007; t = -2.441, P = 0.043; Methylglyoxal: difference ± SE: 0.015 ± 0.004; t = -3.97, P = 0.0005) and August (Glucose: difference ± SE: 0.028 ± 0.007; t =-4.09, P = 0.0002; Methylglyoxal -0.015 ± 0.005; t = -3.205, P = 0.005) than their baseline (**Figure 5**).The age had a positive effect in the control group, with older birds showing higher CML levels (slope ± SE: 0.006 ± 0.003; CI_95_[0.0009, 0.012]). CML levels repeatability was relatively high (R = 0.527; SE = 0.11; CI_95_[0.269, 0.704]; P = 7.17*10^-6^).

**Figure 5.**
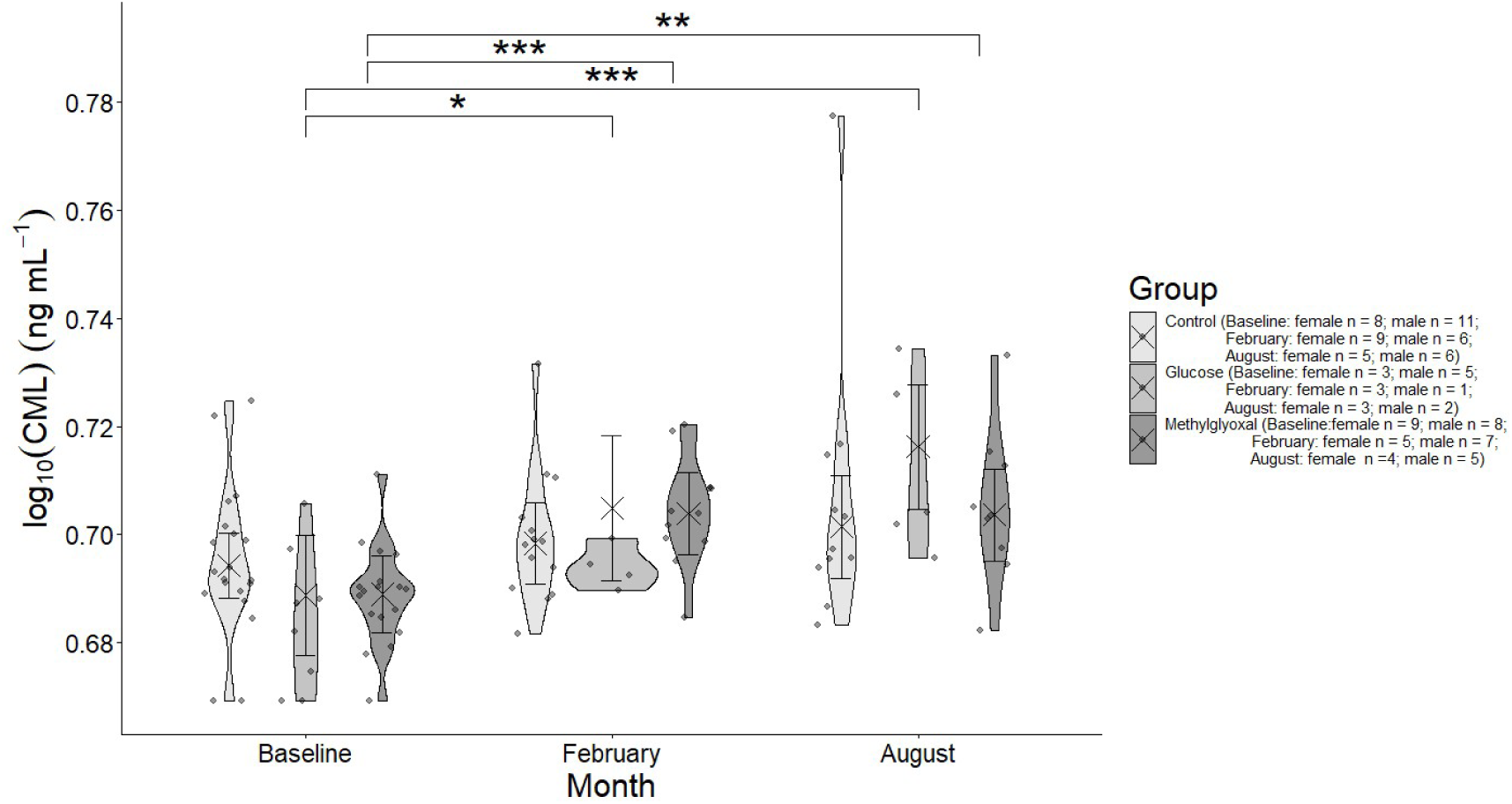
Plasma CML levels ng mL^-1^ in the different treatment groups over the course of the experiment. Crosses and error bars represent model-estimated marginal means ±95% CI. Significance annotations are based on pairwise contrasts performed separately within months and within treatments.

Plasma carboxy-ethyl-lysine (CEL; log10 transformed) levels were significantly modulated only by plasma glyoxal levels (log10), which showed a positive effect (estimate ± SE: 0.051 ± 0.018, t = 2.744, P = 0.007; **Figure 6**). CEL levels showed no significant repeatability (R = 0.14; SE = 0.113; CI95[0, 0.38]; P = 0.179).

**Figure 6.**
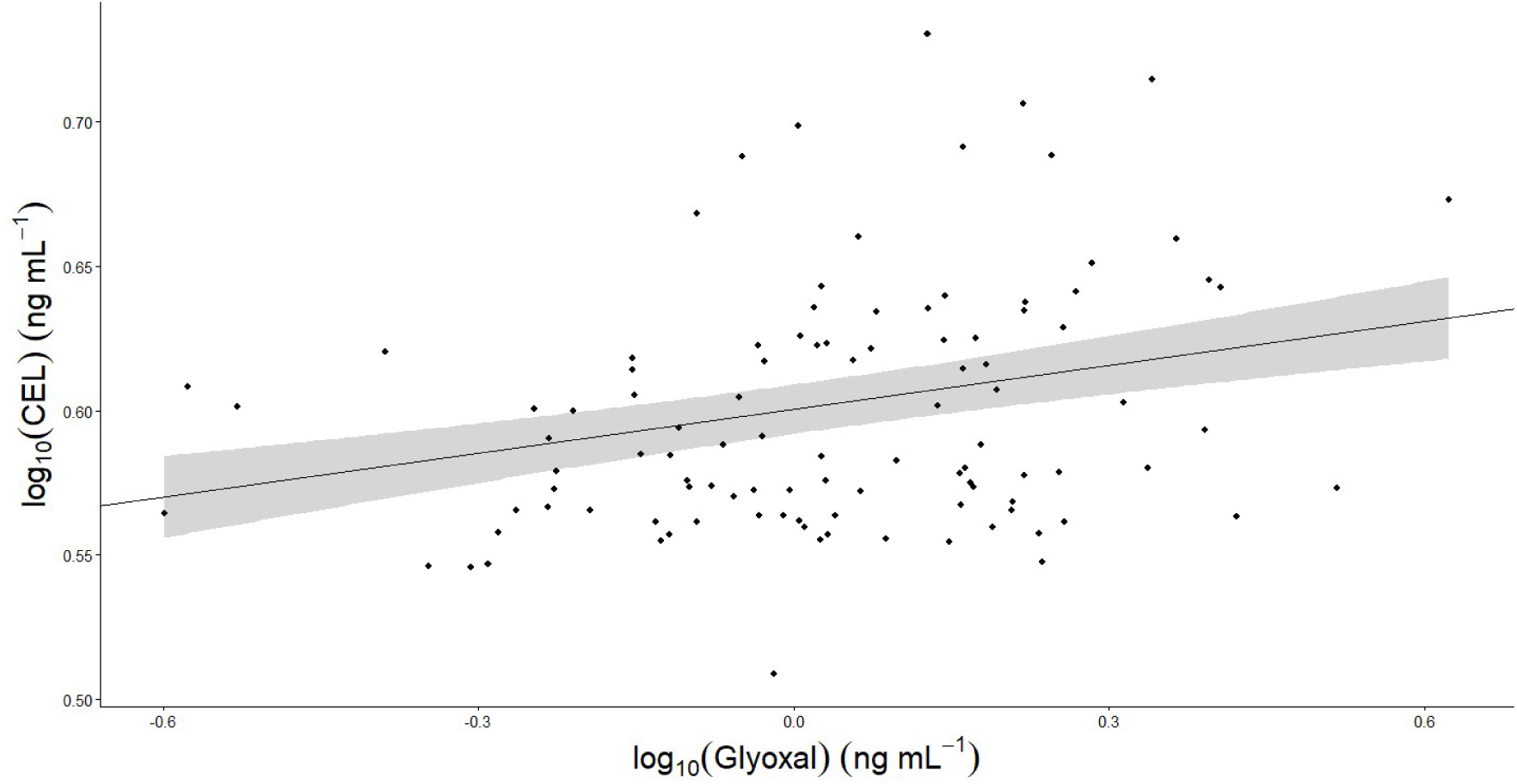
Plasma log_10_(CEL) levels in relation to plasma log_10_(glyoxal) in ng mL^-1^ of glyoxal (see **ESM2**).

### Effects of dietary supplementation on body mass, body condition and body composition

Body mass correlated positively with both tarsus length (slope ± SE: 1.035 g cm^-1^ ± 1.016; t = 2.227, P = 0.028) and head-beak length, the latter showing a stronger association (slope ± SE: 1.086 g cm^-1^ ± 1.013; t = 6.65, P = 5.73*10^-10^, see **Figure ESM1.2.B-C**). Mass was also significantly lower in males (estimate ± SE: -0.954 g ± 1.019; t = -2.579, P = 0.012). No significant treatment or age effects were detected for body mass. Body mass repeatability was very high and significant (R = 0.87; SE = 0.023; CI_95_ = [0.818, 0.906]; P = 4.52*10^-76^).For body condition indices, the only significant difference between groups was a higher level of fat score in the control group in comparison to the methylglyoxal group in August (Control - Methylglyoxal: difference ± SE: 0.96± 0.396; t = 2.424, P = 0.042; Figure **ESM1.3**) and to its own baseline level (August - Baseline: difference ± SE: 0.986 ± 0.023; t = 3.145, P = 0.0156). A negative effect of age was found for fat (slope ± SE: -0.222 ± 0.095; t = -2.347, P = 0.021). Both fat (R = 0.438; SE= 0.063; CI_95_[0.302, 0.551]; P = 1.88*10^-16^) and muscle scores (R = 0.547; SE = 0.054; CI_95_ = [0.434, 0.643]; P = 3.01*10^-22^) were moderately to highly repeatable. Regarding body composition, fat and water content were positively and significantly correlated with the dry and wet masses, respectively of all assessed body compartments (carcass, liver and pectoralis muscle; see **ESM1**). The only significant sex effect was found for fat content in the pectoralis muscle, which was lower in males than in females (estimate ± SE: -0.076 g ± 0.034; t = -2.21, P = 0.033; See **Figure ESM1.4** comparing percentage of fat mass per dry mass between sexes).

### Effects of dietary supplementation on metabolism

The RMR was strongly and positively related to body mass (estimate ± SE: 0.018 ± 1.944*10^-3^; t = 9.126, P = 2.28*10^-16^; **Figure 7.A**). This effect was slightly but significantly steeper in May (estimate ± SE = 5.907*10^-3^ ± 2.227*10^-3^; t = 2.653, P = 0.009) and significantly lower in the glucose-supplemented group (-8.412*10^-3^ ± 2.508*10^-3^; t = -3.355, P = 0.001). Regarding treatment effects (**Figure 7.B**), both groups of treated birds showed a significantly higher RMR in February compared with November (Glucose: difference ± SE: -0.027 ± 0.007; t = -3.917, P = 0.001; Methylglyoxal: difference ± SE: -0.025 ± 0.005; t = -4.774, P < 0.001). Then, the methylglyoxal group decreased its RMR later in May (difference ± SE: 0.015 ± 0.005; t = 2.729, P = 0.035) and August (February - August: difference ± SE: 0.036 ± 0.006; t = 6.229, P < 0.0001; May - August: difference ± SE: 0.021 ± 0.006; t = 3.751, P = 0.001), whereas glucose was significantly decreased only in August (February - August: difference ± SE: 0.026 ± 0.008; t = 3.217, P = 0.008). The RMR was lower in older birds, showing a significant age effect (estimate ± SE: -6.658*10^-3^ ± 2.327*10^-3^; t = -2.862, P = 0.006), that was nevertheless significantly higher (i.e. less negative) and not different from zero in the glucose group (estimate ± SE: 9.597*10^-3^ ± 3.761*10^-3^; t = 2.552, P = 0.012). RMR exhibited moderate repeatability (R = 0.248; SE= 0.079; CI_95_[0.092, 0.409]; P = 0.0003).

**Figure 7.**
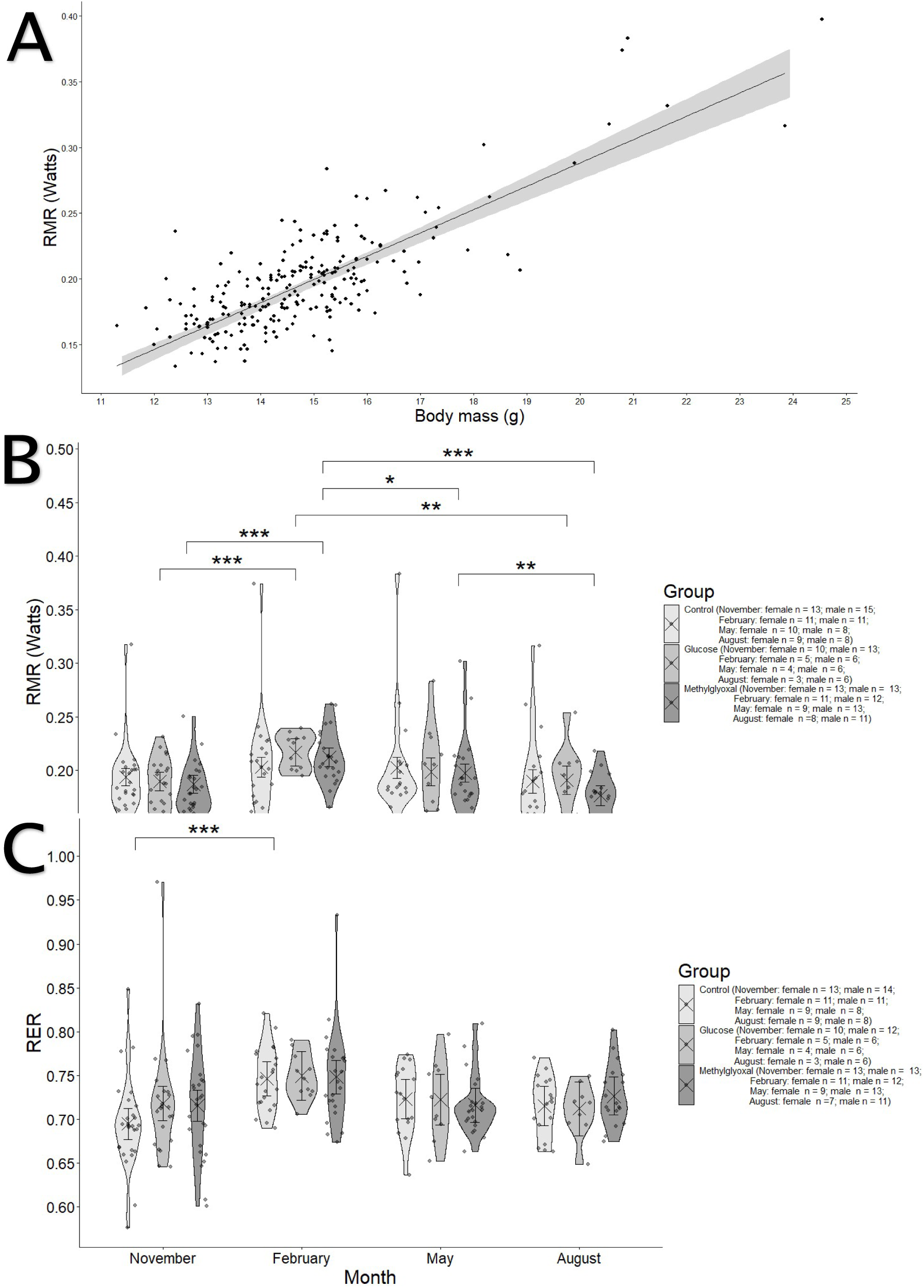
**A** RMR in Watts of birds in the different treatment groups over the course of the experiment and **B** in relation to body mass in grams (slope estimated by the model for the intercept). **C** RER of birds in the different treatment groups over the course of the experiment. Crosses and error bars represent model-estimated marginal means ±95% CI. Significance annotations are based on pairwise contrasts performed separately within months and within treatments.

The RER (**Figure 7.C**) was significantly higher in February for the control groups in comparison to November (difference ± SE: Control: difference ±SE: -0.051 ± 0.013; t = -3.884, P = 0.008).

RMR flexibility was not affected by the treatments, but it decreased with the birds’ age at the beginning of the experiment (**Figure 8**), both in terms of AUC (slope ± SE: -0.018 W year^-1^ ± 0.007; t = -2.693, P = 0.01) and the amplitude of the RMR change (slope ± SE: -0.008 W year^-1^ ± 0.003; t = -2.969, P = 0.005). Body mass correlated significantly and positively with both AUC (slope ± SE: 0.044 W g^-1^ ± 0.008; t = 5.821, P = 9.17*10^-7^) and amplitude (slope ± SE: 0.012 W g^-1^ ± 0.003; t = 3.86, P = 0.0004). Birds with higher initial RMR values showed a higher AUC (slope ± SE: 1.717 W g^-1^ ± 0.384; t = 4.471, P = 6.55*10^-5^) and a lower amplitude (slope ± SE: -0.54 W g ± 0.156; t = -3.458, P = 0.001). Testing the same model on AUC but with age groups (young ≤ 3 years, 4 < medium ≤ 5 and old < 5) showed that both medium (estimate ± SE: -0.043 ± 0.019; t = -2.292, P = 0.027) and old age groups (estimate ± SE: -0.051 ± 0.022; t = -2.376, P = 0.023) had a significant conditional effect, with lower metabolic flexibility than the young group (intercept), although the marginal contrasts between groups (i.e. averaging across sexes) showed only trends to a lower value for both older groups in comparison to the youngest one (Young - Medium ± SE: 0.042 ± 0.019; t = 2.292, P = 0.069; Young - Old ± SE: 0.051 ± 0.022; t = 2.376, P = 0.057). This analysis also showed males having lower values than females (estimate ± SE: 0.034 ± 0.016; t = 2.148, P = 0.038). For variations in RER, no effects were detected except for a significant positive relationship between the AUC and the initial RER value (slope ± SE: 1.285 year^-1^ ± 0.307; t = 4.191, P = 0.0002).

**Figure 8.**
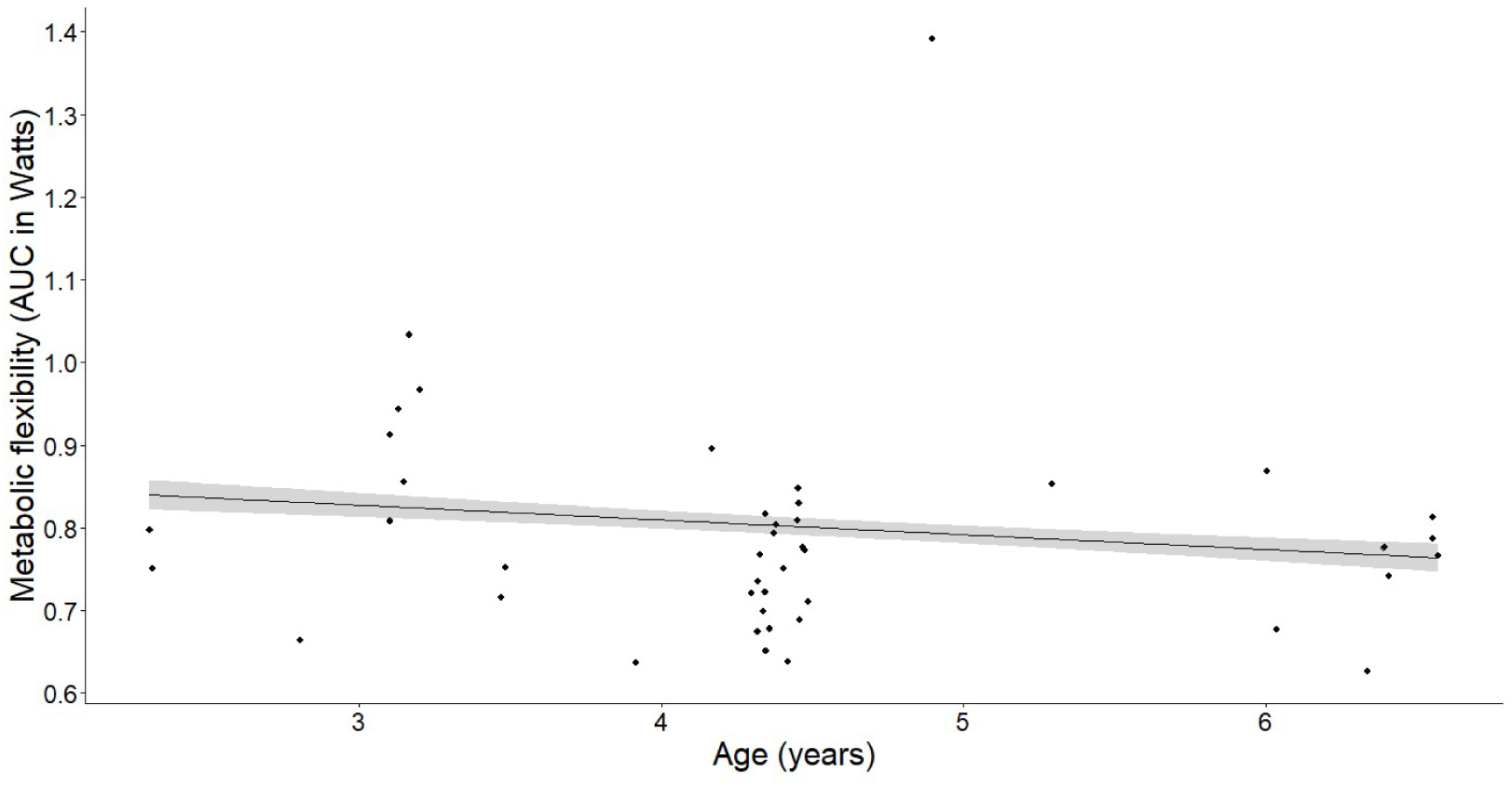
Decrease of RMR flexibility (assessed as AUC in Watts) with age in years.

### Effects of dietary supplementation on flying velocity

In the control group, heavier birds were more likely to fly (slope ± SE: -1.232 ± 0.468; CI_95_[-2.149, -0.314]), whereas the opposite pattern was observed in the methylglyoxal-supplemented group (0.673 ± 0.318; CI_95_[0.049, 1.296]). In the glucose group, body mass had no significant effect. The probability of not flying increased with birds’ age (estimate ± SE: 0.869 ± 0.272; t = 3.196, P = 0.001). There were no differences between birds whatever their treatment in their probability of flying.

For the birds that flew, both the mean (**Figure 9.A**) and maximum (**Figure 9.B**) flying speeds were significantly reduced in the glucose-supplemented group in November compared to both the control (Control - Glucose; Difference ± SE: mean speed: 0.267 m s^-1^ ± 0.071; t = 3.767, P = 0.001; maximum speed: Control - Glucose: 0.318 m s^-1^ ± 0.083; t = 3.852, P = 0.0005) and methylglyoxal groups (Glucose - Methylglyoxal: mean speed: Difference ± SE: -0.172 m s^-1^ ± 0.067, t = -2.63, P = 0.03; maximum speed: Glucose - Methylglyoxal: -0.188 m s^-1^ ± 0.079, t = - 2.375, P = 0.049), and to its May equivalent (mean speed: Difference ± SE: -0.264 m s^-1^ ± 0.071; t = -3.702, P = 0.003; maximum speed: -0.252 m s^-1^ ± 0.088, t = -2.88, P = 0.035). In the baseline, glucose-supplemented birds also showed lower flying speed than the methylglyoxal group (Glucose - Methylglyoxal: mean speed: -0.234 m s^-1^ ± 0.065; t = -3.609, P = 0.001; maximum speed: 0.288 m s^-1^ ± 0.075; t = -3.038, P = 0.008). Finally, methylglyoxal-supplemented birds showed lower mean but not maximum flying speed in November in comparison with the baseline (Baseline - November: 0.143 m s^-1^ ± 0.045, t = 3.162, P = 0.016). Females also exhibited a lower mean speed in November compared to baseline (0.151 m s^-1^ ± 0.039; t = 3.882, P = 0.001).

**Figure 9.**
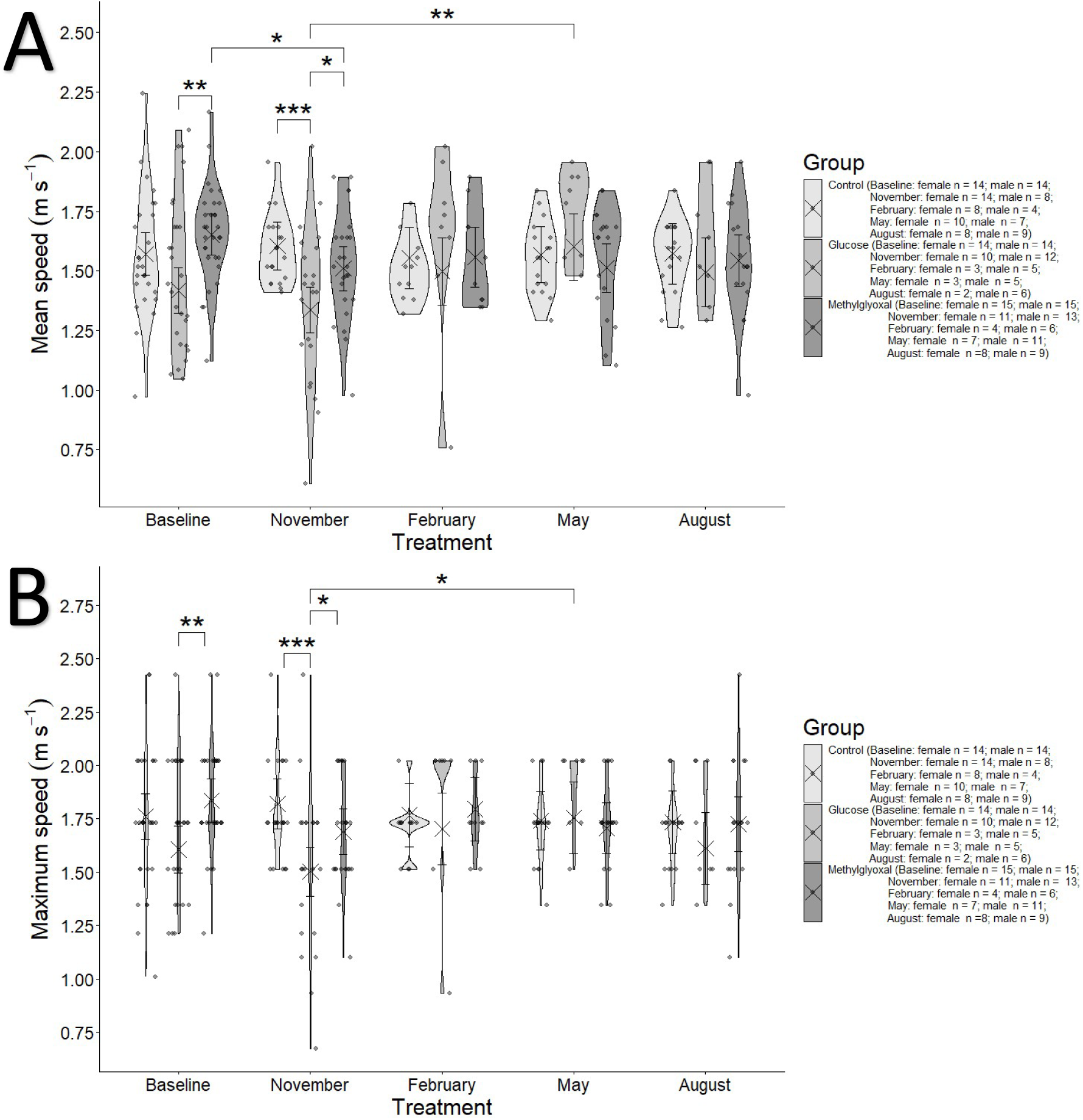
**A** Mean and **B** Maximum flying speed (m s^-1^) in the different treatment groups over the course of the experiment, including only the birds who flew sufficiently. Crosses and error bars represent model-estimated marginal means ±95% CI. Significance annotations are based on pairwise contrasts performed separately within months and within treatments.

The horizontal age component had a significant negative effect on mean speed, i.e. the older the bird the lower its flying velocity (estimate ± SE: -0.049 ± 0.022, t = -2.257, P = 0.027), whereas this effect was only present in the glucose group for the maximum speed (slope ± SE: -0.144 ± 0.041, CI_95_[-0.225, -0.063]). Finally, body mass had a significant negative effect on mean speed (slope ± SE: -0.027 ± 0.013; t = -2.173, P = 0.031). Both mean flight speed (R = 0.536; SE = 0.068; CI_95_[0.39, 0.66]; P = 1.02*10^-15^) and maximum flight speed (R = 0.47; SE = 0.068; CI_95_[0.351, 0.584]; P =7.69*10^-11^) showed substantial repeatability.

### Effects of dietary supplementation on beak coloration

Most of the variance in coloration was located along the side of the tetrahedron formed by the red, green and UV vertices (see **Figure ESM1.6**), but closer on average to the red than green side. Males exhibited significantly redder bills than females (i.e., higher Y values in females; effect:3.462*10^-2^, CI_95_[0.016, 0.053], P = 0.0004). Glucose treatment significantly affected coloration, resulting in a higher Z value (i.e., increased UV; effect = 4.283*10^-2^, CI_95_[0.0005, 0.083], P = 0.0448). Methylglyoxal treatment also led to a higher Z value (effect = 0.04, CI_95_[0.0005, 0.083], P = 0.006) and to a lower X (i.e., a shift away from red towards a “bluer” hue; effect: -0.02, CI_95_[-0.041, -0.001], P = 0.05). Significant deviations from baseline values were observed in all months, with higher Y levels (i.e., a “greener” hue; effects: November = 0.018, CI_95_[0.003, 0.034], P = 0.018; February = 0.038, CI_95_[0.021, 0.055], P < 0.0001; May = 0.035, CI_95_[0.016, 0.053], P = 0.0002; August = 0.061, CI_95_[0.039, 0.08], P < 0.0001) and lower X in May (-0.048, CI_95_[-0.068, -0.027], P < 0.0001). Significant interaction effects were detected for methylglyoxal-supplemented birds in May for the X axis (i.e. an attenuation of the loss in redness observed in May; effect = 0.032, CI_95_[0.008, 0.056], P = 0.011) and in August for the Z axis (i.e. an avoidance in August of the methylglyoxal-induced increase of UV; estimate = -0.055, CI_95_[-0.106, -0.003], P = 0.04). Sex-specific interactions were also significant for the Y value in November, with lower effects in females (estimate = -0.025, CI_95_[-0.04, -0.009], P = 0.002). Euclidean distances between all interaction groups, estimated using MRPP (Multi-Response Permutation Procedures), are reported in **ESM5**.

### Effects of dietary supplementation on survival

Kaplan-Meier survival curves for each bird group are presented in **Figure 10**. In the initial Cox model, performed on the complete population and including only treatment, sex and their interaction, the glucose-supplemented group showed a trend towards significance in its hazard ratio compared with the control group (coefficient ± s. e.: β = 0.876 ± 0.468, P = 0.061), corresponding to a 2.4-fold higher mortality risk (CI_95_[0.96, 6]).

**Figure 10.**
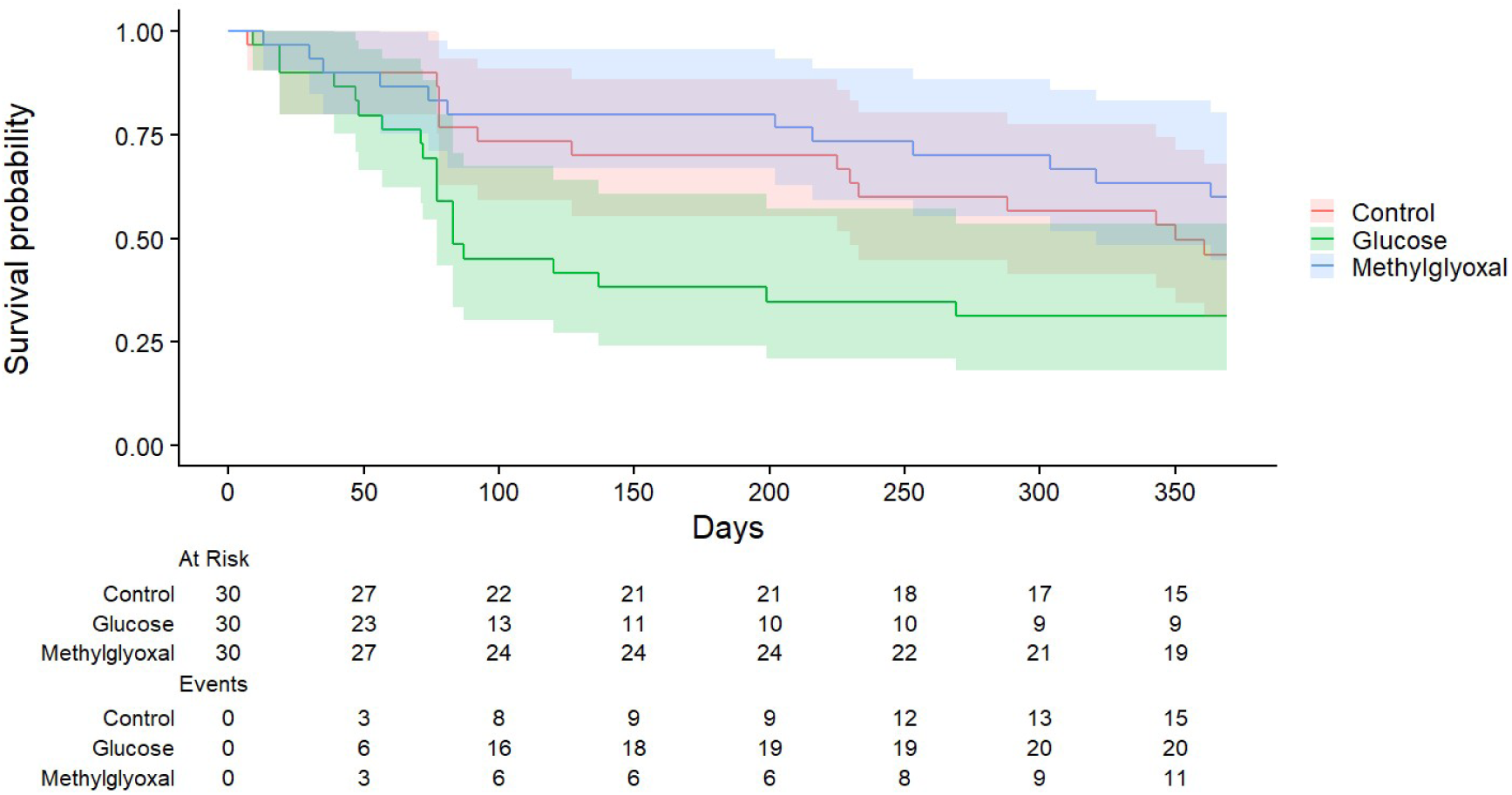
Survival probability across time in days for each experimental treatment group. **At risk** indicates the number of birds still alive at each time, and **Events** the cumulative number of deaths occurred to that time.

After adjusting for age and body mass (resulting in the exclusion of four birds of unknown age: two from the control group and two from the glucose group), and including the interaction between body mass and treatment (selected as the best model by the **`dredge()`** function), the effect of glucose treatment on hazard became significant (β = 1.57 ± 0.506, P = 0.002). The glucose group exhibited a 4.8-fold higher hazard than the control group (P = 0.005), and 3.6-fold higher hazard than the methylglyoxal group (P = 0.023).

Additionally, both body mass and age significantly influenced hazard. Body mass was a protective factor (i.e., associated with lower death hazard) in the control group (β = -1.04 ± 0.273; CI_95_[-1.574, -0.505]) and the methylglyoxal group (β = -0.811 ± 0.331, CI_95_[-1.459, - 0.1634]), but not in the glucose group. In contrast, increasing age was associated with a higher hazard (β = 0.617 ± 0.143, P = 0.00002). No other variable tested had a significant effect on mortality hazard.

## Discussion

### 1. Glucose supplementation increases albumin glycation independently of fasting glycaemia

Our results reveal that glucose supplementation elevated albumin glycation by the end of the experiment (August 2023), despite having no effect on fasting plasma glucose levels. This aligns with previous studies in birds, where dietary carbohydrate supplementation rarely alters blood glucose regulation (Basile et al. 2022). This is potentially modulated by compensatory mechanisms such as reduced food intake (as observed in our pilot study, see **ESM1**). The increase in glycation suggests higher or prolonged postprandial glucose peaks, exposing plasma proteins to glucose for extended periods and promoting glycation (Aragno and Mastrocola 2017). In humans, transient glucose spikes during oral glucose tolerance tests enhance methylglyoxal formation (Zhang et al. 2023), supporting this hypothesis. Indeed, Luo et al. (2025) shows an increase in glucose, fructosamine (an indicator of glycated proteins) and methylglyoxal in streptozotocin-injected chickens, despite a lack of increase in AGEs with the same treatment. In addition, seasonal variations in protein turnover—such as by a cold-induced increase in metabolic demand in February (Bauchinger et al. 2010), may have temporarily masked glycation effects (see **ESM4** for temperature data).

Whole blood glucose levels fluctuated seasonally, with elevated levels in May (persisting in the methylglyoxal group) and a decline in the glucose group by August. These patterns mirror seasonal glucose variations in other outdoor-housed zebra finches, with higher levels during colder, shorter days (Montoya et al. 2018). This is also consistent with our results showing that daily minimum temperature and temperature range were associated with whole blood glucose levels (see **ESM1**). Apart from environmental effects, it is possible that differences in whole blood glucose between groups, particularly the higher levels in the methylglyoxal supplemented birds in August, is linked to haematocrit variations, perhaps due to methylglyoxal increasing cell death (Moreno Borrallo et al. 2026).While plasma glucose levels showed no repeatability, whole blood glucose exhibited moderate repeatability (19.8%), and albumin glycation showed higher repeatability (30.2%), likely reflecting, as expected, medium-term glycaemia more reliably than direct glucose measurements (Kim and Lee 2012).

### 2. Glucose and methylglyoxal supplementation increase plasma CML, but not CEL

The main effects of the supplementations on dicarbonyls and AGEs were an increase in plasma CML levels by both treatments and the prevention of methylglyoxal levels decline over time in the methylglyoxal-supplemented group, where glyoxal levels also increased, particularly in males. The unexpected glyoxal – CEL correlation (despite CEL not being elevated by the treatments), given that CEL is derived from methylglyoxal (Ahmed et al. 1997), may reflect a complex interplay between reactive dicarbonyls. For instance, glyoxal may impact mechanisms like protein turnover (releasing CEL into circulation) or renal filtration (preventing its elimination) differently than methylglyoxal, contributing to reflect its levels more tightly, even if it is not its main precursor.

Szwergold and Miller (2014) proposed that high levels of free amino acids in birds may scavenge reactive carbonyl compounds, potentially reducing their availability for AGE formation. This scavenging effect could explain the lower levels of protein-bound AGEs in birds, although it would not directly affect free AGEs, such as those measured in our study. Notably, AGEs levels in zebra finches were substantially lower than in humans (CML: 4.991 ± 0.184 ng mL^-1^ vs. 173.587-196.051 ng mL^-1^; Wang et al. 2024), suggesting a greater resistance to AGEs accumulation. However, Baker et al. (2022) reported higher AGE-modified albumin in bird plasma compared to mammals (∼40 µg mL^-1^ vs. ∼15 µg mL^-1^), though methodological limitations (i.e., inespecificity of the AGE compounds assessed) complicate any direct comparison.

### 3. Increased mortality under glucose supplementation is independent of glycaemia and AGE accumulation

Only glucose supplementation increased mortality hazard, while methylglyoxal treatment had no effect. This was not driven by elevated fasting glucose, as neither plasma nor whole blood glucose predicted survival. Similarly, neither CEL nor CML, both increased by treatments, affected mortality, suggesting glucose-induced toxicity to operate independently of AGEs accumulation in zebra finches. This contrasts with human studies, where plasma AGEs (including CML and CEL) predict mortality (Sharifi-Zahabi et al. 2021), although cohort differences and other uncontrolled effects complicate direct comparison with birds. However, acute systemic stress (e.g., nutritional imbalance) could have potentially masked the long-term role of AGEs in physiological decline. Alternatively, the observed increased mortality may be due to processes such as: (i) postprandial glucose spikes (rather than a higher sustained fasting glucose) causing oxidative stress, liver damage, or renal strain, (ii) compensatory feeding behaviour (as observed in our pilot study) leading to nutritional deficiencies, and (iii) microbiota alterations, high-sugar diets reducing microbiota diversity in birds (Teyssier et al., 2020; Schmiedová et al. 2022), potentially inducing dysbiosis and inflammation (Do et al., 2018; Zhang et al. 2021; Kawano et al. 2022). Diarrhoea observed at death for some individuals supports this hypothesis. Whole blood glucose, which has been previously shown to negatively impact lifespan in zebra finches (Montoya et al. 2018), was not elevated by glucose supplementation, although it showed higher levels already at the start of the experiment. However, it did not show any direct influence on mortality hazard in our birds. More generally, we recognize that glucose supplementation may have negatively affected the short-term health of the birds through acute physiological or metabolic effects, which represents a limitation of our study. This reflects a broader experimental challenge: increasing circulating glucose levels sufficiently to test their effects without simultaneously inducing nutritional or metabolic stress. Addressing this trade-off will be important in future studies.

Finally, beyond the effects discussed above, mortality hazard also increased with age, consistent with actuarial senescence in captive zebra finches (Briga et al. 2017). Body mass generally acted as a protective factor (i.e. decreased mortality hazard), except in the glucose-supplemented group, contrasting with Montoya et al. (2022), who found no effect of body mass on mortality in captive zebra finches.

### 4. Lack of impact of treatments on body mass, body condition and body composition

Body mass was lower in males and increased in females but was unaffected by dietary supplements, as was body condition. This stability contrasts with glucose-supplemented chickens, which show increased fat deposition (Shapira et al. 1978; Jiang et al. 2008). The lack of treatment effects on body mass or condition, combined with increased mortality in the glucose group, suggests that subtle changes in body composition—such as ectopic fat deposition—may have gone undetected.

### 5. Age reduces seasonally induced metabolic flexibility

As expected, resting metabolic rate (RMR) declined with horizontal age (Moe et al. 2009; Briga 2016; Briga and Verhulst 2021). However, this decline was absent in the glucose group, possibly due to sustained energy intake leading to higher RMR. In addition, seasonal RMR fluctuations (see Swanson 2010; Briga and Verhulst 2021 for zebra finches), with higher values in winter/spring (Versteegh et al., 2012; Beamonte-Barrientos and Verhulst 2013; Stier et al. 2014; Briga and Verhulst 2017; Campbell et al. 2024), were more pronounced in supplemented birds, potentially compensating, again, for a higher caloric intake. The RER also increased modestly in the control group, from 0.69 in November to 0.75 in February, perhaps reflecting a moderate increase in carbohydrate utilization, or a change in the type of lipids metabolized (Price and Mager 2020).

Importantly, the seasonal RMR flexibility (McKechnie 2008; Norin and Metcalfe 2019; Swanson et al. 2022) was lower in older birds, suggesting that they have a reduced capacity to acclimate to environmental temperature changes. This loss in RMR flexibility was produced mainly after 4 years of age, i.e. only at ages not usually observed in natural populations (Zann 1996). The loss of RMR flexibility may contribute to explaining the age-related decline in RMR previously observed in zebra finches (e.g. Moe et al. 2009; Briga 2016; Briga and Verhulst 2021), and represents a promising new marker of physiological ageing in zebra finches. Seemingly, a study in black-capped chickadees (*Poecile atricapillus*) found that summit metabolic rate (maximum cold-induced metabolic rate) strongly predicted winter survival, while BMR did not (Petit et al. 2017). Our findings focused on RMR plasticity (i.e. cold acclimation capacity), combined with these results on acute metabolic responses, suggest that reduced plasticity in metabolic rate could be an important physiological driver of age-related mortality.

### 6. Glucose impairs flying performance and this and beak coloration show senescence

While flying performance did not directly affect mortality hazard, its decline with age reflects a deterioration in health and physiological condition, consistent with ageing patterns in zebra finches (Briga et al. 2017). Moreover, glucose-supplemented birds exhibited lower flying speed than the methylglyoxal group at the baseline and all other groups in November, with no improvement in May. Given the fact that most mortality events happened around November, some process affecting flying performance may also underlie the higher mortality observed in the glucose group (see section 3). Also, heavier birds had a lower mean speed and probability of flying, whereas this pattern was not observed in glucose-supplemented birds. One possible explanation is that the greater energy availability may have helped offset the greater mechanical and energetic constraints associated with flight. Nevertheless, these constraints may be less apparent during startled flight, when birds are more likely to sustain higher speeds despite the associated energetic costs, presumably because rapid flight increases their chances of escaping perceived predators or other threats (Veasey et al. 1998).

Beak coloration, a potential proxy of general health status (Simons et al. 2012), shifted with age (both horizontal and longitudinal) towards less red (mostly more orange), especially in males, which have redder beaks than females. Thus, our results pointing to the existence of senescence in this sexually selected trait (Simons and Verhulst 2011; Fernández-Eslava et al. 2021 for crossbills plumage), contrary to previous results suggesting that decline in beak colour happens only in males the year before their death (Simons et al. 2016).Both methylglyoxal and glucose supplementation increased UV reflectance, while only methylglyoxal reduced redness, producing less saturated beaks (moving towards the centre of the tetrahedron; see **Figure ESM1.5**). Thus, treatment effects, especially methylglyoxal, on beak coloration, may have an impact on bird fitness.

## 7. Conclusions

Chronic glucose supplementation increases mortality in zebra finches, but the underlying mechanism remains unclear. Neither the elevation of protein glycation rate nor the high levels of AGEs explained mortality, suggesting acute glucose dysregulation or microbiota disruption as potential drivers. The lack of mortality effects in the methylglyoxal group further supports AGE-independent pathways for glucose toxicity.

Our findings challenge traditional views of glucose-induced aging and highlight the need for integrative approaches, i.e. combining real-time metabolic profiling, like glucose tolerance test, microbiota analysis, and fine-scale health monitoring, to elucidate glucose toxicity mechanisms. Such studies could advance understanding of avian metabolic adaptations and broader evolutionary trade-offs in aging and longevity.

## Supporting information

ESM5 - Extra results coloration

ESM1 - Extra results

ESM2 - Methods

ESM3 Bird's list

ESM4 - Weather

## Acknowledgements

This research was funded by the Agence Nationale de la Recherche (ANR, project AGEs, ANR21-CE02-0009), and the CNRS/University of Toronto Joint Call 2021. We thank Helène Gachot-Neveu, David Bock, Aurélie Hranitzky and Nicolas Spanier for their work in the animal facility, Alexandre Zahariev for the technical support in respirometry assessments, Fabrice Auge for help in body condition measurements, Pierre Ulrich for 3D printing of pieces used on respirometry and photography set ups and Yaël Metzger for help on part of the baseline data collection. We also thank Max Planck Institute for Biological Intelligence (Pöcking-Seewiesen, Bavaria, Germany) and Bielefeld University (Bielefeld, Germany) for providing part of the birds used in this study. Finally, we would like to thank all the people who contributed with useful discussions about metabolic assessment by indirect calorimetry (Yves Handrich, Roger Colominas Ciuró, Manfred Enstipp, Kenneth Welch Jr and Adrien Levillain). Parts of this paper are reproduced from the PhD thesis of Adrián Moreno Borrallo (Institut Pluridisciplinaire Hubert Curien, CNRS-Strasbourg University, 2024).

## Data availability

All datasets and the R script from this manuscript will be made available on Figshare after publication.

## AI statement

ChatGPT helped providing or improving R code for the stats and graphs (these were always done in R, the AI only provided code). This was always checked by a human before implementing it.

