## Supplementary material for "Decoupling glycation from mortality: glucose, but not methylglyoxal, reduces survival in zebra finches": ESM1 - Extra results

#### 1. Additional information on models shown on the main text

Means and SE of all dependent variables obtained from the models, reported on their original (i.e. de-transformed when required) units, are shown in the following tables (**Tables ESM1.1-8**).

**Table ESM1.1.** Means of whole blood glucose in mg dL$^{-1}$ (de-transformed from the raw log10 outcomes of the model) for each treatment group, with standard errors (SE), degrees of freedom (df) and 0.95 confidence limits (CL).

| Treatment | Month | Mean | SE | df | Lower CL | Upper CL |
|---|---|---|---|---|---|---|
| Control | Baseline | 208,9296131 | 1,029674601 | 231 | 194,98446 | 218,7761624 |
| Glucose | Baseline | 234,4228815 | 1,028726672 | 229 | 218,7761624 | 245,4708916 |
| Methylglyoxal | Baseline | 213,796209 | 1,028726672 | 228 | 204,1737945 | 223,8721139 |
| Control | February | 223,8721139 | 1,034189204 | 239 | 208,9296131 | 239,8832919 |
| Glucose | February | 239,8832919 | 1,047369686 | 248 | 218,7761624 | 263,0267992 |
| Methylglyoxal | February | 229,0867653 | 1,032048246 | 236 | 213,796209 | 245,4708916 |
| Control | May | 263,0267992 | 1,037050726 | 244 | 245,4708916 | 281,8382931 |
| Glucose | May | 239,8832919 | 1,049784123 | 249 | 218,7761624 | 263,0267992 |
| Methylglyoxal | May | 269,1534804 | 1,034189204 | 241 | 251,1886432 | 288,4031503 |
| Control | August | 218,7761624 | 1,038006325 | 246 | 199,5262315 | 234,4228815 |
| Glucose | August | 208,9296131 | 1,052204125 | 250 | 186,2087137 | 229,0867653 |
| Methylglyoxal | August | 251,1886432 | 1,035857465 | 245 | 234,4228815 | 269,1534804 |

**Table ESM1.2.** Means of albumin glycation in % for each treatment group and month, with standard errors (SE), degrees of freedom (df) and 0.95 confidence limits (CL).

| Group | Month | Mean | SE | df | Lower CL | Upper CL |
|---|---|---|---|---|---|---|
| Control | Baseline | 19.6 | 0.534 | 122 | 18.6 | 20.7 |
| Glucose | Baseline | 20.4 | 0.559 | 125 | 19.3 | 21.5 |
| Methylglyoxal | Baseline | 20.9 | 0.536 | 124 | 19.8 | 21.9 |
| Control | February | 19.4 | 0.616 | 130 | 18.1 | 20.6 |
| Glucose | February | 20.5 | 0.698 | 134 | 19.1 | 21.8 |
| Methylglyoxal | February | 20.4 | 0.593 | 129 | 19.2 | 21.5 |
| Control | August | 19.7 | 0.729 | 137 | 18.3 | 21.1 |
| Glucose | August | 23.3 | 0.758 | 136 | 21.8 | 24.8 |
| Methylglyoxal | August | 21.2 | 0.721 | 137 | 19.8 | 22.6 |

**Table ESM1.3.** Plasma glucose group means (mg dL$^{-1}$) for each treatment group and month, with standard errors (SE), degrees of freedom (df) and 0.95 confidence limits (CL).

| Group | Month | Mean | SE | df | Lower CL | Upper CL |
|---|---|---|---|---|---|---|
| Control | Baseline | 367 | 10.9 | 134 | 345 | 388 |
| Glucose | Baseline | 377 | 11.5 | 134 | 354 | 399 |
| Methylglyoxal | Baseline | 364 | 10.9 | 134 | 342 | 385 |
| Control | February | 369 | 12.7 | 134 | 344 | 394 |
| Glucose | February | 389 | 14.4 | 134 | 360 | 417 |
| Methylglyoxal | February | 376 | 12.3 | 134 | 352 | 400 |
| Control | August | 394 | 15.2 | 134 | 364 | 424 |
| Glucose | August | 380 | 15.9 | 134 | 349 | 412 |
| Methylglyoxal | August | 389 | 15.2 | 134 | 359 | 419 |

**Table ESM1.4.** Plasma derived methylglyoxal means (ng $mL^{-1}$) for each treatment group and month, with standard errors (SE), degrees of freedom (df) and 0.95 confidence limits (CL).

| Group | Month | Mean | SE | df | Lower CL | Upper CL |
|---|---|---|---|---|---|---|
| Control | Baseline | 0.124 | 1.066 | 115 | 0.11 | 0.141 |
| Glucose | Baseline | 0.111 | 1.097 | 115 | 0.092 | 0.133 |
| Methylglyoxal | Baseline | 0.117 | 1.072 | 114 | 0.102 | 0.134 |
| Control | February | 0.1 | 1.079 | 115 | 0.086 | 0.116 |
| Glucose | February | 0.071 | 1.16 | 115 | 0.053 | 0.096 |
| Methylglyoxal | February | 0.113 | 1.089 | 115 | 0.096 | 0.134 |
| Control | August | 0.075 | 1.093 | 115 | 0.063 | 0.09 |
| Glucose | August | 0.08 | 1.141 | 115 | 0.061 | 0.104 |
| Methylglyoxal | August | 0.092 | 1.104 | 115 | 0.076 | 0.112 |

**Table ESM1.5.A** Plasma derived glyoxal means (ng $mL^{-1}$) for each treatment group and month, with standard errors (SE), degrees of freedom (df) and 0.95 confidence limits (CL).

| Group | Month | Mean | SE | df | Lower CL | Upper CL |
|---|---|---|---|---|---|---|
| Control | Baseline | 1.063 | 1.117 | 114 | 0.853 | 1.325 |
| Glucose | Baseline | 0.933 | 1.195 | 112 | 0.656 | 1.328 |
| Methylglyoxal | Baseline | 1.025 | 1.124 | 113 | 0.813 | 1.292 |
| Control | February | 1.106 | 1.141 | 115 | 0.851 | 1.437 |
| Glucose | February | 1.044 | 1.286 | 110 | 0.634 | 1.718 |
| Methylglyoxal | February | 1.258 | 1.154 | 115 | 0.948 | 1.671 |
| Control | August | 0.874 | 1.161 | 115 | 0.65 | 1.174 |
| Glucose | August | 1.242 | 1.248 | 115 | 0.801 | 1.925 |
| Methylglyoxal | August | 1.569 | 1.18 | 114 | 0.885 | 2.177 |

**Table ESM1.5.B** Plasma derived glyoxal means (ng $mL^{-1}$) for each treatment group and sex, with standard errors (SE), degrees of freedom (df) and 0.95 confidence limits (CL).

| Group | Sex | Mean | SE | df | Lower CL | Upper CL |
|---|---|---|---|---|---|---|
| Control | female | 1.209 | 1.125 | 48.2 | 0.955 | 1.531 |
| Glucose | female | 1.229 | 1.198 | 35.1 | 0.851 | 1.775 |
| Methylglyoxal | female | 1.097 | 1.14 | 45.1 | 0.842 | 1.428 |
| Control | male | 0.842 | 1.118 | 52.6 | 0.672 | 1.054 |
| Glucose | male | 0.924 | 1.218 | 92.9 | 0.625 | 1.366 |
| Methylglyoxal | male | 1.459 | 1.125 | 48.4 | 1.151 | 1.849 |

**Table ESM1.6.** Plasma derived CML means (ng mL$^{-1}$) for each treatment group and month, with standard errors (SE), degrees of freedom (df) and 0.95 confidence limits (CL).

| Group | Month | Mean | SE | df | Lower CL | Upper CL |
|---|---|---|---|---|---|---|
| Control | Baseline | 4.955 | 1.007 | 81.3 | 4.875 | 5.023 |
| Glucose | Baseline | 4.831 | 1.011 | 83.9 | 4.732 | 4.943 |
| Methylglyoxal | Baseline | 4.898 | 1.008 | 76.4 | 4.819 | 4.977 |
| Control | February | 5 | 1.009 | 97.7 | 4.92 | 5.082 |
| Glucose | February | 5.058 | 1.015 | 112.5 | 4.909 | 5.212 |
| Methylglyoxal | February | 5.07 | 1.009 | 98.3 | 4.989 | 5.164 |
| Control | August | 5.023 | 1.011 | 111.2 | 4.92 | 5.14 |
| Glucose | August | 5.224 | 1.014 | 105.3 | 5.082 | 5.458 |
| Methylglyoxal | August | 5.035 | 1.01 | 107.6 | 4.943 | 5.14 |

**Table ESM1.7.** Plasma derived CEL means (ng mL$^{-1}$) for each treatment group and month, with standard errors (SE), degrees of freedom (df) and 0.95 confidence limits (CL).

| Group | Month | Mean | SE | df | Lower CL | Upper CL |
|---|---|---|---|---|---|---|
| Control | Baseline | 3.999 | 1.021 | 113 | 3.837 | 4.169 |
| Glucose | Baseline | 3.873 | 1.031 | 113 | 3.648 | 4.111 |
| Methylglyoxal | Baseline | 3.917 | 1.023 | 109 | 3.741 | 4.093 |
| Control | February | 3.972 | 1.025 | 115 | 3.776 | 4.169 |
| Glucose | February | 4.217 | 1.049 | 116 | 3.837 | 4.634 |
| Methylglyoxal | February | 4.121 | 1.028 | 115 | 3.899 | 4.355 |
| Control | August | 3.945 | 1.03 | 116 | 3.724 | 4.178 |
| Glucose | August | 4.083 | 1.044 | 116 | 3.75 | 4.446 |
| Methylglyoxal | August | 3.846 | 1.033 | 116 | 3.606 | 4.102 |

**Table ESM1.8.** Means of RMR in Watts for each treatment group and month, with standard errors (SE), degrees of freedom (df) and 0.95 confidence limits (CL).

| Group | Month | Mean | SE | df | Lower CL | Upper CL |
|---|---|---|---|---|---|---|
| Control | November | 0.193 | 0.00410 | 225 | 0.185 | 0.202 |
| Glucose | November | 0.189 | 0.00441 | 216 | 0.180 | 0.197 |
| Methylglyoxal | November | 0.189 | 0.00440 | 224 | 0.180 | 0.198 |
| Control | February | 0.203 | 0.00463 | 238 | 0.194 | 0.213 |
| Glucose | February | 0.212 | 0.00646 | 246 | 0.199 | 0.224 |
| Methylglyoxal | February | 0.212 | 0.00435 | 233 | 0.204 | 0.221 |
| Control | May | 0.202 | 0.00506 | 240 | 0.192 | 0.212 |
| Glucose | May | 0.197 | 0.00675 | 249 | 0.183 | 0.210 |
| Methylglyoxal | May | 0.198 | 0.00440 | 239 | 0.189 | 0.207 |
| Control | August | 0.188 | 0.00570 | 246 | 0.176 | 0.199 |
| Glucose | August | 0.190 | 0.00699 | 250 | 0.177 | 0.204 |
| Methylglyoxal | August | 0.176 | 0.00473 | 243 | 0.167 | 0.186 |

The results of the selected final models exploring the effects of the treatments across the experiment months on the assessed variables not shown in the main text are presented in the following tables (**Table ESM1.9-16**). **Figures ESM1.1-6** show some of the findings from these models whose graphical representations are not included in the main text.

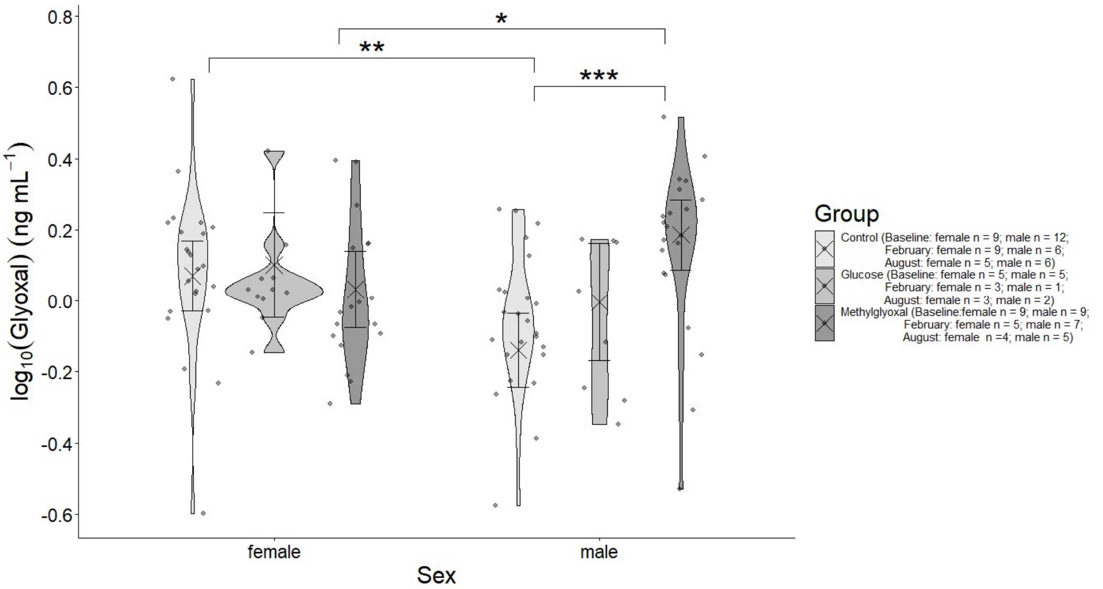


**Figure ESM1.1.** Plasma glyoxal levels in ng $mL^{-1}$ (see **ESM2**) in the different treatment groups and across sexes. Crosses and error bars represent model-estimated marginal means ±95% CI. Significance annotations are based on pairwise contrasts performed separately within months and within treatments.

**Table ESM1.9.** Main model outcomes for plasma glucose (mg $dL^{-1}$).

| | Estimate | StandardError | df | t value | P-value |
|---|---|---|---|---|---|
| **Intercept** | **366.526** | **10.487** | **124** | **34.949** | **$<2*10^{-16}$** |
| Group Glucose | 10.003 | 15.261 | 124 | 0.655 | 0.513 |
| Group Methylglyoxal | -2.947 | 14.831 | 124 | -0.199 | 0.843 |
| Month February | 2.759 | 16.101 | 124 | 0.171 | 0.864 |
| Month August | 27.174 | 17.859 | 124 | 1.522 | 0.131 |
| Group Glucose:Month February | 9.438 | 23.920 | 124 | 0.395 | 0.694 |
| Group Methylglyoxal:Month February | 9.728 | 22.551 | 124 | 0.431 | 0.667 |
| Group Glucose:Month August | -23.370 | 25.963 | 124 | -0.900 | 0.370 |
| Group Methylglyoxal:Month August | -2.053 | 25.257 | 124 | -0.081 | 0.935 |

<u>Shapiro-Wilk normality test</u>

W = 0.98828, P-value = 0.3696

**Table ESM1.10.** Main model outcomes for body mass (g).

| | Estimate | Standard Error | df | t value | P-value |
|---|---|---|---|---|---|
| **Intercept** | **14.77** | **0.2647** | **161.8** | **55.818** | **$< 2*10^{-16}$** |
| Group Glucose | 0.3233 | 0.3384 | 153.3 | 0.955 | 0.34094 |
| Group Methylglyoxal | 0.1301 | 0.2560 | 318.2 | 0.508 | 0.61156 |
| *Month November* | *0.3459* | *0.1912* | *237.8* | *1.810* | *0.07163* |
| Month February | 0.3245 | 0.2126 | 240.4 | 1.527 | 0.12817 |
| Month May | 0.2284 | 0.2163 | 240.5 | 1.056 | 0.29213 |
| **Month August** | **0.5703** | **0.2288** | **240.6** | **2.492** | **0.01337** |
| **Sex male** | **-0.7306** | **0.2776** | **8753** | **-2.632** | **0.01003** |
| **C_Tarsus** | **0.7040** | **0.2330** | **8811** | **3.022** | **0.00329** |
| **C_Head_beak** | **1.304** | **0.1879** | **9049** | **6.940** | **$5.73*10^{-10}$** |
| *Group Glucose:Month November* | *-0.5280* | *0.2776* | *239.9* | *-1.902* | *0.05836* |
| Group Methylglyoxal:Month November | -0.4231 | 0.2738 | 238.6 | -1.545 | 0.12363 |
| Group Glucose:Month February | -0.4482 | 0.3438 | 244.4 | -1.304 | 0.19355 |
| Group Methylglyoxal:Month February | $4.541*10^{-2}$ | 0.2935 | 241.1 | 0.155 | 0.87715 |
| Group Glucose:Month May | -0.4684 | 0.3542 | 244.2 | -1.322 | 0.18736 |
| Group Methylglyoxal:Month May | $6.427*10^{-4}$ | 0.3005 | 241.2 | 0.002 | 0.99830 |
| *Group Glucose:Month August* | *-0.6342* | *0.3710* | *243.7* | *-1.710* | *0.08862* |
| *Group Methylglyoxal:Month August* | *-0.5525* | *0.3146* | *241.2* | *-1.756* | *0.08034* |

<u>Shapiro-Wilk normality test</u>

W = 0.96665, P-value = $9.392*10^{-7}$

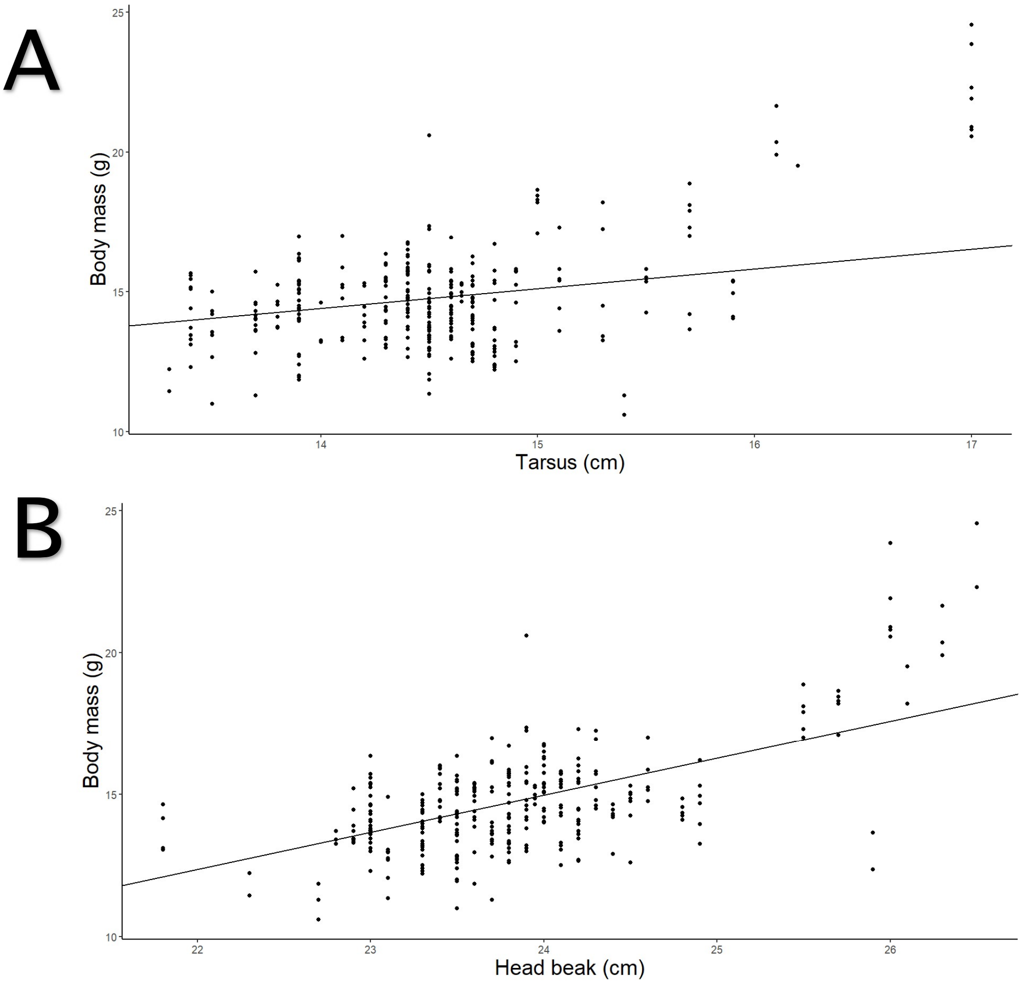


**Figure ESM1.2.** Variation in body mass in grams in function of **A** tarsus length and **B** head-beak length, both in centimetres.

**Table ESM1.11.** Main model outcomes for fat score.

| | Estimate | Standard Error | df | t value | P-value |
|---|---|---|---|---|---|
| **Intercept** | 2.02541 | 0.23052 | 193.89058 | 8.786 | $8.2*10^{-16}$ |
| **Group Glucose** | 0.33245 | 0.32956 | 190.29561 | 1.009 | 0.314380 |
| **Group Methylglyoxal** | 0.44789 | 0.32336 | 190.96566 | 1.385 | 0.167633 |
| **Month November** | 0.56741 | 0.25726 | 242.56322 | 2.206 | 0.028352 |
| **Month February** | 0.32507 | 0.27887 | 255.82964 | 1.166 | 0.244837 |
| **Month May** | 0.38112 | 0.28367 | 270.29897 | 1.344 | 0.180228 |
| **Month August** | 0.98610 | 0.30518 | 284.16248 | 3.231 | 0.001378 |
| **C_Age_years** | -0.22248 | 0.09478 | 82.19807 | -2.347 | 0.021319 |
| **Group Glucose:Month November** | -0.54758 | 0.36698 | 240.60418 | -1.492 | 0.136971 |
| **Group Methylglyoxal:Month November** | -0.61629 | 0.35475 | 237.15124 | -1.737 | 0.083642 |
| **Group Glucose:Month February** | -0.63914 | 0.44394 | 254.02213 | -1.440 | 0.151182 |

| | | | | | |
|---|---|---|---|---|---|
| **Group Methylglyoxal:Month February** | -0.39318 | 0.37571 | 241.86937 | -1.047 | 0.296373 |
| **Group Glucose:Month May** | -0.40966 | 0.45446 | 254.40199 | -0.901 | 0.368210 |
| **Group Methylglyoxal:Month May** | -0.21487 | 0.38093 | 242.41543 | -0.564 | 0.573231 |
| **Group Glucose:Month August** | -1.29942 | 0.47617 | 253.59405 | -2.729 | 0.006799 |
| **Group Methylglyoxal:Month August** | -1.40804 | 0.39811 | 243.20748 | -3.537 | 0.000485 |

Shapiro-Wilk normality test

W = 0.99727, p-value = 0.8801

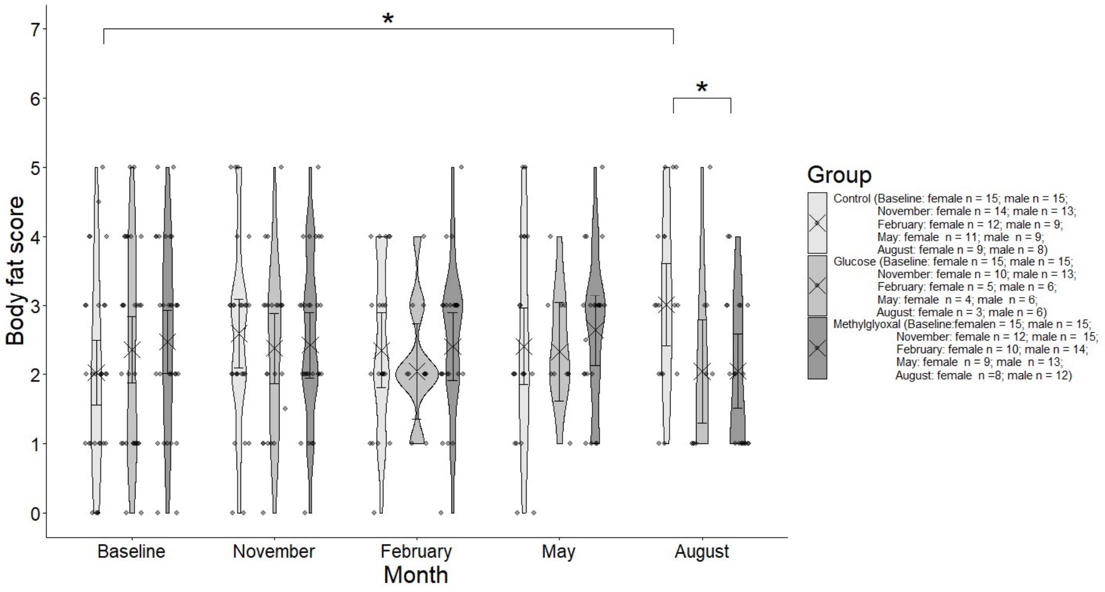


**Figure ESM1.3.** Body fat score in the different treatment groups over the course of the experiment. Points and error bars represent model-estimated marginal means ±95% CI. Significance annotations are based on pairwise contrasts performed separately within months and within treatments.

**Table ESM1.12.** Model on muscle score.

| | Estimate | Standard Error | df | t value | P-value |
|---|---|---|---|---|---|
| **Intercept** | 2.337007 | 0.117578 | 140.592082 | 19.876 | $<2*10^{-16}$ |
| **Group Glucose** | 0.050000 | 0.149495 | 161.243310 | 0.334 | 0.738 |
| **Group Methylglyoxal** | 0.116667 | 0.149495 | 161.243310 | 0.780 | 0.436 |
| **Month November** | 0.046778 | 0.105230 | 234.166254 | 0.445 | 0.657 |
| **Month February** | 0.086232 | 0.114595 | 239.164800 | 0.752 | 0.452 |
| **Month May** | 0.104367 | 0.116560 | 239.716807 | 0.895 | 0.371 |
| **Month August** | -0.127189 | 0.123301 | 240.223083 | -1.032 | 0.303 |
| **Sex male** | -0.107347 | 0.102958 | 81.647502 | -1.043 | 0.300 |
| **Group Glucose:Month November** | -0.157669 | 0.153524 | 239.386719 | -1.027 | 0.305 |
| **Group Methylglyoxal:Month November** | 0.162970 | 0.148511 | 233.527511 | 1.097 | 0.274 |
| **Group Glucose:Month February** | 0.141425 | 0.188052 | 250.146041 | 0.752 | 0.453 |
| **Group Methylglyoxal:Month February** | 0.087124 | 0.158368 | 237.470593 | 0.550 | 0.583 |
| **Group Glucose:Month May** | -0.020218 | 0.193816 | 250.249267 | -0.104 | 0.917 |

| | | | | | |
|---|---|---|---|---|---|
| **Group Methylglyoxal:Month May** | 0.079624 | 0.162041 | 238.238481 | 0.491 | 0.624 |
| **Group Glucose:Month August** | 0.003206 | 0.203039 | 249.579948 | 0.016 | 0.987 |
| **Group Methylglyoxal:Month August** | 0.210935 | 0.169425 | 238.758297 | 1.245 | 0.214 |

<u>Shapiro-Wilk normality test</u>

W = 0.98482, p-value = 0.001834

**Table ESM1.13.A.1.** Main model outcomes for body composition: carcass fat.

| | **Estimate** | **Standard Error** | **t value** | **P-value** |
|---|---|---|---|---|
| **Intercept** | **-0.474844** | **0.041945** | **-11.321** | **$9.74*10^{-14}$** |
| **C_logDry_mass** | **2.235854** | **0.221611** | **10.089** | **$2.67*10^{-12}$** |
| Group Glucose | 0.034724 | 0.046355 | 0.749 | 0.458 |
| Group Methylglyoxal | 0.007832 | 0.057608 | 0.136 | 0.893 |
| Sex male | -0.021303 | 0.044012 | -0.484 | 0.631 |

Residual standard error: 0.1347 on 38 degrees of freedom

Multiple $R^2$: 0.7594, Adjusted $R^2$: 0.7341

F-statistic: 29.99 on 4 and 38 DF, P-value: $2.701*10^{-11}$

<u>Shapiro-Wilk normality test</u>

W = 0.94737, P-value = 0.0478

**Table ESM1.13.A.2.** Main model outcomes for body composition: carcass water.

| | **Estimate** | **Standard Error** | **t value** | **P-value** |
|---|---|---|---|---|
| **Intercept** | **0.734821** | **0.006877** | **106.855** | **$<2*10^{-16}$** |
| **C_logFresh_mass** | **0.974862** | **0.047286** | **20.616** | **$<2*10^{-16}$** |
| *Sex male* | *0.013198* | *0.007184* | *1.837* | *0.0742* |
| Group Glucose | 0.008910 | 0.007805 | 1.141 | 0.2610 |
| Group Methylglyoxal | 0.006292 | 0.009549 | 0.659 | 0.5141 |
| C_logFresh_mass:Sex male | 0.147045 | 0.091405 | 1.609 | 0.1162 |

Residual standard error: 0.02247 on 37 degrees of freedom

Multiple $R^2$: 0.9475, Adjusted $R^2$: 0.9404

F-statistic: 133.5 on 5 and 37 DF, P-value: $< 2.2*10^{-16}$

<u>Shapiro-Wilk normality test</u>

W = 0.96823, P-value = 0.2744

**Table ESM1.13.B.1.** Main model outcomes for body composition: liver fat.

| | Estimate | Standard Error | t value | P-value |
|---|---|---|---|---|
| **Intercept** | **-1.57189** | **0.02824** | **-55.670** | **< $2*10^{-16}$** |
| **C_logDry_mass** | **0.51579** | **0.08957** | **5.758** | **$1.33*10^{-6}$** |
| Group Glucose | 0.02540 | 0.03230 | 0.786 | 0.437 |
| Group Methylglyoxal | 0.01651 | 0.03896 | 0.424 | 0.674 |
| Sex male | -0.03125 | 0.02972 | -1.052 | 0.300 |

Residual standard error: 0.09211 on 37 degrees of freedom

Multiple $R^2$: 0.5304, Adjusted $R^2$: 0.4796

F-statistic: 10.45 on 4 and 37 DF, P-value: $9.146*10^{-6}$

<u>Shapiro-Wilk normality test</u>

W = 0.96202, P-value = 0.1743

**Table ESM1.13.B.2.** Main model outcomes for body composition: liver water.

| | Estimate | Standard Error | t value | P-value |
|---|---|---|---|---|
| **Intercept** | **-0.585973** | **0.003656** | **-160.264** | **<$2*10^{-16}$** |
| **C_logFresh_mass** | **0.993673** | **0.014416** | **68.926** | **<$2*10^{-16}$** |
| Group Glucose | -0.008036 | 0.005097 | -1.577 | 0.123 |
| Group Methylglyoxal | -0.004484 | 0.006092 | -0.736 | 0.466 |

Residual standard error: 0.01461 on 38 degrees of freedom

Multiple $R^2$: 0.9921, Adjusted $R^2$: 0.9915

F-statistic: 1586 on 3 and 38 DF, P-value: < $2.2*10^{-16}$

<u>Shapiro-Wilk normality test</u>

W = 0.97786, P-value = 0.5799

**Table ESM1.13.C.1.** Main model outcomes for body composition: muscle (pectorals) fat.

| | Estimate | Standard Error | t value | P-value |
|---|---|---|---|---|
| **Intercept** | **-1.29958** | **0.03437** | **-37.809** | **< $2*10^{-16}$** |
| **C_logDry_mass** | **0.93117** | **0.11247** | **8.279** | **$4.92*10^{-10}$** |
| Group Glucose | 0.01706 | 0.03924 | 0.435 | 0.6663 |
| Group Methylglyoxal | -0.03274 | 0.04652 | -0.704 | 0.4858 |
| **Sex male** | **-0.07614** | **0.03445** | **-2.210** | **0.0332** |

Residual standard error: 0.1112 on 38 degrees of freedom

Multiple $R^2$: 0.6725, Adjusted $R^2$: 0.6381

F-statistic: 19.51 on 4 and 38 DF, P-value: $8.464*10^{-9}$

<u>Shapiro-Wilk normality test</u>

W = 0.87466, P-value = 0.0002322

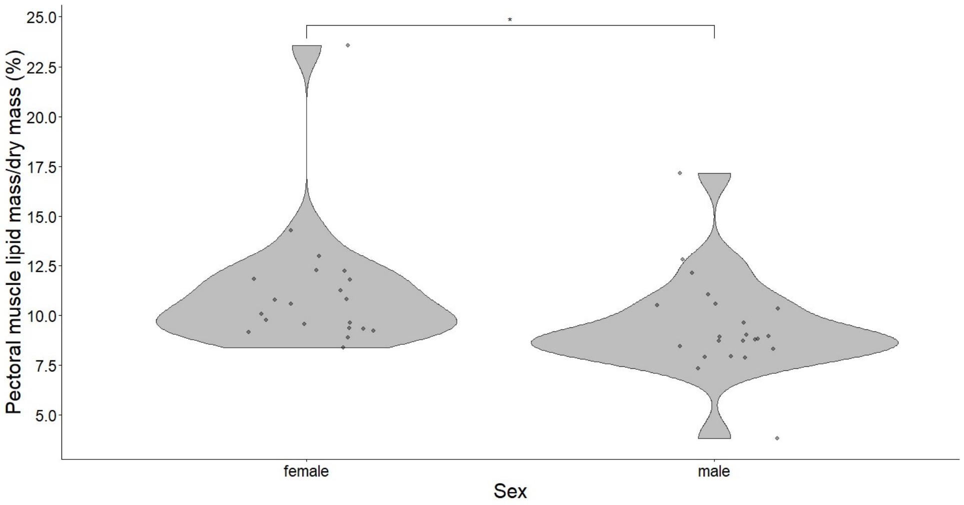


**Figure ESM1.4.** Pectoral muscle lipid mass in birds dead during the experiment as a percentage of total dry mass across sexes. Significance annotation is based on a marginal contrast performed on absolute lipid mass from the model adjusting by (centred) total dry pectoral mass. Mass proportion is nevertheless represented for an improved visualization.

**Table ESM1.13.C.2.** Main model outcomes for body composition: muscle (pectorals) water.

| | Estimate | Standard Error | t value | P-value |
|---|---|---|---|---|
| **Intercept** | **0.122431** | **0.003093** | **39.584** | **$<2*10^{-16}$** |
| **C_logFresh_mass** | **1.011731** | **0.012237** | **82.676** | **$<2*10^{-16}$** |
| Group Glucose | -0.003535 | 0.004305 | -0.821 | 0.417 |
| Group Methylglyoxal | -0.002117 | 0.005123 | -0.413 | 0.682 |

Residual standard error: 0.0123 on 39 degrees of freedom

Multiple $R^2$: 0.9946, Adjusted $R^2$: 0.9942

F-statistic: 2380 on 3 and 39 DF, P-value: $< 2.2*10^{-16}$

<u>Shapiro-Wilk normality test</u>

W = 0.92583, p-value = 0.008387

**Table ESM1.14.A.** Main model outcomes for RMR.

| | Estimate | Standard E rror | df | t value | P-value |
|---|---|---|---|---|---|
| **Intercept** | **$1.939*10^{-1}$** | **$3.849*10^{-3}$** | **$1.969*10^{2}$** | **50.389** | **$< 2*10^{-16}$** |

| | | | | | |
|---|---|---|---|---|---|
| Group Glucose | -3.972*$10^{-3}$ | 5.666*$10^{-3}$ | 1.948*$10^{2}$ | -0.701 | 0.48412 |
| Group Methylglyoxal | -6.671*$10^{-3}$ | 5.584*$10^{-3}$ | 2.037*$10^{2}$ | -1.195 | 0.23359 |
| *Month February* | *9.185*$10^{-3}$* | *5.130*$10^{-3}$* | *1.752*$10^{2}$* | *1.791* | *0.07508* |
| Month May | 8.763*$10^{-3}$ | 5.408*$10^{-3}$ | 1.864*$10^{2}$ | 1.621 | 0.10681 |
| Month August | -3.943*$10^{-3}$ | 5.749*$10^{-3}$ | 2.001*$10^{2}$ | -0.686 | 0.49354 |
| **C_Age_years** | **-6.658*$10^{-3}$** | **2.327*$10^{-3}$** | **6.698*10** | **-2.862** | **0.00562** |
| **C_BM** | **1.775*$10^{-2}$** | **1.944*$10^{-3}$** | **1.666*$10^{2}$** | **9.126** | **2.28*$10^{-16}$** |
| **Group Glucose:Month February** | **1.793*$10^{-2}$** | **8.330*$10^{-3}$** | **1.828*$10^{2}$** | **2.152** | **0.03270** |
| **Group Methylglyoxal:Month February** | **1.594*$10^{-2}$** | **7.207*$10^{-3}$** | **1.744*$10^{2}$** | **2.212** | **0.02827** |
| Group Glucose:Month May | 9.238*$10^{-5}$ | 8.773*$10^{-3}$ | 1.916*$10^{2}$ | 0.011 | 0.99161 |
| Group Methylglyoxal:Month May | 1.646*$10^{-3}$ | 7.537*$10^{-3}$ | 1.873*$10^{2}$ | 0.218 | 0.82735 |
| Group Glucose:Month August | 4.893*$10^{-3}$ | 9.101*$10^{-3}$ | 1.981*$10^{2}$ | 0.538 | 0.59144 |
| Group Methylglyoxal:Month August | -6.711*$10^{-3}$ | 8.030*$10^{-3}$ | 2.003*$10^{2}$ | -0.836 | 0.40434 |
| **Group Glucose:C_Age_years** | **9.597*$10^{-3}$** | **3.761*$10^{-3}$** | **8.909*10** | **2.552** | **0.01242** |
| Group Methylglyoxal:C_Age_years | -6.247*$10^{-4}$ | 3.345*$10^{-3}$ | 8.042*10 | -0.187 | 0.85234 |
| **Group Glucose:C_BM** | **-8.412*$10^{-3}$** | **2.508*$10^{-3}$** | **1.165*$10^{2}$** | **-3.355** | **0.00107** |
| Group Methylglyoxal:C_BM | -3.361*$10^{-3}$ | 2.462*$10^{-3}$ | 8.638*10 | -1.365 | 0.17581 |
| Month February:C_BM | 2.874*$10^{-3}$ | 2.254*$10^{-3}$ | 1.800*$10^{2}$ | 1.275 | 0.20387 |
| **Month May:C_BM** | **5.907*$10^{-3}$** | **2.227*$10^{-3}$** | **1.819*$10^{2}$** | **2.653** | **0.00869** |
| Month August:C_BM | -2.206*$10^{-3}$ | 1.991*$10^{-3}$ | 1.856*$10^{2}$ | -1.108 | 0.26923 |

Shapiro-Wilk normality test

W = 0.98204, p-value = 0.005877

**Table ESM1.14.B.** Main model outcomes for RER.

| | Estimate | Standard Error | df | t value | P-value |
|---|---|---|---|---|---|
| **Intercept** | **0.694817** | **0.008722** | **220.426042** | **79.661** | **< 2*$10^{-16}$** |
| *Group Glucose* | *0.023328* | *0.013011* | *221.936995* | *1.793* | *0.074336* |
| *Group Methylglyoxal* | *0.020752* | *0.012446* | *222.321773* | *1.667* | *0.096863* |
| **Month February** | **0.051418** | **0.012860** | **136.585061** | **3.998** | **0.000104** |
| **Month May** | **0.028559** | **0.013887** | **153.903487** | **2.057** | **0.041423** |
| Month August | 0.020246 | 0.013887 | 153.903487 | 1.458 | 0.146915 |
| Group Glucose:Month February | -0.019934 | 0.020969 | 147.451796 | -0.951 | 0.343333 |
| Group Methylglyoxal:Month February | -0.018603 | 0.018137 | 126.334891 | -1.026 | 0.307002 |
| Group Glucose:Month May | -0.024421 | 0.022040 | 157.555561 | -1.108 | 0.269541 |
| Group Methylglyoxal:Month May | -0.027887 | 0.019005 | 146.759527 | -1.467 | 0.144426 |
| Group Glucose:Month August | -0.026283 | 0.022550 | 161.137707 | -1.166 | 0.245516 |
| Group Methylglyoxal:Month August | -0.009316 | 0.019546 | 153.712957 | -0.477 | 0.634310 |

Shapiro-Wilk normality test

W = 0.94934, p-value = $5.34*10^{-7}$

**Table ESM1.15.A** Main model outcomes for metabolic flexibility: RER AUC.

| | Estimate | Standard Error | t value | P-value |
|---|---|---|---|---|
| **Intercept** | **2.846714** | **0.054316** | **52.410** | **< $2*10^{-16}$** |
| Group Glucose | 0.005592 | 0.029748 | 0.188 | 0.851963 |
| Group Methylglyoxal | 0.018646 | 0.025889 | 0.720 | 0.476021 |
| **C_RQ** | **1.284935** | **0.306616** | **4.191** | **0.000172** |
| Age_years | 0.009025 | 0.010821 | 0.834 | 0.409787 |

Shapiro-Wilk normality test

W = 0.98166, p-value = 0.7383

**Table ESM1.15.B** Main model outcomes for metabolic flexibility: RER amplitude**.**

| | Estimate | Standard Error | t value | P-value |
|---|---|---|---|---|
| **Intercept** | **-1.10553** | **0.21673** | **-5.101** | **$1.1*10^{-5}$** |
| Group Glucose | -0.01219 | 0.11874 | -0.103 | 0.919 |
| Group Methylglyoxal | 0.03594 | 0.09721 | 0.370 | 0.714 |
| Age_years | -0.02986 | 0.04271 | -0.699 | 0.489 |
| C_BM | -0.04221 | 0.02532 | -1.667 | 0.104 |

Shapiro-Wilk normality test

W = 0.96142, p-value = 0.1763

**Table ESM1.16.A.** Main model outcomes forprobability of not flying

| | Estimate | Standard Error | z value | P-value |
|---|---|---|---|---|
| **Intercept** | **-4.6011** | **1.2608** | **-3.649** | **0.000263** |
| Group Glucose | 0.8886 | 1.6565 | 0.536 | 0.591667 |
| Group Methylglyoxal | -14.9134 | 1798.1364 | -0.008 | 0.993383 |
| Month November | 1.7203 | 1.1761 | 1.463 | 0.143531 |
| Month February | -16.0055 | 2826.7693 | -0.006 | 0.995482 |
| Month May | -0.3964 | 1.4674 | -0.270 | 0.787039 |
| Month August | -16.3126 | 2240.1363 | -0.007 | 0.994190 |
| **C_Age** | **0.8687** | **0.2718** | **3.196** | **0.001392** |
| **C_BM** | **-1.2316** | **0.4680** | **-2.632** | **0.008493** |
| Group Glucose:Month November | -0.7021 | 1.7700 | -0.397 | 0.691612 |
| Group Methylglyoxal:Month November | 14.9249 | 1798.1365 | 0.008 | 0.993377 |

| | | | | |
|---|---|---|---|---|
| Group Glucose:Month February | 18.5823 | 2826.7698 | 0.007 | 0.994755 |
| Group Methylglyoxal:Month February | 32.4586 | 3350.2118 | 0.010 | 0.992270 |
| Group Glucose:Month May | 2.8708 | 2.0615 | 1.393 | 0.163746 |
| Group Methylglyoxal:Month May | 17.1601 | 1798.1367 | 0.010 | 0.992386 |
| Group Glucose:Month August | 18.2372 | 2240.1368 | 0.008 | 0.993504 |
| Group Methylglyoxal:Month August | 32.1874 | 2872.5432 | 0.011 | 0.991060 |
| Group Glucose:C_BM | 0.8345 | 0.5549 | 1.504 | 0.132571 |
| Group Methylglyoxal:C_BM | 1.9042 | 0.5815 | 3.274 | 0.001059 |

**Table ESM1.16.B.** Main model outcomes for  mean flying velocity.

| | Estimate | St andardE rror | df | t value | P-value |
|---|---|---|---|---|---|
| **Intercept** | **1.610764** | **0.052950** | **143.994741** | **30.420** | **< $2*10^{-16}$** |
| **Group Glucose** | **-0.158919** | **0.066537** | **126.093053** | **-2.388** | **0.01840** |
| Group Methylglyoxal | 0.080992 | 0.063626 | 127.862826 | 1.273 | 0.20535 |
| Month November | -0.054246 | 0.055282 | 179.518279 | -0.981 | 0.32779 |
| Month February | -0.090830 | 0.069106 | 194.565435 | -1.314 | 0.19028 |
| Month May | -0.089943 | 0.064116 | 226.211118 | -1.403 | 0.16204 |
| Month August | -0.062524 | 0.070894 | 238.674683 | -0.882 | 0.37870 |
| C_Age | -0.001547 | 0.035283 | 62.621500 | -0.044 | 0.96517 |
| Sex M | -0.080568 | 0.049741 | 185.545223 | -1.620 | 0.10699 |
| *Group Glucose:Month November* | *-0.114481* | *0.069229* | *184.978334* | *-1.654* | *0.09989* |
| **Group Methylglyoxal:Month November** | **-0.175771** | **0.067256** | **182.977757** | **-2.613** | **0.00971** |
| Group Glucose:Month February | 0.097048 | 0.095805 | 199.522929 | 1.013 | 0.31230 |
| Group Methylglyoxal:Month February | -0.076907 | 0.090846 | 198.079111 | -0.847 | 0.39826 |
| **Group Glucose:Month May** | **0.186570** | **0.093602** | **226.962284** | **1.993** | **0.04743** |
| *Group Methylglyoxal:Month May* | *-0.135957* | *0.079742* | *233.169281* | *-1.705* | *0.08953* |
| Group Glucose:Month August | 0.075161 | 0.098378 | 239.700599 | 0.764 | 0.44561 |
| Group Methylglyoxal:Month August | -0.109770 | 0.085343 | 238.784724 | -1.286 | 0.19961 |
| **Group Glucose:C_Age** | **-0.152688** | **0.051490** | **71.162213** | **-2.965** | **0.00411** |
| Group Methylglyoxal:C_Age | -0.006571 | 0.048494 | 64.495009 | -0.136 | 0.89263 |
| **Month November:Sex M** | **0.174242** | **0.054920** | **173.892378** | **3.173** | **0.00179** |
| *Month February:Sex M* | *0.146895* | *0.075161* | *176.342693* | *1.954* | *0.05223* |
| **Month May:Sex M** | **0.172179** | **0.064184** | **176.643630** | **2.683** | **0.00800** |
| *Month August:Sex M* | *0.127330* | *0.064921* | *176.788067* | *1.961* | *0.05141* |

Shapiro-Wilk normality test

W = 0.99069, p-value = 0.09225

**Table ESM1.16.C.** Main model outcomes for m maximum flying velocity.

| | Estimate | Standard Error | df | t value | P-value |
|---|---|---|---|---|---|
| **Intercept** | **1.760679** | **0.053306** | **141.679988** | **33.029** | **$<2*10^{-16}$** |
| **GroupGlucose** | **-0.159502** | **0.076932** | **135.346061** | **-2.073** | **0.0400** |
| GroupMethylglyoxal | 0.074445 | 0.073696 | 137.604082 | 1.010 | 0.3142 |
| MonthNovember | 0.059685 | 0.061485 | 185.574702 | 0.971 | 0.3329 |
| MonthFebruary | 0.007121 | 0.077580 | 205.560952 | 0.092 | 0.9269 |
| MonthMay | -0.019221 | 0.069858 | 233.790461 | -0.275 | 0.7834 |
| MonthAugust | -0.027015 | 0.075221 | 243.942130 | -0.359 | 0.7198 |
| *C_BM* | *-0.024275* | *0.012587* | *139.593870* | *-1.929* | *0.0558* |
| C_Age | -0.007789 | 0.039127 | 56.999582 | -0.199 | 0.8429 |
| *GroupGlucose:MonthNovember* | *-0.163958* | *0.084813* | *183.411511* | *-1.933* | *0.0548* |
| **GroupMethylglyoxal:MonthNovember** | **-0.204480** | **0.082646** | **182.135351** | **-2.474** | **0.0143** |
| GroupGlucose:MonthFebruary | 0.089749 | 0.115252 | 198.559434 | 0.779 | 0.4371 |
| GroupMethylglyoxal:MonthFebruary | -0.046065 | 0.109759 | 198.815826 | -0.420 | 0.6752 |
| GroupGlucose:MonthMay | 0.167167 | 0.112422 | 225.886566 | 1.487 | 0.1384 |
| GroupMethylglyoxal:MonthMay | -0.108736 | 0.095479 | 232.476343 | -1.139 | 0.2559 |
| GroupGlucose:MonthAugust | 0.031092 | 0.117278 | 242.129453 | 0.265 | 0.7911 |
| GroupMethylglyoxal:MonthAugust | -0.081815 | 0.102581 | 243.996804 | -0.798 | 0.4259 |
| **GroupGlucose:C_Age** | **-0.136455** | **0.056430** | **64.945818** | **-2.418** | **0.0184** |
| GroupMethylglyoxal:C_Age | 0.014024 | 0.053790 | 59.445243 | 0.261 | 0.7952 |

<u>Shapiro-Wilk normality test</u>

W = 0.99347, p-value = 0.3084

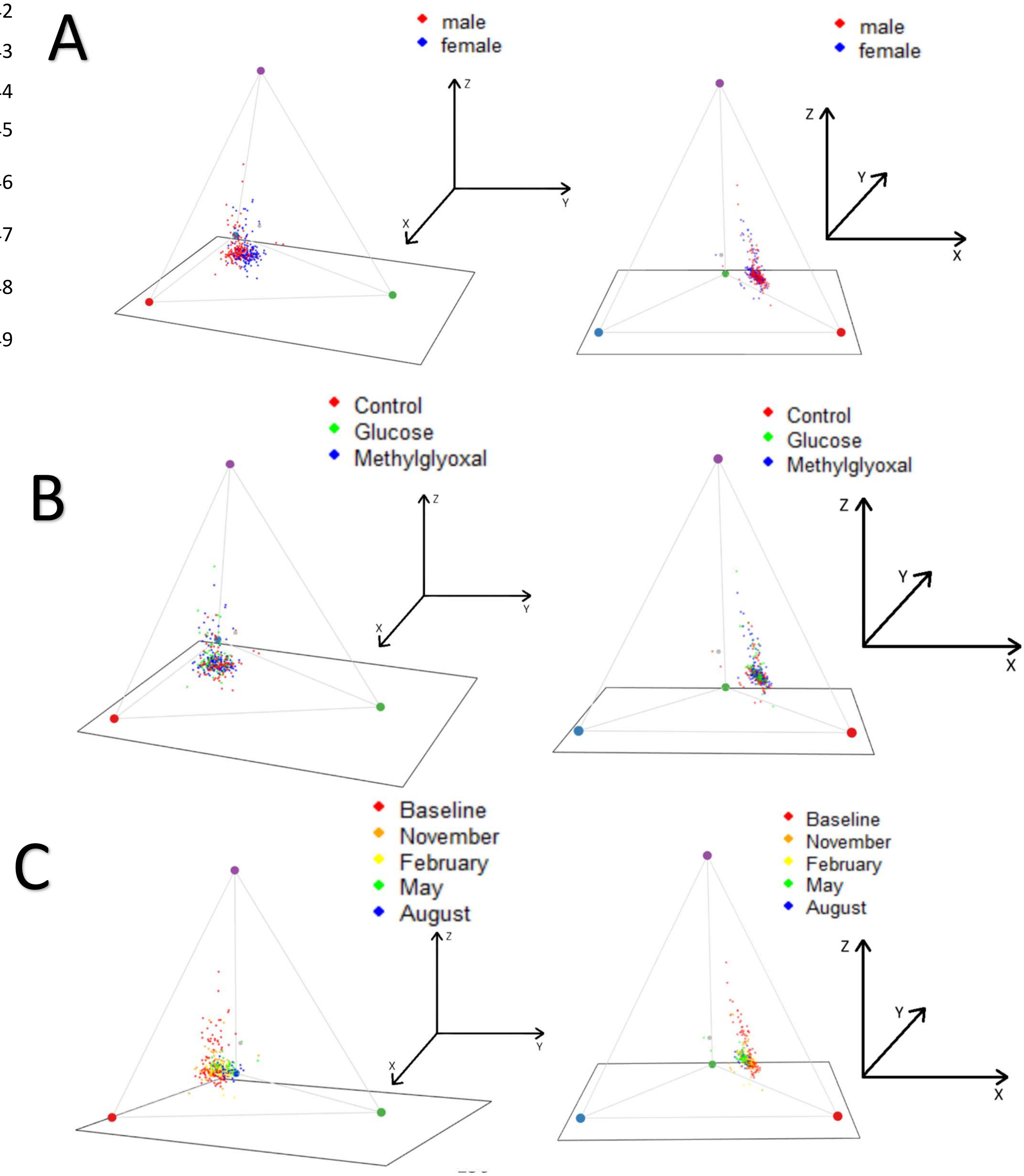


**Figure ESM1.5.** Representation of coloration values of the birds in the avian tetrahedral colour space. The vertices represent the four colour cones, i.e. red, green, blue and UV). Left, view of the tetrahedron with the red and green vertices on the front, allowing the appreciation of most of the variance. Right, view of the tetrahedron with the green vertex (i.e. higher Y values) in the back; in this view, the X coordinate is positive (i.e. redder) to the right and negative (i.e. bluer) to the left. **A.** Sex differences, with males showing redder beaks. **B.** Treatment effects. **C.** Month effects.

### Additional model results

Regarding the relationship between glucose and metabolism, RER was not affected by whole blood glucose, and blood glucose was not affected either by RMR per gram of mass. For plasma glucose levels, no significant effects of neither body mass nor RMR were found. The results of the models of blood glucose in function of temperature variables are shown in **Table ESM1.17** and **Figure ESM1.6**. No significant effects of sampling time on whole blood glucose were found ($\beta \pm SE$= -0.004 ± 0.003, P-value=0.166). For RMR variation with aviary temperature, only the (centred) minimum temperature had a negative significant effect ($\beta \pm SE$= -0.003 ± 0.001; P-value= 0.002; **Figure ESM1.15**). Breed differences were detected for body mass and muscle score. Differences in ($log_{10}$) body mass among breeds showed that the birds from unknown breed were bigger on average than all the other breeds (difference ± s.e.: Unknown - Cracow: 3.014 ± 0.475; t ratio = 6.344, P < 0.0001; Unknown - Davis: 2.535 ± 0.507; t ratio = 5.003, P < 0.0001; Unknown - Seewiesen: 3.699 ± 0.653; t = 5.664, P < 0.0001; Unknown - Bielefeld/Bielefeld-AUS: 4.253 ± 0.884; t ratio = 4.811, P = 0.0001). Differences in body mass among breeds are nevertheless generally expected (Forstmeier et al. 2007), and in our case it was likely driven by the presence of a few very big females in the unknown breed group coming from pet shops, which were above 20 g. The difference in muscle score observed between Cracow and David was small (Cracow - Davis: 0.373 ± 0.129; t = 2.889, P = 0.0375), probably unnoticeable and unlikely to be biologically meaningful. RMR also differed among breeds, with the birds from unknown breed having a higher RMR on average than all others, but this effect disappeared after when adjusting for body mass, suggesting that this difference is due to their bigger mass.

**Table ESM1.17.A** Temperature effects on glucose. All explanatory variables are centred.

| | Estimate | StandardError | df | t value | P-value |
|---|---|---|---|---|---|
| **Intercept** | **2.4806105** | **0.0608198** | **12.5404823** | **40.786** | **$1.04*10^{-14}$** |
| **Minimum Temperature** | **-0.0226522** | **0.0090057** | **28.7042546** | **-2.515** | **0.0178** |
| **Minimum Temperature$^2$** | **0.0011766** | **0.0004568** | **33.5222776** | **2.576** | **0.0146** |
| **Temperature difference** | **-0.0335545** | **0.0141628** | **13.6343640** | **-2.369** | **0.0332** |
| **Temperature difference$^2$** | **0.0019611** | **0.0007359** | **16.2846798** | **2.665** | **0.0168** |

Shapiro-Wilk normality test

W = 0.98507, P-value = 0.0138

**Table ESM1.17.B** Temperature effects on glucose. Segmented model. All explanatory variables are centred.

| | Estimate | Standard Error | df | t value | P-value |
|---|---|---|---|---|---|
| **Intercept** | **2.381371** | **0.010983** | **235.150642** | **216.821** | **<2e-16** |
| Minimum Temperature | 0.001513 | 0.001096 | 189.491804 | 1.381 | 0.169 |
| MinTpre_break | 0.013672 | 0.019135 | 219.961023 | 0.714 | 0.476 |
| **Minimum Temperature:MinTpre_break** | **-0.009076** | **0.003199** | **208.116752** | **-2.837** | **0.005** |

Shapiro-Wilk normality test

W = 0.98771, P-value = 0.03996

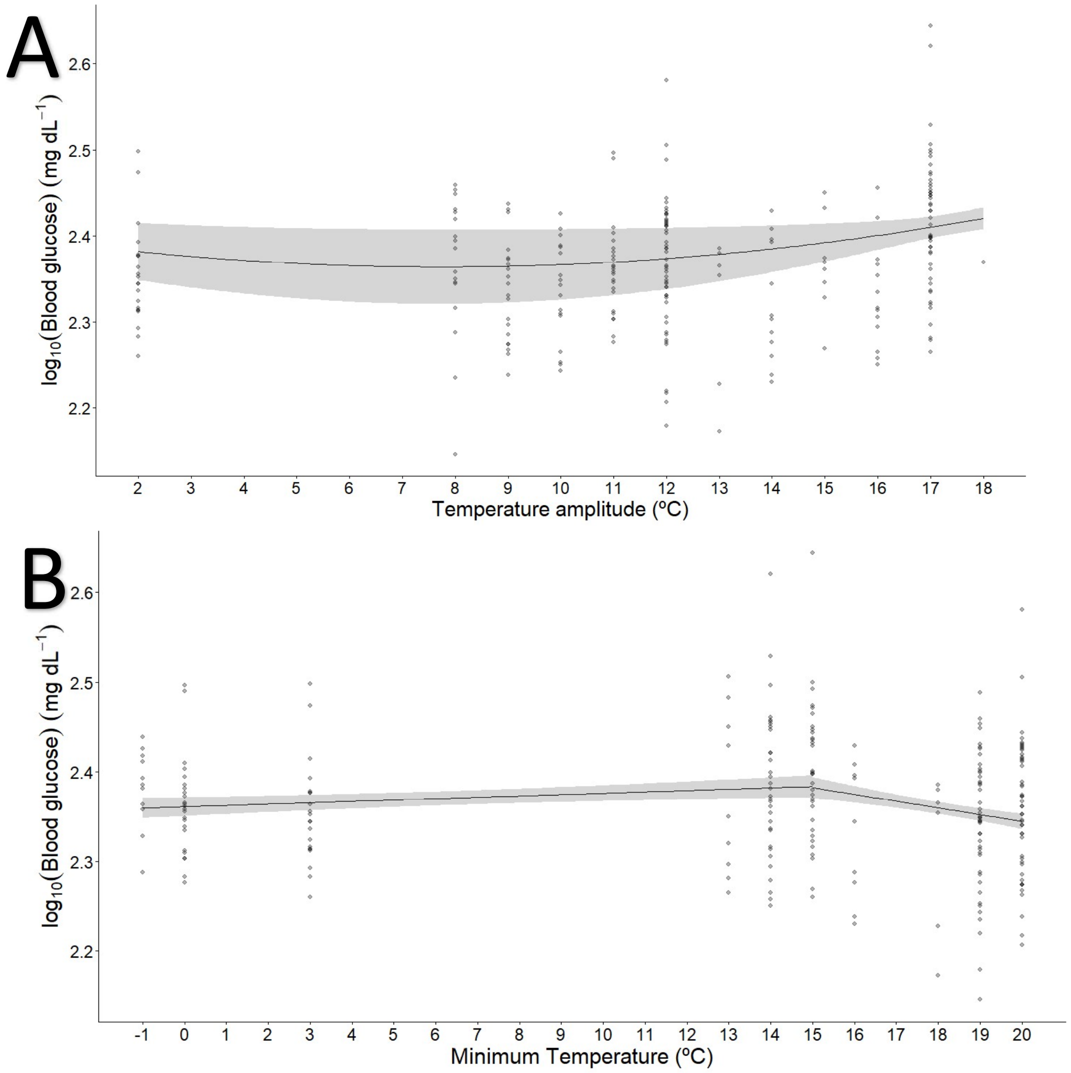


**Figure ESM1.6.** Blood glucose variation in function of **A.** daily minimum temperature and **B.** daily temperature amplitude. Predictions from the segmented models are represented. The grey area represents the 95 % confidence intervals.

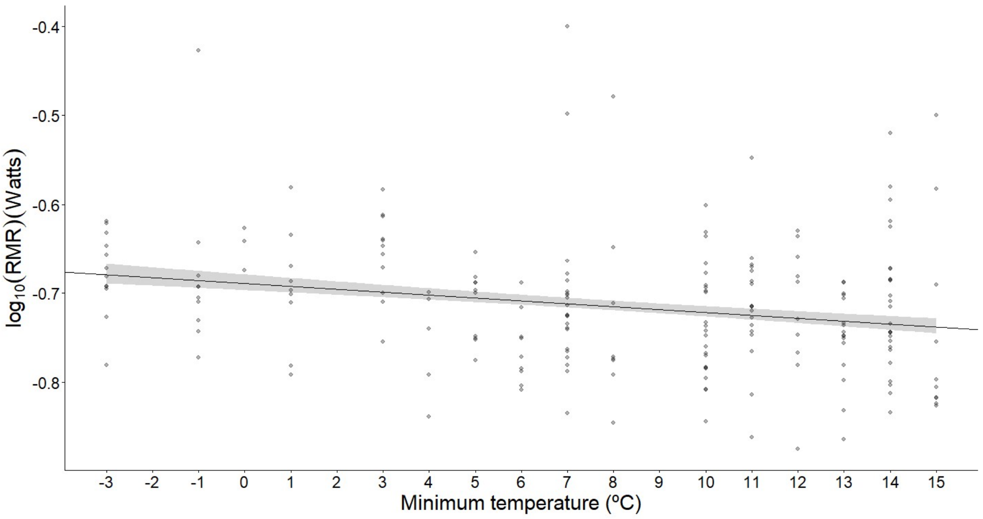

**Figure ESM1.7.** Variation of birds RMR (log10 transformed) with daily minimum temperature experienced in the aviary the day before the respirometry measurement.

### Pilot study results

#### Methylglyoxal

Water drunk by all groups was lower than in the control group (**Figure ESM1.8**).

**Table ESM1.18.** Model on water intake in the methylglyoxal supplementation experiment.

| | Estimate | Standard Error | t value | P-value |
|---|---|---|---|---|
| **Intercept** | **64.265** | **4.199** | **15.305** | **<2*10$^{-16}$** |
| **Group1** | **-44.102** | **5.938** | **-7.427** | **1.29*10$^{-9}$** |
| **Group2** | **-49.473** | **5.938** | **-8.332** | **5.13*10$^{-11}$** |
| **Group3** | **-56.344** | **5.938** | **-9.489** | **9.12*10$^{-13}$** |
| **Group4** | **-59.408** | **5.938** | **-10.005** | **1.58*10$^{-13}$** |

Residual standard error: 13.93 on 50 degrees of freedom

Multiple $R^2$: 0.7258, Adjusted $R^2$: 0.7038

F-statistic: 33.08 on 4 and 50 DF, p-value: 1.718*10$^{-13}$

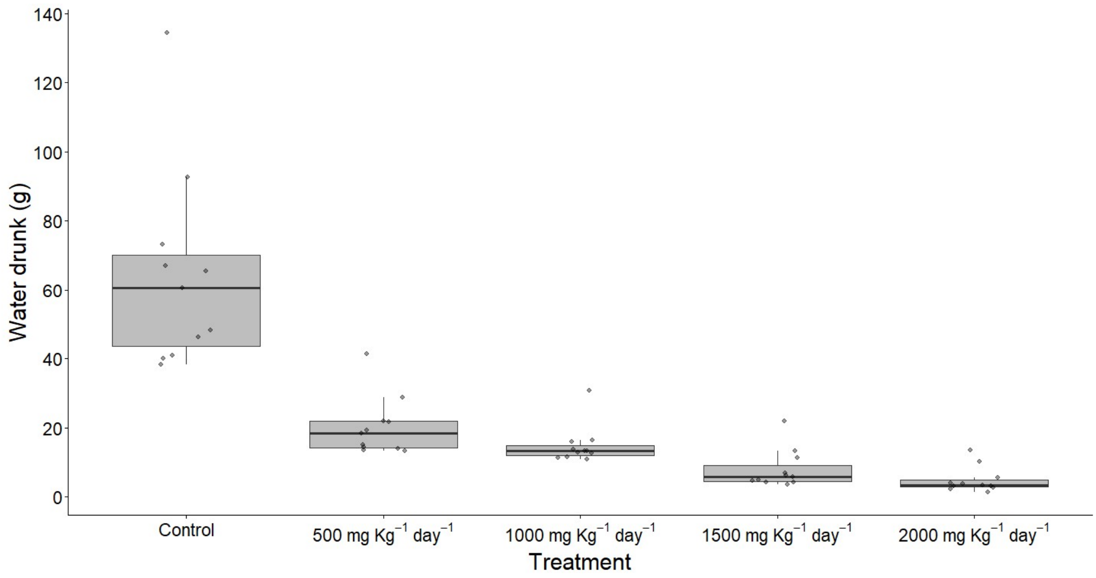


**Figure ESM1.8**. Average water drunk per day in grams by birds in each group in the methylglyoxal supplementation pilot study.

Plasma methylglyoxal levels ($log_{10}$ transformed) were significantly higher than in the control group in the medium dosages (3 and 4) during the second week of treatment (**Figure ESM1.9**).

**Table ESM1.19.** Model on plasma methylglyoxal levels.

| | Estimate | Standard Error | df | t value | P-value |
|---|---|---|---|---|---|
| **Intercept** | **5.87383** | **0.13576** | **43.77009** | **43.266** | **$< 2*10^{-16}$** |
| Cage1 | 0.24894 | 0.19199 | 43.77009 | 1.297 | 0.20156 |
| Cage2 | 0.05991 | 0.19199 | 43.77009 | 0.312 | 0.75650 |
| Cage3 | 0.06625 | 0.19199 | 43.77009 | 0.345 | 0.73169 |
| Cage4 | 0.05758 | 0.19199 | 43.77009 | 0.300 | 0.76568 |
| WeekWeek2 | -0.08429 | 0.18026 | 30.00000 | -0.468 | 0.64345 |
| WeekWeek3 | 0.20962 | 0.18026 | 30.00000 | 1.163 | 0.25403 |
| Cage1:WeekWeek2 | 0.41400 | 0.25492 | 30.00000 | 1.624 | 0.11483 |
| **Cage2:WeekWeek2** | **0.77199** | **0.25492** | **30.00000** | **3.028** | **0.00502** |
| **Cage3:WeekWeek2** | **0.75957** | **0.25492** | **30.00000** | **2.980** | **0.00567** |
| *Cage4:WeekWeek2* | *0.43872* | *0.25492* | *30.00000* | *1.721* | *0.09555* |
| Cage1:WeekWeek3 | 0.34584 | 0.25492 | 30.00000 | 1.357 | 0.18501 |
| Cage2:WeekWeek3 | 0.33609 | 0.25492 | 30.00000 | 1.318 | 0.19734 |
| Cage3:WeekWeek3 | 0.41027 | 0.25492 | 30.00000 | 1.609 | 0.11801 |
| *Cage4:WeekWeek3* | *0.51521* | *0.25492* | *30.00000* | *2.021* | *0.05227* |

<u>Shapiro-Wilk normality test</u>

W = 0.98216, p-value = 0.5266

Repeatability for Bird_ID

R = 0.119; SE = 0.124; CI = [0, 0.408]; P = 0.187 [LRT]

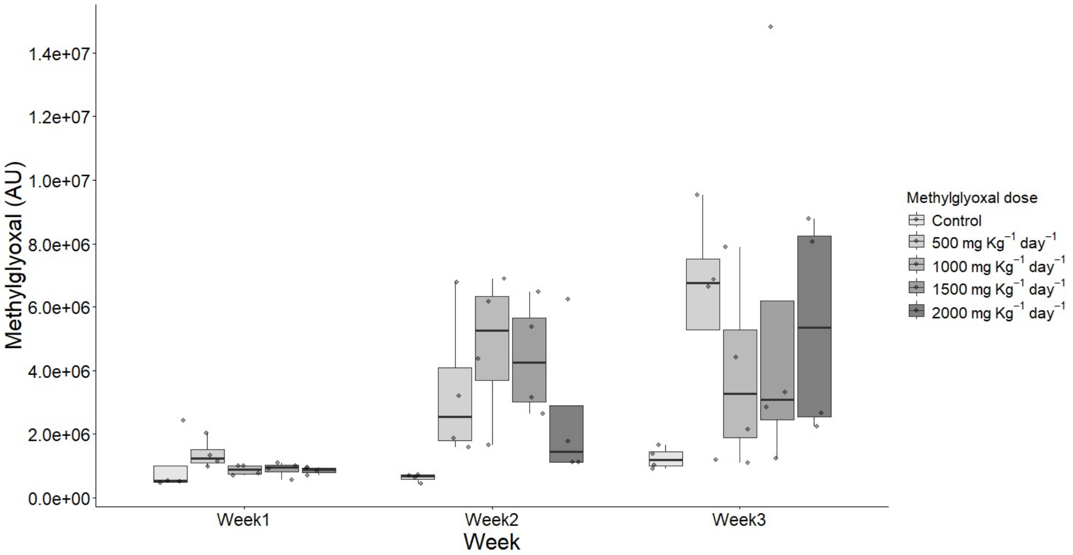


**Figure ESM1.9.** Plasma methylglyoxal levels in arbitrary units (AU) of derived compound (see **AGE section of ESM2**) from each treatment group and week from the methylglyoxal supplementation pilot study.

Body mass was lost from week 1 to week 3 in the group with highest methylglyoxal dosage (**Figure ESM1.10**).

**Table ESM1.20.** Model on body mass loss.

| | Estimate | Standard Error | t value | P-value |
|---|---|---|---|---|
| Intercept | 0.02982 | 0.04354 | 0.685 | 0.5039 |
| Cage1 | 0.03242 | 0.06158 | 0.526 | 0.6063 |
| Cage2 | 0.08826 | 0.06158 | 1.433 | 0.1723 |
| Cage3 | 0.02874 | 0.06158 | 0.467 | 0.6474 |
| **Cage4** | **-0.13783** | **0.06158** | **-2.238** | **0.0408** |

Residual standard error: 0.08708 on 15 degrees of freedom

Multiple $R^2$: 0.5018, Adjusted $R^2$: 0.3689

F-statistic: 3.776 on 4 and 15 DF, p-value: 0.02563

<u>Shapiro-Wilk normality test</u>

W = 0.97154, p-value = 0.787

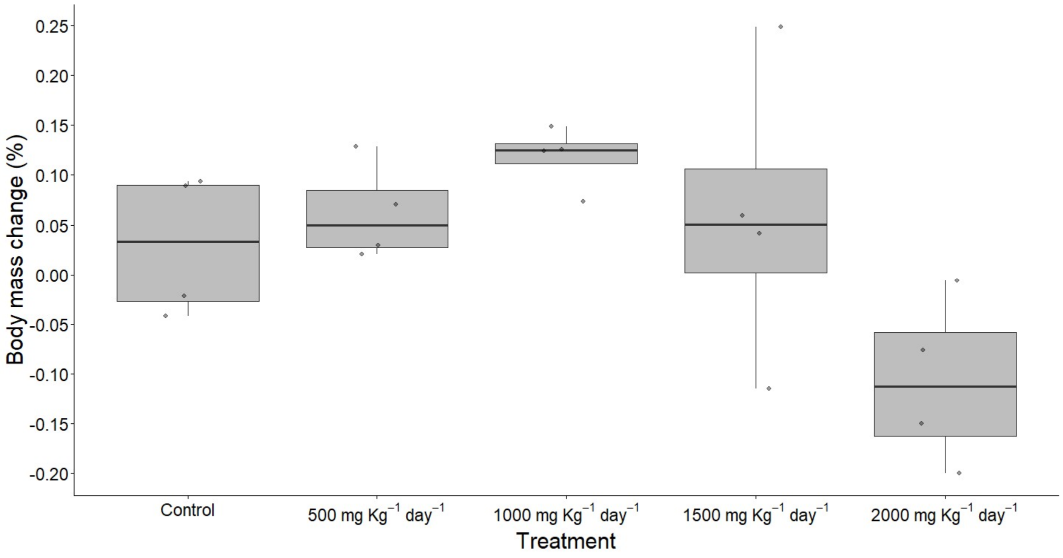


**Figure ESM1.10.** Percentage of body mass change from week 1 to week 3 of treatment at different methylglyoxal dosages.

Fat score is marginally increased in the group 2 (1000 mg $Kg^{-1}$ per day; **Figure ESM1.11**).

**Table ESM1.21.** Model on fat score.

| | Estimate | Standard Error | t value | P-value |
|---|---|---|---|---|
| Intercept | -0.7500 | 0.7746 | -0.968 | 0.3483 |
| Cage1 | 1.5000 | 1.0954 | 1.369 | 0.1911 |
| *Cage2* | *2.2500* | *1.0954* | *2.054* | *0.0578* |
| Cage3 | 1.0000 | 1.0954 | 0.913 | 0.3757 |
| Cage4 | -1.5000 | 1.0954 | -1.369 | 0.1911 |

Residual standard error: 1.549 on 15 degrees of freedom

Multiple $R^2$: 0.4842, Adjusted $R^2$: 0.3467

F-statistic: 3.521 on 4 and 15 DF, p-value: 0.03229

Shapiro-Wilk normality test

W = 0.95342, p-value = 0.422

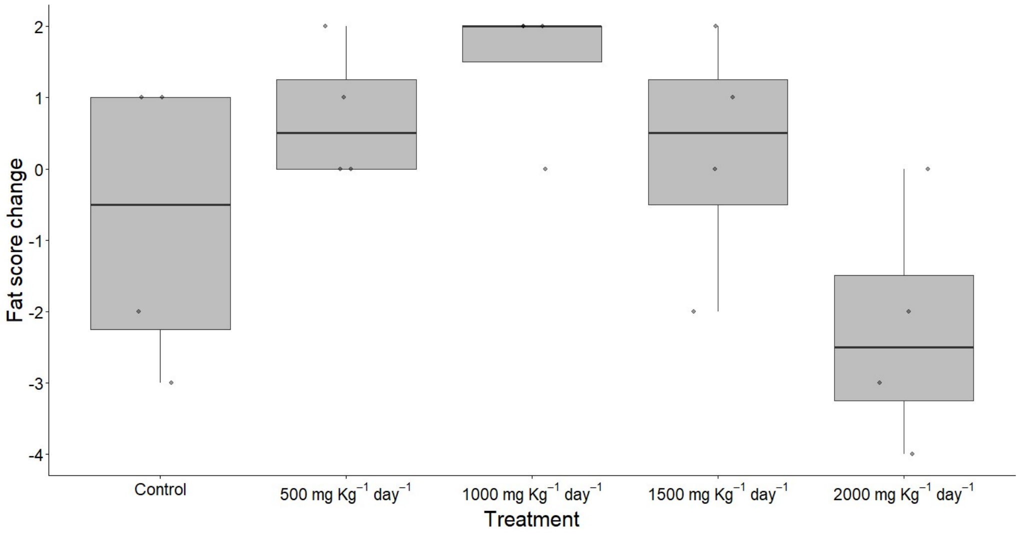


**Figure ESM1.11.** Change in fat score (see **body condition section** from Material and methods) from week 1 to week 3 of treatment (Week 1: Baseline; Week 2: after 1 week of experiment; Week 3: after two weeks of experiment) depending on methylglyoxal dosage.

Muscle score is decreased in the group 4 (2000 mg Kg$^{-1}$ per day; **Figure ESM1.12**).

**Table ESM1.22.** Model on muscle score.

| | Estimate | Standard Error | t value | P-value |
|---|---|---|---|---|
| Intercept | 0.3750 | 0.2700 | 1.389 | 0.185 |
| Cage1 | -0.3750 | 0.3819 | -0.982 | 0.342 |
| Cage2 | 0.2500 | 0.3819 | 0.655 | 0.523 |
| Cage3 | -0.1250 | 0.3819 | -0.327 | 0.748 |
| **Cage4** | **-1.1250** | **0.3819** | **-2.946** | **0.010** |

Residual standard error: 0.5401 on 15 degrees of freedom

Multiple $R^2$:  0.5028, Adjusted $R^2$: 0.3703

F-statistic: 3.793 on 4 and 15 DF, p-value: 0.02526

<u>Shapiro-Wilk normality test</u>

W = 0.92533, p-value = 0.1255

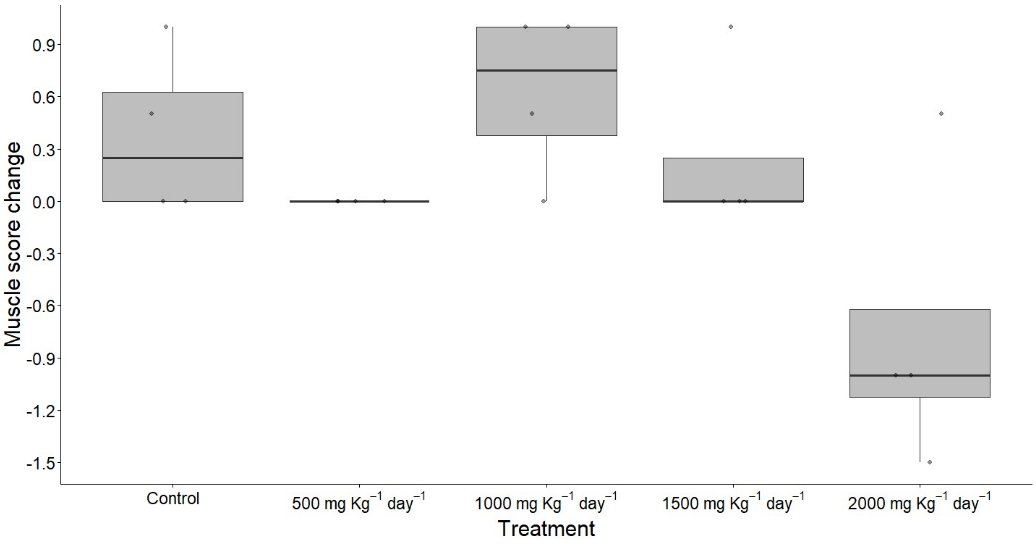


**Figure ESM1.12.** Change in Muscle score (see **body condition section** from Material and methods) from week 1 to week 3 of treatment depending on methylglyoxal dosage.

### Glucose

Food consumption is reduced by the higher glucose supplementation levels (**Figure ESM1.13**).

**Table ESM1.23.** Model on food consumption levels.

| | Estimate | Standard Error | t value | P-value |
|---|---|---|---|---|
| **Intercept** | **9.1150** | **0.3830** | **23.800** | **< $2*10^{-16}$** |
| Group1 | 0.1008 | 0.5416 | 0.186 | 0.852707 |
| *Group2* | *-0.9200* | *0.5416* | *-1.699* | *0.092663* |
| **Group3** | **-2.1267** | **0.5416** | **-3.927** | **0.000163** |
| **Group4** | **-2.3833** | **0.5416** | **-4.400** | **$2.82*10^{-5}$** |

Residual standard error: 1.713 on 95 degrees of freedom

Multiple $R^2$: 0.2785, Adjusted $R^2$: 0.2482

F-statistic: 9.169 on 4 and 95 DF, p-value: $2.62*10^{-6}$

Shapiro-Wilk normality test

W = 0.96304, p-value = 0.006662

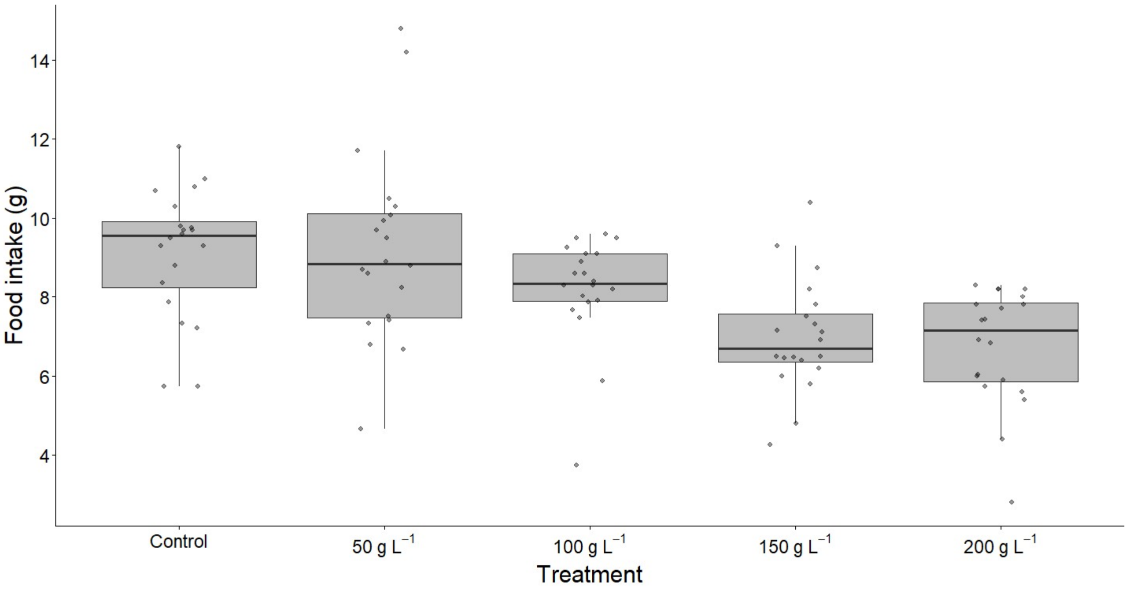


**Figure ESM1.13**. Average food intake per day in grams by birds in each group in the glucose supplementation pilot study.

Water intake is reduced by the medium (150 g $L^{-1}$ and marginally for 200 g $L^{-1}$) glucose supplementation levels (**Figure ESM1.14**).

**Table ESM1.24.** Model on water intake in the glucose supplementation experiment.

| | Estimate | Standard Error | df | t value | P-value |
|---|---|---|---|---|---|
| **Intercept** | **1.30348** | **0.08207** | **2.35534** | **15.882** | **0.00189** |
| Group1 | 0.03809 | 0.07671 | 94.00000 | 0.497 | 0.62066 |
| **Group2** | **-0.20198** | **0.07671** | **94.00000** | **-2.633** | **0.00989** |
| *Group3* | *-0.14037* | *0.07671* | *94.00000* | *-1.830* | *0.07041* |
| Group4 | -0.11682 | 0.07671 | 94.00000 | -1.523 | 0.13111 |

Shapiro-Wilk normality test

W = 0.9856, p-value = 0.3508

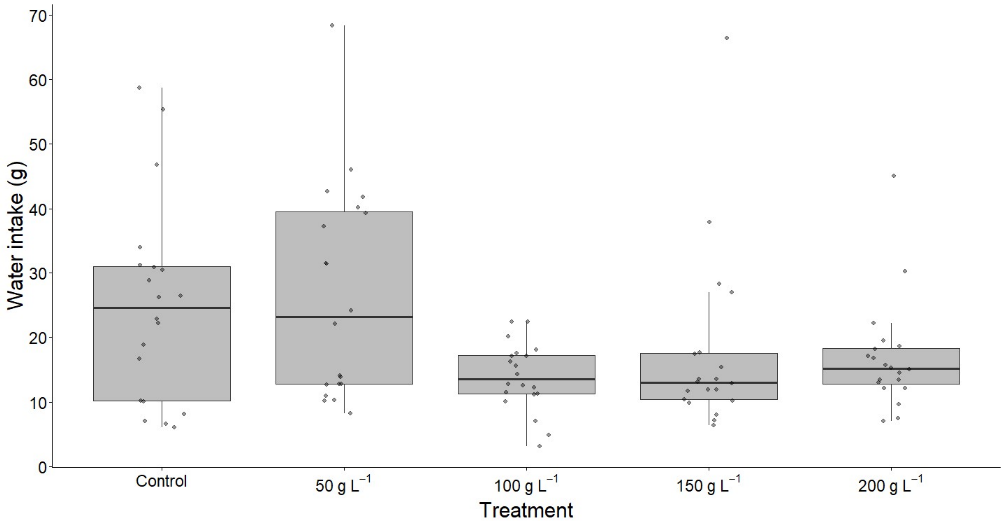


**Figure ESM1.14**. Average water drunk per day in grams by birds in each group in the glucose supplementation pilot study.

Whole blood glucose levels are not consistently affected by the treatment (**Figure ESM1.15**), but decrease with fasting time (**Figure ESM1.16**).

**Table ESM1.25.** Model on blood glucose levels.

| | Estimate | Standard Error | df | t value | P-value |
|---|---|---|---|---|---|
| **Intercept** | **2.3823261** | **0.0296387** | **31.2861532** | **80.379** | **< $2*10^{-16}$** |
| Group1 | -0.0291927 | 0.0418272 | 31.1632874 | -0.698 | 0.490393 |
| Group2 | 0.0004765 | 0.0420833 | 31.5176554 | 0.011 | 0.991038 |
| Group3 | -0.0606215 | 0.0418612 | 31.2106512 | -1.448 | 0.157545 |
| Group4 | -0.0256376 | 0.0424066 | 31.9564816 | -0.605 | 0.549735 |
| Week2 | 0.0457600 | 0.0315750 | 28.0476208 | 1.449 | 0.158357 |
| Week3 | 0.0085119 | 0.0314882 | 28.0036253 | 0.270 | 0.788896 |
| **C_Timespan** | **-0.0005315** | **0.0001468** | **36.2437976** | **-3.621** | **0.000891** |
| Group1:Week2 | -0.0056128 | 0.0447020 | 28.0648267 | -0.126 | 0.900975 |
| Group2:Week2 | -0.0443495 | 0.0440327 | 27.8217065 | -1.007 | 0.322521 |
| **Group3:Week2** | **0.0977451** | **0.0445642** | **28.0155800** | **2.193** | **0.036745** |
| Group4:Week2 | -0.0271011 | 0.0446832 | 28.0581126 | -0.607 | 0.549049 |
| Group1:Week3 | 0.0261806 | 0.0441076 | 27.8493713 | 0.594 | 0.557596 |
| Group2:Week3 | -0.0366166 | 0.0453704 | 28.2980873 | -0.807 | 0.426365 |
| Group3:Week3 | 0.0612403 | 0.0440447 | 27.8261254 | 1.390 | 0.175425 |
| Group4:Week3 | 0.0193426 | 0.0467803 | 28.8802155 | 0.413 | 0.682312 |

<u>Shapiro-Wilk normality test</u>

W = 0.98307, p-value = 0.5824

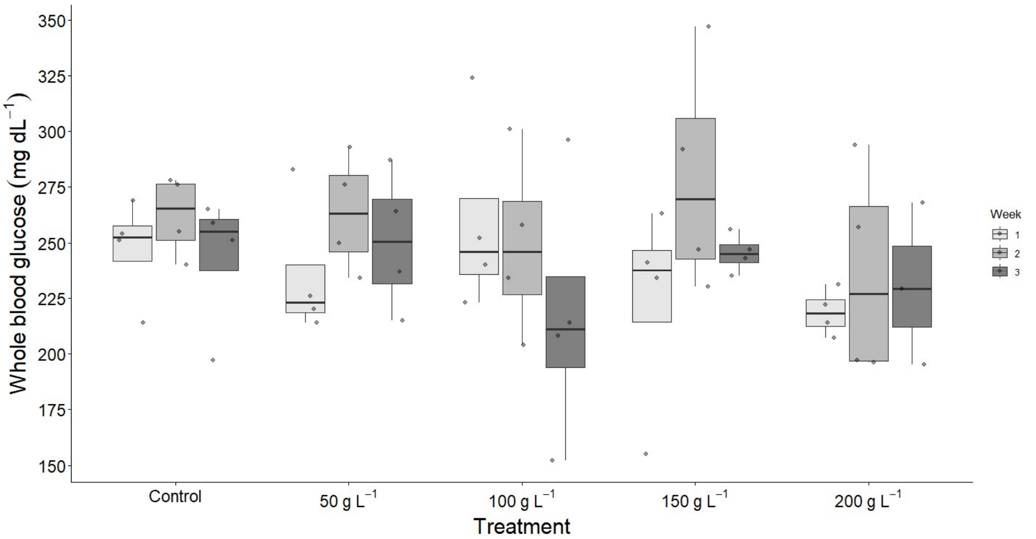


**Figure ESM1.15**. Whole blood glucose in mg dL$^{-1}$ of birds in each group across the experiment weeks (1-3; i.e. from baseline to two weeks after the start of the experiment; see **Figure ESM1.11** caption) in the glucose supplementation pilot study.

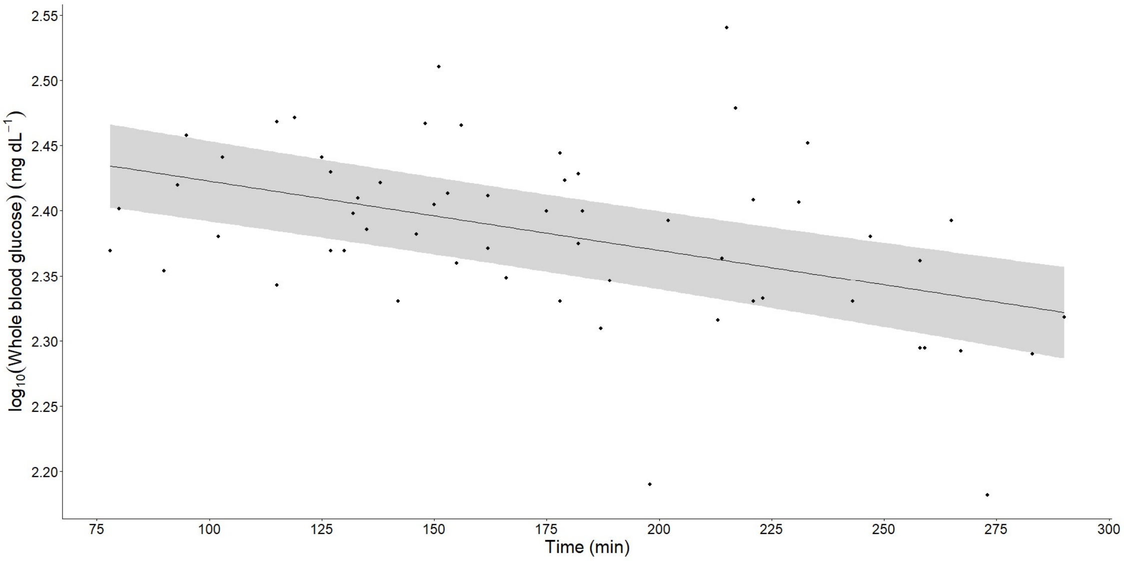


**Figure ESM1.16**. Variation in $\log_{10}$(whole blood glucose) levels (mg dL$^{-1}$) in function of the time in minutes at which it was measured, counting from the start of the fasting period, in the glucose supplementation pilot study.
