## Supplementary material for "Decoupling glycation from mortality: glucose, but not methylglyoxal, reduces survival in zebra finches": ESM2 - Methods

#### Housing conditions

The aviaries included several roosting perches and branches and, except from late summer, the floor was covered by twigs and tree bark, contributing to environmental enrichment. A shelter with an infrared light working in permanence most of the year, except late spring and summer, was set up in each aviary for shielding birds against cold. The lateral fences that delimit each aviary were coated with cardboard in winter and bedsheets during summer, so that birds from one group could not see or interact with the contiguous group, be altered by the researchers or technicians when going inside other cages, and to protect them from cold in winter.

#### Flight performance measurements

What is described in this section constitutes an adaptation from Reichert et al., 2015.

Flight performance, which is directly linked to an individual’s capacity to forage and escape predators, is used as a proxy for individual maintenance (Lindhe Norberg 2002). It was measured using vertical take-off when alarmed, with a similar set up to that used in previous studies (Criscuolo et al., 2011; Kullberg et al., 2002; Reichert et al., 2015). Each bird is released on a perch situated at 20 cm from the ground, at the base of a transparent vertical plastic tube covered by a metallic grid to make it possible for the bird to see its borders and not to try to escape through it (**Figure ESM2.1**). At the top of the flying tube (120 cm from the first perch), there is a perch where the bird can be collected after each flight. The birds were released five times in the base of the tube and allowed to rest for 30 seconds between each flight. If the bird did not complete any flight after 5 min, or spent more than 5 min in total without flying during all the trials, it was recorded as “no flight” and the trial was terminated to avoid unnecessary stress for the animal. During the experimental trial, all flights were recorded with a camera (Canon EOS 7D, see **Coloration**) or, on its defect, with a mobile phone. To determine flight speed, the videos were observed and the time the birds take to cover a distance of 40 cm is calculated. This measure is made by counting the number of video frames (recordings are made at approximately 30 fps, thus covering 0.033 s frame$^{-1}$) between two marks on the tube at 40 and 80 cm height. Flight speed is calculated with the formula:

$$V = \frac{d}{[F * 0.033]} \quad (1)$$

where *d* is the distance between the two marks (40 cm), *F* is the number of frames the bird has spent between both marks. The fastest of the five flights is taken as a measurement of the bird's maximum escape flight ability (Kullberg et al., 2002). The time to start flying was also registered. 12 randomly sampled (by the means of R `sample()` function) birds that had been kept indoors in small cages (40.5 cm x54.5 cm x29.5 cm) during the previous night were measured each day (**see Main text**) on the morning (Tuesday-Friday), from 9:00 to 12:00 approximately. They were released afterwards in their corresponding outdoors aviaries.

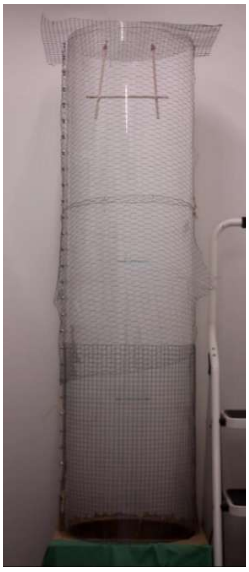

**Figure ESM2.1**. Tube used for flying test. The bird is put in the bottom through the curtain and startled by clapping, making it fly up to the top perch. The speed is then calculated by the number of frames spent between the marks.

### Respirometry

RMR was measured at 35ºC because the zebra finch thermoneutral zone, have been reported between 33ºC and 38ºC (Calder 1964; Briga and Verhulst 2017; Wojciechowski et al., 2021), in post-absorptive state (determined for zebra finches as described below), in resting but awake individuals (McNab 1997). Even if we tried to meet these criteria, it is usually heavily argued if BMR is achievable, as some authors inquire that an actual metabolic nadir has to be reached, often after the animal has spent some time inside the metabolic chamber, and assuming it is not growing or investing any energy into reproduction (the latter not being totally controllable in our case, particularly for females, that may be producing eggs at some parts of the experiment), in order to call the measurement BMR, so we will call it Resting Metabolic Rate (RMR) here on. Resting metabolic rate (RMR) was measured by oxygen consumption, carbon dioxide production, and water vapour using a flow-through system with excurrent flow (**Figure**

**ESM2.2.A**). The system included an Oxzilla (full-cell) oxygen analyser (Sable Systems International, SSI, Las Vegas, Nevada, USA), a CA-2A infrared carbon dioxide analyser (SSI), a RH-300 water vapour analyser (SSI), five metabolic chambers (530 ml, with hermetic lid fitted with inlet and outlet ports, four for the birds and one empty for the baseline; **Figure ESM2.2.B**) and therefore five channels. We used an eight-channel multiplexer (TR-RM8, SSI) to control the flow of air to each of the four chambers holding individual birds, and to the chamber used for baseline recordings. Airflow through the chambers was maintained by the multiplexer even when the metabolic rates were not being measured. The excurrent air flow rate was 650 ml $min^{-1}$ maintained by five independent pumps (Sierra, one per chamber) with their respective flowmeters, placed right upstream the pumps. Before placing the birds inside the metabolic chambers, they were placed in small wire cages (**Figure ESM2.2.C**), to facilitate their handling and rapid exchange from the system and prevent contact with the mineral oil placed at the bottom of the metabolic chamber, which was used to collect cloacal fluid and prevent the production of excess water vapour resulting from the evaporation of the cloacal fluid. The base of the wire mesh cage was placed 2 cm above the bottom of the metabolic chamber. A mathematical correction was performed on $O_2$ and $CO_2$ values, to control for the water vapour dilution effect (Equation 8.6 from Lighton 2008). We calculated the respiratory exchange ratio (RER, whole body VCO2/VO2 ratio) as a proxy of the respiratory quotient (RQ, CO2/O2 from substrate oxidation), which constitutes an index of the substrate being consumed (i.e. carbohydrates, proteins or lipids), and therefore the metabolic state of the animal (Péronnet and Massicotte, 1991).

To measure RMR all metabolic chambers with de birds inside were introduced in a 30 L temperature-controlled cabinet (PTC-1, SSI, **Figure ESM2.2.D**). To keep the temperature inside the cabinet at around 35°C, we used a temperature controller (Pelt-5, SSI). The temperature controller of the cabinet was turned on about an hour before the start of the measurements to reach the target temperature. Birds were fasted for at least 240 min before being placed in a metabolic chamber to ensure that they were in a post-absorptive state. Birds were weighed (to 0.1 g; Sartorius, USA) and the body temperatures were measured (to 0.01ºC, Testo 925) at the start and the end of each experiment. Body temperature was measured by inserting a type-K thermocouple approximately 1 cm in the cloacae. This allowed us to adjust oxygen consumption to mass (as average mass from both measures) and control for possible thermal stress. Body condition parameters were also measured right before (see **Body condition**) and coloration afterwards (see **Coloration**). Water vapor and carbon dioxide were removed from the incurrent air using a single filter for all chambers with Drierite® or silica gel and soda lime, respectively, before entering the metabolic chambers, placed in the following order (in the sense of the flow): drier, soda lime and again the drying agent, as soda lime produces water when reacting with $CO_2$. Cotton wool or fabric filters were placed between these layers to avoid the mixing of the products. Water vapour was reduced to a level always lower than 10% of water vapour saturation. The filter was changed when the colour indicators of the drier and the soda lime had turned pink/violet respectively, showing they were saturated. Both drierite/silica gel and soda lime were regenerated every day putting it in a Plexiglass tray for 2 hours inside an oven at 210º C. We measured 3 groups of 4 birds every day, starting at around 14:00 for the first group. Each measurement lasted one hour, comprising two bouts of 30 min each, i.e. 6 min for each individual and for a baseline (divided into two spans of 3 min at the beginning and end of each round), with the multiplexer programmed to record the excurrent flow from each metabolic chamber sequentially, including de baseline. Birds were given 15 minutes of habituation inside of the chambers before starting the measure, which also served for the temperature and humidity inside the chambers to return to their adequate values. Recordings were taken every 1 s for each bird during each cycle.

The respirometry system was turned on several hours before starting the measurement in order to reach the adequate stationary state needed for the sensors to work properly. We randomly sampled 12 birds by R (see general protocol) and they were placed indoors in small cages (40.5 cm x54.5 cm x29.5 cm) at 20ºC and the same photoperiod they were experiencing in outdoor aviaries cages the day before of experiments in order to avoid thermal stress and maintain the same conditions right before the measurement, regardless of outdoor temperature, which would vary with the season (see **main text**). They were provided unrestricted food and still water in the indoor cages, but the food was removed at 9:00 in the morning, in order for them to be in postprandial state when measured in the afternoon. This was determined by a preliminary test in which the $O_2$ consumption and RER were measured in a group of 4 birds from 9:00 to 18:00, showing a reduction from a RER of about 1 (representing generally carbohydrates catabolism and associated with absorptive state) at the start up to around 0.7 (representing mainly fat catabolism and associated with fasting) (see e.g. Walsberg and Wolf, 1995; McCue et al., 2011) after about 3 hours of measurement (**Figure ESM2.3**). As occasional birds were detected storing seeds in their crops up to about 3 hours after feeding, we finally set a period of at least 4 hours of fasting before metabolic assessments. The room in which respirometry is performed is separated by a wall from the respirometry system to decrease the perturbation to the animals and it was kept in the dark during the measurements. After finishing, we left the birds indoors in the same cages they spent the previous night in order to be used for flying tests (see above) the next day, except in the case of Fridays, in which they would be released to their aviaries. The total procedure lasted a maximum of 8 days, distributed into two weeks (Tuesday-Friday) for each round of measurement performed every 3 months (see **Main text**).

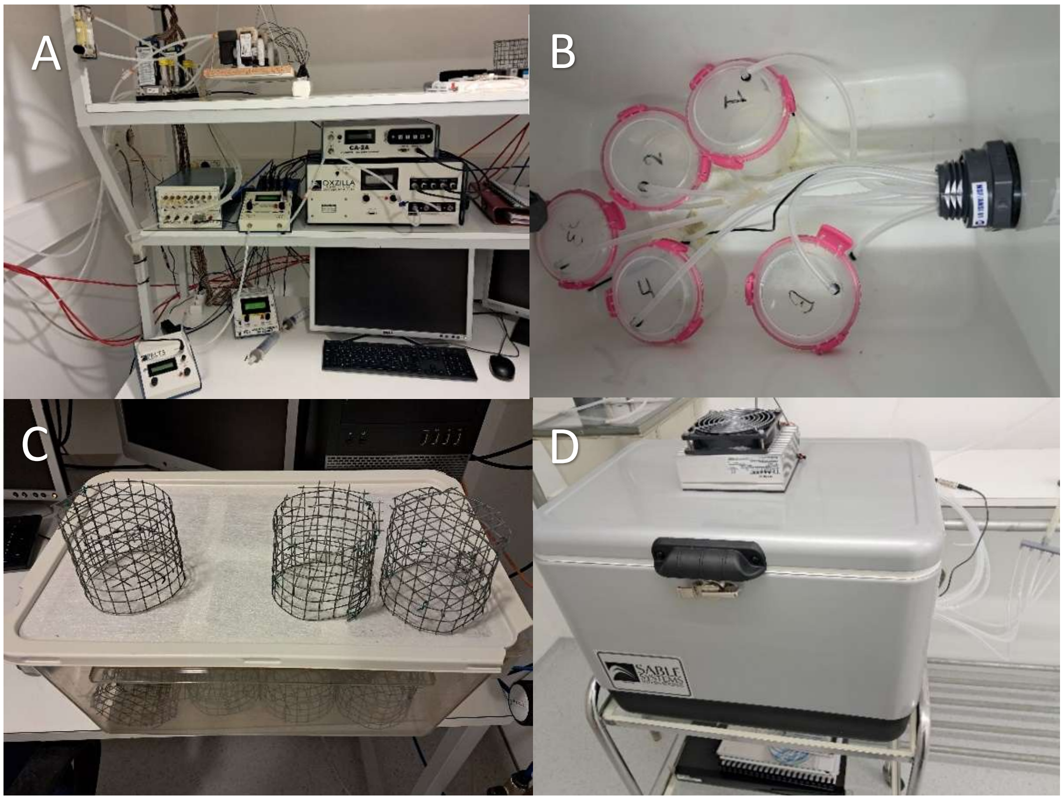


**Figure ESM2.2.A.** Sable System® respirometry system with (**B**) metabolic chambers. Wire cages (**C**) were used within the chambers to restrain the birds during the measurement. **D.** Temperature controlling cabinet.

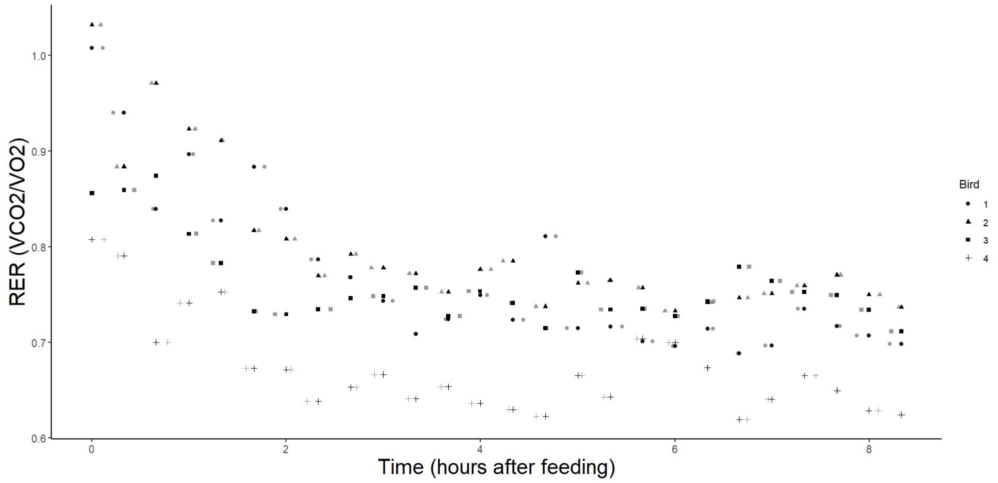


**Figure ESM2.3.** Respiratory Exchange Ratio variation over time from a preliminary test on a group of 4 zebra finches. Birds were placed in a cage in the indoors animal facilities and given free access to food and water, and then put into a respirometry chamber to measure RMR and RER.

Data were recorded and processed using the ExpeData software package (SSI), with a macro correcting the lag for each channel ($H_2O$ was measured first, then $CO_2$ and finally $O_2$), the water vapour dilution and performing the baseline correction. The average $O_2$ and $CO_2$ of the most stable 180 seconds within each 6 min measurement spans of each cycle were calculated (determined automatically by the software) and averaged to be used as the consumption values for those birds. An average RER was then calculated from those values. RMR were calculated using equation (3) from Lighton et al. (1987), modified by dividing the $VO_2$ by 60 to convert it to mL $s^{-1}$ and then report the RMR in Watts.

Metabolic flexibility (sensu e.g. McKechnie 2008; Norin and Metcalfe 2019; Swanson et al. 2022) was also assessed with two parameters: AUC and amplitude. The first is calculated as the sum of all RMR values, while adjusting for the initial RMR (November) in the models (see **Statistics**), as higher RMR would imply higher total sum independently of the AUC. Amplitude is estimated as the difference between initial and maximum RMR, i.e. between RMR in November and the highest RMR of all measurement rounds (normally February).

#### Coloration

Beak coloration was measured by the means of photography with a Canon EOS 7D modified for full spectrum sensibility, to be able to detect UV. A Novoflex Novoflexar objective, with a considerable transmittance for UV, was used, together with a CMOS-optimized UV/IR-filter (Baader UV/IR Cut) or visible-IR spectrum filter (Baader U-filter), for each of the two photos to be taken. 12 birds were photographed each day, in groups of 4, corresponding to those of respirometry bouts, and being performed after each round of it (see **Main text**). The photographs were taken in a room without windows, with a lightbulb resembling daylight with an adequate UV production (Osram Ultravitalux UV-A) hanging at 80 cm over the table, with the abovementioned camera set up in a tripod over a table with a green cloth covering its surface to avoid reflections. Two photos were taken consecutively to each bird, one with each filter. The objective focus was set at the 1.5 feet mark of the objective (between 40 and 50 cm marks). The camera was parallel to the surface and its settings adjusted manually with an ISO of 400, a white balance of 4100 K and the metering being performed over the surface of a Spectralon (by Labsphere®) diffuse white standard (95% grey reflectance), placed over the table, used to standardize the luminance. Another grey Spectralon standard with 10 % reflectance was photographed under the same conditions to control for possible over exposition of the white standard. A calliper was included into the photos in order to provide a size reference, and the bird to be photographed was held with the left hand and therefore with the right side of the beak upwards and parallel to the surface (**Figure ESM2.4)**.

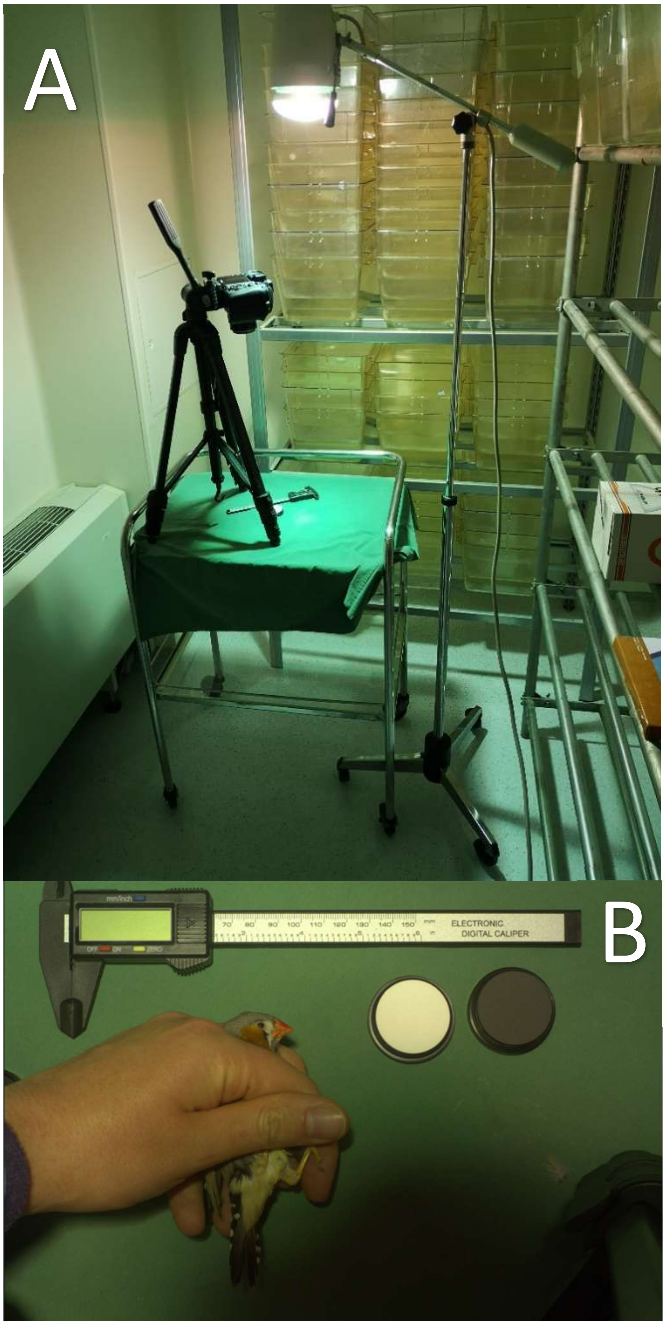


**Figure ESM2.4.A.** Set up used for the photography. **B.** Example of bird picture.

Photos were stored as .CR2 files (RAW files from Canon®) in the camera SD card, and later on transferred to a computer and processed with micaToolbox ImageJ plugin, developed by Troscianko & Stevens (2015). This toolbox extracts 4 colour channels (R, G, B and UV), with both photographs of each bird being superposed in a multispectral stack, manually aligned, and signals automatically linearized and normalized, considering the sensitivities of the camera (available on the program) and the characteristics of the light source (D65 used). The toolbox can also model the animal perception of the image with a colour space specifically developed for each species. Here, blue tit (*Cyanistes caeruleus*) was used, as is the closest species available within the program, used as a model species for passerine birds. This way, cone catch quanta values (for uv, long, medium, short wavelength and double cones) were obtained from the photos taken. The values were posteriorly expressed as coordinates in a tetrahedral avian vision space, by using equation 20 from Endler & Mielke (2005), in which the Euclidean distances between individual values can be estimated. These coordinates are X, Y, Z (as it is a three-dimensional space). An additional "luminosity" value can be obtained from the double cone absorption values. X represent the axis going from short (blue) to long (red) wavelength, being higher the redder the hue, Y the axis from the middle point of the previous axis to medium wavelength (green), being higher the greener the hue, and Z represents the position between the middle of all the previous three (similar to a human grey point) and UV (or violet for other bird species), being higher the more UV is the hue (**Figure ESM2.5**).

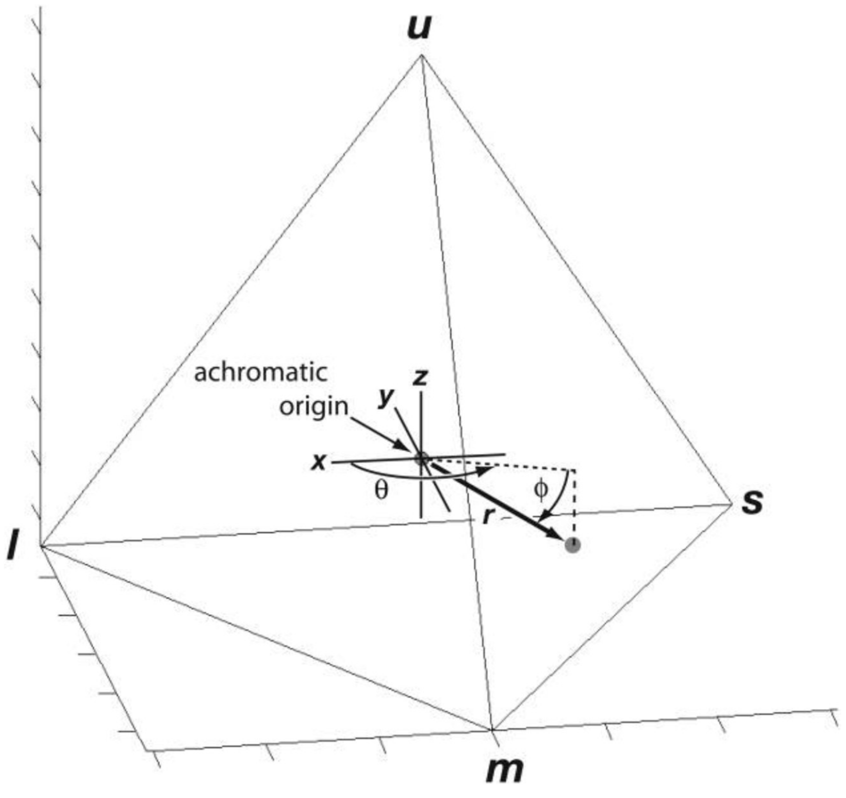


**Figure ESM2.5.** Tetrahedral space with its x, y and z axes and the vertices representing the pure "primary" colours in the tetra-chromatic visual system modelled (i.e. s=short wavelength, m=medium wavelength, l=long wavelength, u=UV). Image from Stoddard and Prum (2008).

After the photos had been taken, birds were left in the same small cages (40.5 cm x54.5 cm x29.5 cm) in which they were placed before respirometry, and in which they will stay overnight, for flying tests to be performed with them during the following day (see **Main text**).

### Body condition

12 birds randomly sampled (see **Main text**) were assessed each day just before respirometry. They would be grabbed from a cage (40.5 cm x54.5 cm x29.5 cm) in which they had remained indoors for the previous night and morning (see **Main text** and **Respirometry**) and weighed by the means of a Sartorius scale with a maximum resolution of 0.1 g. For that the bird was placed upside down into a plastic tube with which the scale would have been previously tared. Afterwards, a measure of both muscle and fat conditions were performed, following a score from 0 to 8 in the case of fat and from 0 to 3 in the case of muscle, such that the higher the score the more fat or muscle they had. These scores were assessed by blowing in ventral and lateral sides of the birds to remove the feathers and visually check the degree of development of each of them, and giving them a number depending on the following charts (adapted from Aranzadi Science Society):

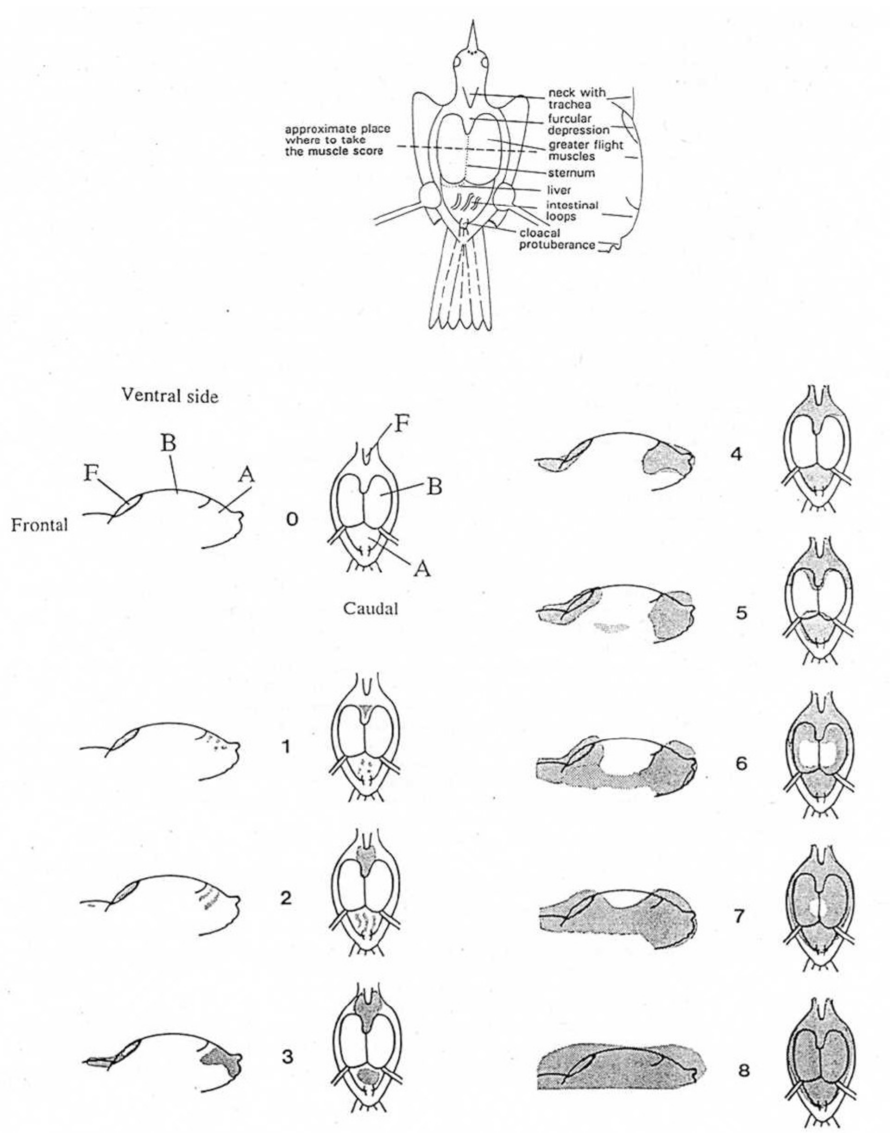


**0.** Without distinguishable fat.

**1.** Traces. Clues of fat (dark zone).

**2.** Concave interclavicular region with visible fat. Abdominal fat in traces.

**3.** Interclavicular depression flat, fully covered. Belly still not fully covered.

**4.** Fully covered abdominal region.

**5.** Convex interclavicular fat. Abdominal fat covering part of the breast muscle.

**6.** Fat, visible in the sides of the breast muscle, that joins the interclavicular and abdominal fat. Partially covered musculature.

**7.** Only a small part of the breast muscle visible.

**8.** Body fully covered of fat. Musculature not visible.

If the bird’s fat or muscle did not fit a specific level of the table, because of excess or defect, it was adjusted with fractional values of 0.5 over or under the given integers.

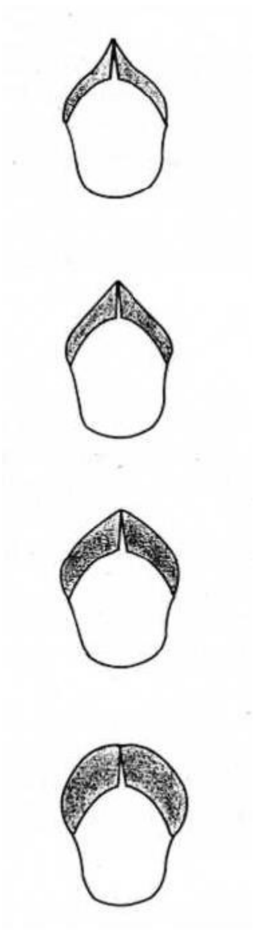

0. Quill very patent. Concave musculature.

1. Quill distinguishable. Flat musculature.

2. Quill still distinguishable. Convex muscles.

3. Quill undistinguishable. Very convex muscles.

Translation and adaptation from Aranzadi Sciences Society.

#### Blood sampling

Birds were bled twice every measurement round (each three months, see **Main text**), within two consecutive weeks (i.e. one sampling per week). The birds were left indoors in small cages (40.5 cm x54.5 cm x29.5 cm), with no more than 6 birds per cage, the previous evening without food, so that they were fasting.

Blood samples were taken by brachial vein puncture with a lancet and a heparinised Microvette®. After the extraction of around 100 µL (varying depending on the size and health status of the birds), the birds were returned to a cage with free access to food and water, until being released in their outdoor aviaries later that day. Sampling always started in the morning, beginning at as early as around 7 a.m. (although usually at around 9 a.m.) and ending at maximum at around 18:30, although normally much earlier (80 % of the cases within a range of less than 4 hours, and 60 % under 2 hours; see **Statistics** section for discussion on possible stress effects). Blood was kept in ice after collected, to be centrifuged at 4ºC and 3500 g for 10 min right after the end of the sampling in order to separate plasma and cells. The plasma was separated into aliquots for future analyses and all samples were stored at -80ºC until measures were performed.

### AGE

A subset of plasma sample aliquots from every treatment groups at the periods previously indicated (see **Main text**) were processed in order to obtain the extracts to be subsequently analysed by targeted metabolomics. Methylglyoxal, glyoxal, lactate, pentosidine, CML (Carboxymethyl-lysine) and CEL (Carboxyethyl-lysine) were targeted in the analysis (**Figure ESM2.6**). For this, methylglyoxal, glyoxal and lactate were first derivatized (i.e. transformed into measurable derivatives) to enable their detection. This derivatization was performed in aliquots of 15µL from each sample, to which 70µL of methanol with 100 ng $mL^{-1}$ of internal standard (D-Abscisic Acid: D-ABA) was added. Then, it was incubated for 2 h at -80°C and centrifuged 10 minutes at 4°C and 13200 rpm, from where 50 µL of supernatant were recovered. Finally, 1µL of reagent (o-phenylenediamine) 500µg $mL^{-1}$ was added, and the resulting samples were incubated overnight at 4°C, after which they were analysed by ultrahigh-performance liquid chromatography (UHPLC) on the UltiMate 3,000 UHPLC system (Thermo) coupled to an EvoQ Elite (Bruker) mass spectrometer equipped with an electrospray ionization (ESI) source in MS/MS mode. For this, 5 µL from each extract were injected, on an Acquity UPLC ® HSS T3 C18 column (2.1 × 100 mm, 1.8 µm, Waters) coupled to an Acquity UPLC HSS T3 C18 precolumn (2.1 × 5 mm, 1.8 µm, Waters) for chromatographic separation. Samples were carried through the column following a gradient of solvent A ($H_2O$; 0.1% formic acid) and B (methanol; 0.1% formic acid) at a flux of 300 µL $min^{-1}$, starting with 2% B for 0.5 minutes, reaching 100% B at 8.5 minutes, holding 100% B for 3 minutes and returning to 2% B in 1 minutes, for a total run time of 13.5 minutes. The column was operated at 35 °C. Nitrogen was generated from pressurized air by a Nitro 35 nitrogen generator (GenGaz) and used as cone gas (20 L $h^{-1}$), heated probe gas (25 L $h^{-1}$) and nebulizing gas (25 L $h^{-1}$). The cone and heated probe temperatures were 300°C and 350°C, respectively, and the capillary voltage was set at 3.5 kV. Metabolites were analysed by multiple reaction monitoring (MRM) after determining the retention time and mode (positive or negative) by scan and the cone voltage, daughter ion and collision energy using the MRM builder function on standards. Washing solvent (80% $H_2O$, 20% MeOH) was used to wash the syringe. A mix of the different standards served as a positive control. Standard curves made with standards of known concentration of each compound were used to determine the concentration of each of the sample. As glyoxal and methylglyoxal were derived, the only compounds for which the real concentration (in ng $mL^{-1}$) of the original molecules were estimated were CML and CEL.

Unfortunately, pentosidine measurements did not work, and lactate values showed a big number of values under the limit of detection, with also an unbalanced distribution, which strongly difficulted its proper modelling, so they were subsequently not considered for further analyses. These models were performed with the same independent variables as the ones described for other dependent variables, but without including the random factor for the model on the zeros. As these models were found to show convergence issues, they were finally not considered.

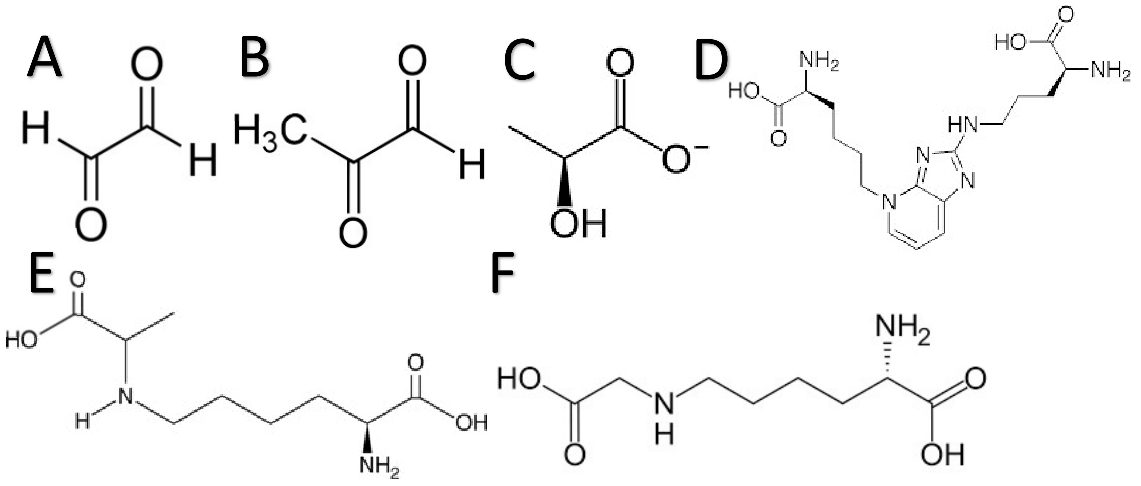


**Figure ESM2.6.** The products measured in the targeted metabolomics procedure. **A.** Glyoxal. **B.** Methylglyoxal. **C.** Lactate. **D.** Pentosidine. **E.** CEL (Carboxyethyl-lysine). **F.** CML (Carboxymethyl-lysine).

### **Albumin glycation**

Glycation levels were quantified by liquid chromatography coupled to mass spectrometry (LC-MS), a gold standard approach for assessing protein glycation (e.g. Priego-Capote et al. 2014) that has also previously been used in avian studies (Ingram et al. 2017; Zuck et al. 2017; Brun et al. 2022). Samples were analysed after completion of the experiment, late 2023. A single sample per individual and time point was analysed. Only a subsample from all the birds was selected, given logistic limitations on the total number of samples that could be processed. Briefly, 3 µL of plasma were diluted with 22 µL of distilled water containing 0.1% of formic acid, and 5 µL of that solution were injected onto a column of reverse-phase chromatography (Vydac 208TP C8; i.d. 2.1 × 250 mm, 300 Å, 5 µm particle size; Grace, Columbia, MD, USA) using an Agilent 1200

Series HPLC system (Agilent Technologies, Paolo Alto, USA) coupled to a quadrupole-time-of-flight (Q-TOF) mass spectrometer equipped with an electrospray source (maXis II, Bruker Daltonik GmbH, Bremen, Germany). For each albumin form, glycated and native, extracted ion chromatograms were obtained by summing the signals of the ten most abundant charge states. Peak areas were used to calculate relative abundances of each form, and glycation levels were obtained as the percentage ratio of glycated to total albumin peak area (see **Figure ESM2.7**). The glycation values used in the statistical analyses represent the total percentage of glycated albumin, obtained by adding the percentages of singly and doubly glycated albumin. Further details on the method and data processing can be found in (Brun et al. 2022). Individuals were excluded from the statistical analyses when spectra were too heterogeneous to resolve albumin forms or albumin glycation values were not properly detected (i.e. zero glycation levels reported). This decision is made based on the fact that it is not possible to know why the glycated albumin signal is under the limit of detection (i.e. if this is a "real value" or a technical issue). This is so as it is not even possible to determine reliably a limit of detection per se, as these are proportional values calculated by dividing peak areas (see above), and this limit may also be different for each species or even for each sample, depending on relative albumin abundance (which is nevertheless irrelevant for the calculation of the index). Moreover, albumin glycation levels are normally well above 0 in all the species we measured, which makes us think that this is an artefact, and therefore discard these data, at least until a strong reason to discriminate between these options is found. These still constituted a minority of the cases.

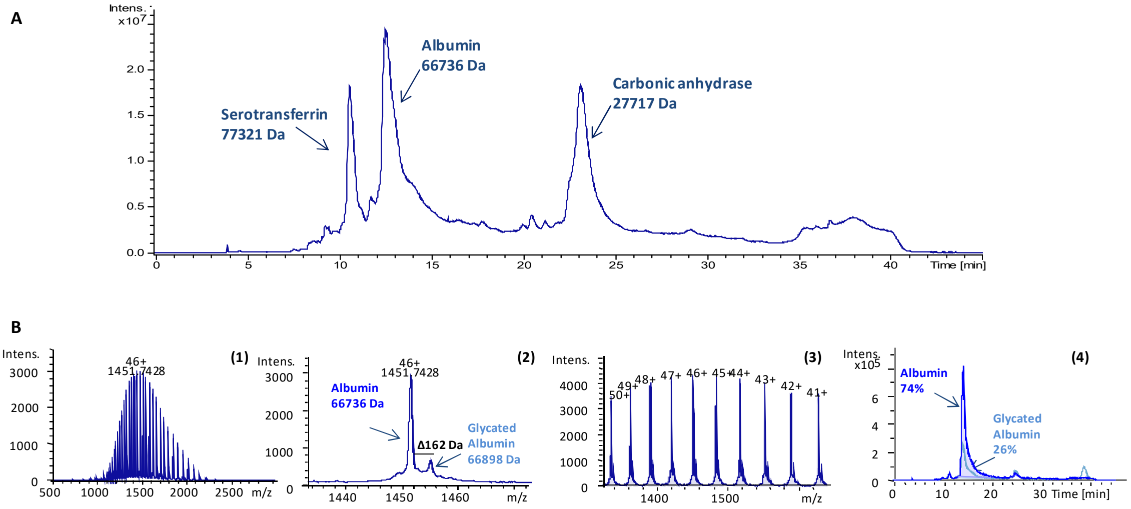


**Figure ESM2.7. A** Typical chromatogram of a zebra finch plasma sample showing the main identifiable proteins. **B** Schematic of the data processing workflow used to obtain glycation values. First, (**1**) the albumin mass spectrum is selected and (**2**) zoomed in to identify native and glycated albumin based on their masses and the Δ *m/z* of 162 Da corresponding to glucose. Then, (**3**) the ten most intense ions are selected and extracted ion chromatograms are generated. (**4**) The area under the curve is calculated for each form to determine the relative percentage of native and glycated albumin.

### Body composition

Individuals that died during the experiment were collected and kept frozen at -20ºC until dissection, taking place in two sessions during 2023. A total of 48 birds died during experiment, with 43 corpses conserved in appropriate conditions to use them for body composition assessments. Liver and breast muscle (*pectoralis major*) were separated and weighed to the nearest 0.0001 g in a precision scale along the remaining rest of the body, referred hereon as "carcase". After that, they were lyophilized by freezing at -20ºC and subsequent sublimation by applying near void conditions (pressure 0.006 bar) during approximately two weeks and then weighed again, so the water proportion of each of the parts (liver, *pectoralis* and carcase) could be estimated, allowing us to test for possible dehydration or water retention effects of the treatments. This weighing was repeated once a week until the mass was stable. In 2024, these portions were independently ground with a Retsch ZM 200 and the resulting powder stored at room temperature for later assessment of lipid content by the Folch method. For this, 1g of the aforementioned powder from each sample in duplicate, was introduced into labelled round vessels and 30 ml of a 2:1 mix of chloroform and methanol (CM mix hereon) was added, in order to separate the lipids from the samples. After that, these samples were exposed 30 s to ultrasounds and stirred with a glass bar (these were always rinsed with 1ml of the CM mix to recuperate the remaining on it) in order to break the cells. Subsequently, the samples were agitated during 12 hours and then filtered to remove the lumps while transferred to tubes. All the possible remaining present in the previous recipients and filters/funnels used were carefully recuperated by rinsing them with 2 ml of the CM mix. 8 ml of KCl at 0.9 % (polar compound) was added to the tubes with the samples and they were stirred as before to separate the methanol, followed by 10 min of centrifugation at 2000 rpm. After this, the superior phase (methanol with KCl) is aspirated by a void pump and the remaining transferred to a previously weighed round vessel, rinsing the tube with 2 ml of chloroform to recuperate any remainder. The samples were then left in a Büchi 461 Rotavapor (bath at 44ºC and later cooling at 8ºC) rotating at 100 rpm at -760 mm Hg, until the chloroform evaporates. Finally, the vessels with the isolated lipids were left overnight in a stove at 50ºC and then weighed to calculate the percentage of lipids in the original sample. A second extraction with a subsample equally distributed across the different experimental groups was performed on the remainders left in the paper filters previously used, in order to confirm that all lipids were adequately extracted in the first session. For this, the filters were shredded into small pieces and placed into vessels with 40 ml of CM mix, repeating the previous protocol, but with 10 ml of KCl 0.9 % this time.

The lipid percentage was calculated by subtracting the weight of the empty vessel from that of it with the extracted material, dividing this by the initial weight of the sample and multiplying by 100. The values obtained during the second extraction were averaged within groups and added to the individual values from the first extraction.

### **Pilot study**

20 birds (10 males and 10 females; 4 of them later reused on the main study) were used in a pilot study performed from February to March 2022 (i.e. about 4 months before the main study baseline measurements) to determine the dosages of both glucose and methylglyoxal that were going to be used for the main experiment. These individuals were housed indoors at 12:12 L:D cycles and approx. 22ºC in battery cages (8 equal cages per battery; battery dimensions: 100 x 40 x 183.5 cm) in groups of two per cage (1 male and 1 female), and distributed in 5 experimental groups of 4 birds each: a control group with still water and 4 different groups with water including either glucose or methylglyoxal at different concentrations (depending on the study session; see below): 50, 100, 150 and 200 g $L^{-1}$ for glucose (i.e. second session), and 6.886, 13.773, 20.659 and 27.545 mL of methylglyoxal (pure) per L of total solution (i.e. 7.119, 14.238, 21.356 and 28.475 mL of the commercial solution with 40% methylglyoxal in water, respectively; see main text), representing intended dosages of 500 mg $Kg^{-1}$ per day, 1000 mg $Kg^{-1}$ per day, 1500 mg $Kg^{-1}$ per day, 2000 mg $Kg^{-1}$ per day. The methylglyoxal concentrations were calculated from these intended dosages assuming a body mass equal to the average body mass of the birds (15.745 g), and drinking rates as those determined before the start for birds housed in the same conditions but without treatment. All of them had access to *ad libitum* food consisting into a commercial seed mix (Deli Nature® for tropical finches).

The pilot study was performed in two sessions: first, a 2-week session (01/02/2022-15/02/2022) with methylglyoxal supplementation, and later a 2-week session (28/02/2022-14/03/2022) with glucose supplementation (i.e. with a washout of two weeks in between). The birds were randomly assigned to the abovementioned groups using R `sample()` function, independently for each of these sessions, with always 2 males and 2 females per group. An additional group of 2 water feeders without birds was added to control for water lost by evaporation and feeder manipulation. Water and food were weighed every day during these treatments, and final values were averaged within groups, and for water also corrected for average loss by evaporation and manipulation spilling.

Body mass and body condition (see body condition) were measured and blood samples from brachial vein taken at the beginning of the experiment (i.e. before any supplementation) and after the first and the second week of supplementation, to compare either whole blood glucose or plasma methylglyoxal values. The methods for blood sampling were the same as for the main experiment (see **Blood sampling** section). Glucose was measured by a glucometer (Ascensia Countour® plus) on a small drop of whole blood and plasma methylglyoxal by the same means as on the main experiment (see **AGE section**). The samples were performed in the morning after placing all the birds together in a single cage covered by a cloth and without access to food nor water for an hour and then randomly grabbing them to avoid correlation between sampling time and treatment group. The effect of (centred) sampling time was also included in the model. One datapoint was extremely high (~600 mg $dL^{-1}$), and the bird showed signs of pathology (i.e. low activity levels, apparent somnolence and high cloacal temperature) and had recently laid an egg that was found broken. It was therefore considered an outlier and removed from the statistical analyses. This allowed for the model residuals to follow a normal distribution with minor changes on the effects.

Results (see **Table ESM1.25**) showed no significant effects of glucose supplementation on whole blood glucose levels, except for 150 g $L^{-1}$ at the second week only, accompanied by a moderate reduction in food intake and drinking, while methylglyoxal supplementation increased plasma methylglyoxal levels, but with a high reduction in drinking rates. Sampling time had a significant negative effect on glucose levels. The lowest concentrations (50 g $L^{-1}$ of glucose and 500 mg $Kg^{-1}$ $day^{-1}$ of methylglyoxal) were selected for the final experiment, to reduce to the maximum the effects of dehydration that could be confounded with the effects of the supplementation (e.g. increased mortality), especially in the methylglyoxal group, and the severity of the treatment in the glucose group, given the lack of consistent effects on glucose levels over time.

#### Statistics

The main models performed are summarized in the following box (**Box ESM2.1**), with their final structure:

##### Legend

Age_H= Horizontal component of age (i.e. age of the bird

Age_L= Longitudinal component of age

C_*XXX*= Centred variable (e.g. C_BM = Centred body mass)

**Box ESM2.1. Main models and their corresponding final equations.**

**Whole blood glucose.** log10(Glucose) ~ Group*Month + (1| Bird_ID)

**Plasma glucose.** Glucose ~ Group*Month + (1| Bird_ID)

**Albumin glycation.** Glycation ~ Group*Month+C_Glucose*Month + (1| Bird_ID)

**Methylglyoxal.** log10(Methylglyoxal_derived) ~ Group*Month + (1| Bird_ID)

**Glyoxal.** log10(Glyoxal_derived) ~ Group*Month+Group*Sex +C_Age_years*Month+ (1| Bird_ID)

**CML.** log10(CML_ng_ml) ~ Group*Month+Group*C_Age_years+ (1| Bird_ID)

**CEL.** log10(CEL_ng_ml) ~ Group*Month+C_logGlyoxal + (1| Bird_ID)

**Body mass.** log10(AVG_BM) ~ Group*Month +Sex+ C_Head_beak + C_Tarsus +(1| Bird_ID)

**Body condition: fat score.** Fat ~ Group*Month+ C_Age_years+ (1| Bird_ID)

**Body condition: muscle score.** Muscle ~ Group*Month +Sex+ (1| Bird_ID)

**Body composition: carcass fat.** log10(Lipid_mass) ~ C_logDry_mass + Group + Sex

**Body composition: carcass water.** log10(Water)~C_logFresh_mass*Sex+Group

**Body composition: liver fat.** log10(Lipid_mass) ~ C_logDry_mass + Group + Sex

**Body composition: liver water.** log10(Water)~C_logFresh_mass+Group

**Body composition: muscle (pectoralis) fat.** log10(Lipid_mass) ~ C_logDry_mass + Group + Sex

**Body composition: muscle (pectoralis) water.** log10(Water)~C_logFresh_mass+Group

**Metabolism: RMR.** RMR ~ Group*Month+Group*C_Age_years+C_BM*Group+C_BM*Month + (1| Bird_ID)

**Metabolic flexibility: AUC.** Sum ~ C_RMR + C_Age_years + C_BM + Sex

**Metabolic flexibility: Amplitude.** Amplitude~C_Age_years+C_BM+C_RMR

**Metabolism: RER.** RQ ~ Group*Month + (1| Bird_ID)

**Metabolic flexibility: RER AUC.** Sum_RQ~Group+C_RQ+C_Age_years

**Metabolic flexibility: RER amplitude.** Log10(Amplitude_RQ)~Group+C_Age_years+C_BM)

**Flying performance: Mean flying velocity.** I(Mean_V == 0) ~ Group*Month + C_Age+C_BM*Group, family = binomial(link = "logit")

Mean_V ~ Group*Month + C_Age*Group+ Sex*Month + (1 | Bird_ID)

**Flying performance: Maximum flying velocity.** V_Max ~ Group*Month + C_BM + C_Age*Group + (1 | Bird_ID)

#### <u>Coloration</u>

The effects of the experiment on beak coloration were analysed by a Bayesian multivariate LMM framework, with the **`MCMCglmm()`** function, having the response formed by the combination of all the values of the coordinates in the abovementioned tetrahedral space (X, Y and Z, see **coloration section**) as a dependent variable. This, together with the exclusion of the general intercept and the inclusion of an intercept for each coordinate, both for the general and the random part of the model, allows for an estimation of the variation of each coordinate with the explanatory variables. Treatment, sex and age were included as explanatory variables, with two models: one with horizontal age and month and another with longitudinal and initial age. Four and five level interactions were not retained in any of the final models, and the only interactions kept in the model testing for longitudinal age were those involving such variable.

An additional MRPP model, similar to what is advised by Endler and Mielke (2005), was performed with a modification of the **`mrpp()`** function of the **`vegan`** R package (Oksanen et al. 2025), including ID as the function **`strata`** argument, and the interaction groups (i.e. the groups composed by the interaction of treatment, month and sex) as a grouping factor. This way, the significance of the differences between each of those particular groups can be estimated, beyond the effects of the factors in the previous models.

#### Additional models

Additional models were performed to determine the effect of either RMR or RMR by gram of mass on whole blood glucose (WBG) and plasma glucose. The effects of RMR and body mass on whole blood glucose (all $\log_{10}$ transformed) were also tested (see results in **ESM1**). Finally, models were performed to estimate the effects of the ambient temperature in the outdoor aviaries on blood glucose levels and RMR (both response variables $\log_{10}$ transformed; see results in **ESM1**), including both (centred) daily minimum temperature and (centred) daily temperature difference (maximum temperature minus minimum temperature) recorded on the day before the blood sampling (given the birds were put indoors the day before). These variables were chosen as maximum temperature correlates (positively) with temperature difference more strongly (Multiple $R^2$=0.6377, Adjusted $R^2$=0.6372; slope ± SE = 0.4385 ± 0.0125; P-value<0.0001; **Figure ESM2.8.A**) than minimum temperature (Multiple $R^2$= 0.1466, Adjusted $R^2$= 0.1454; slope ± SE = 0.32267 ± 0.02945; P-value<0.0001; **Figure ESM2.8.B**), so minimum and difference can be enough (and would be better) to describe the temperature variation. For two days for which there was no record for the previous day, the temperature from either 2 days or 3 days before was used, considering there had not been severe temperature changes in the surrounding days and therefore the used values could be similar to the observed right the day before. The quadratic components of these variables were also tested in the complete models, and removed if not significant. For glucose, as the quadratic effects were significant, a segmented model was performed for minimum temperature (see **ESM1** for results). Body mass was included as a covariable in the models estimating the effects of temperature on RMR. Bird ID and measurement date were included as random factors, as measurements were not evenly distributed across the year, but clumped around specific dates.

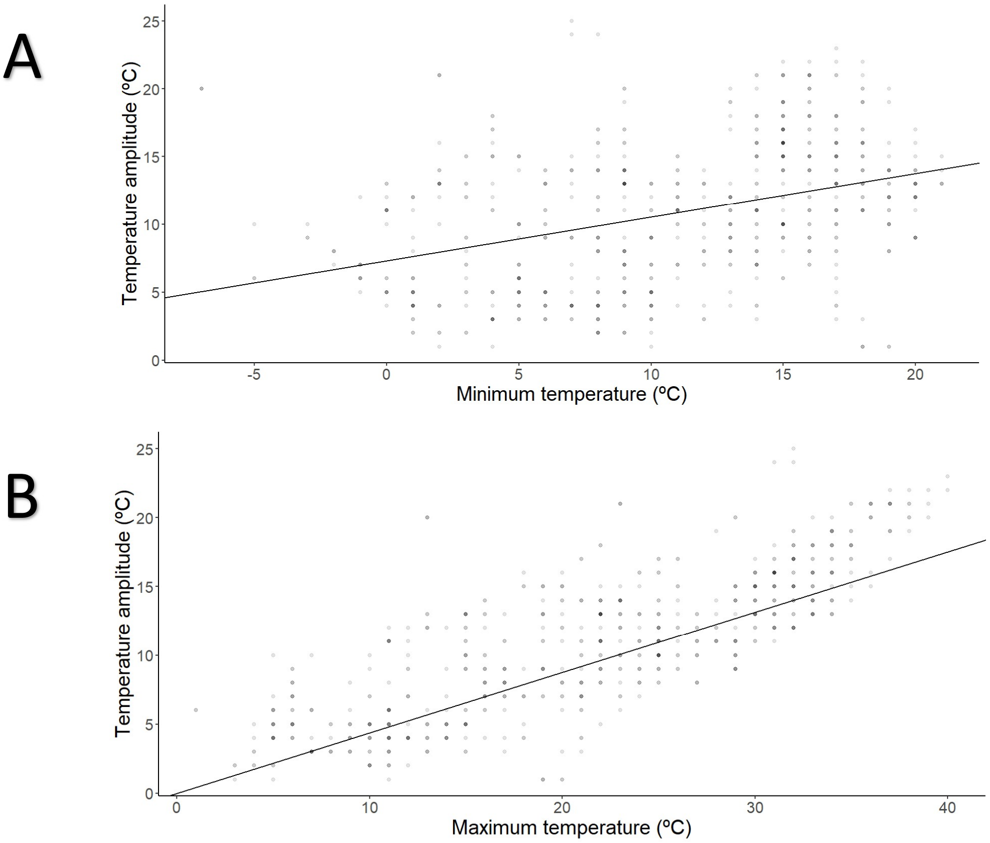


**Figure ESM2.8.** Correlation between **A** minimum daily temperature and **B** maximum daily temperature and daily temperature amplitude in the outdoor aviaries where the birds were located during the experiment. The number of values for a given temperature is represented through the intensity of the dots.

#### Stress effects

To test for possible effects of stress response on blood glucose levels, a model with $\log_{10}$(Glucose) as the dependent variable and sampling time in hours (counting as 0 the time of the first bird sampled each day) as an explanatory variable, including bird ID and sampling day as random factors, was performed on the whole dataset. The total sampling time for a day was normally under 4 hours (within a morning), except for a few birds in the baseline, that were measured in another session in the evening, at around 6 and 8 hours after the first sampled bird (**Figure ESM2.9**).

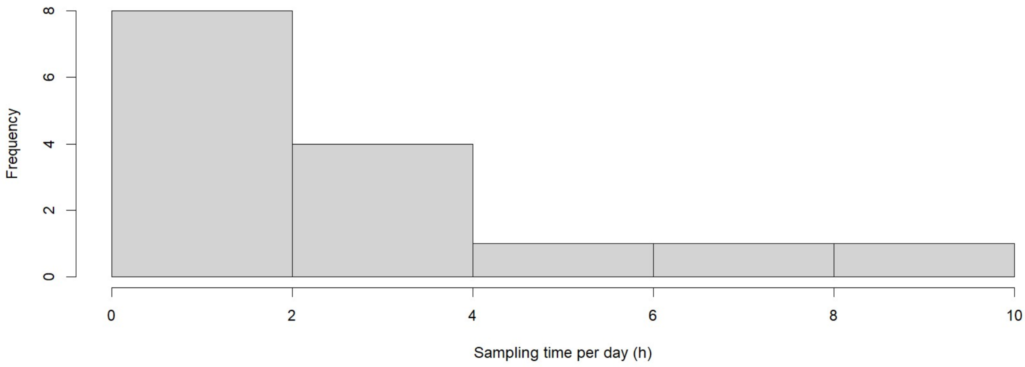


**Figure ESM2.9.** Frequency distribution of daily blood sampling times expressed in hours. 80 % of the samplings were done in less than 4 hours, and 60 % in less than 2 hours (i.e. within a morning).
