## Supplementary material for "Decoupling glycation from mortality: glucose, but not methylglyoxal, reduces survival in zebra finches": ESM4 - Weather

### ESM 4 - Weather

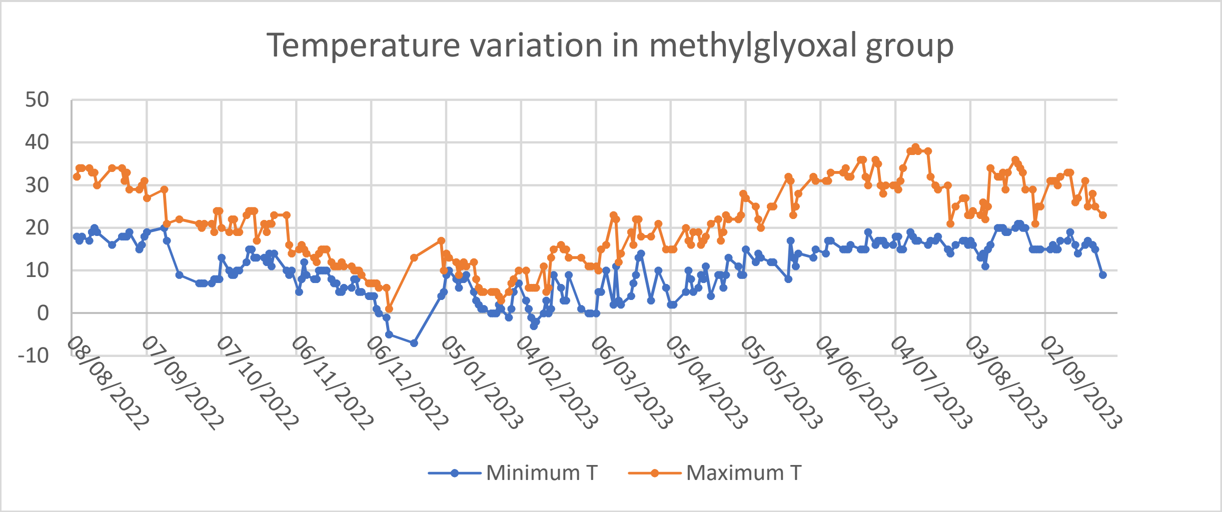


**Figure 1.** Minimum and maximum temperature variation along the year of the experiment in methylglyoxal supplementation group aviary.

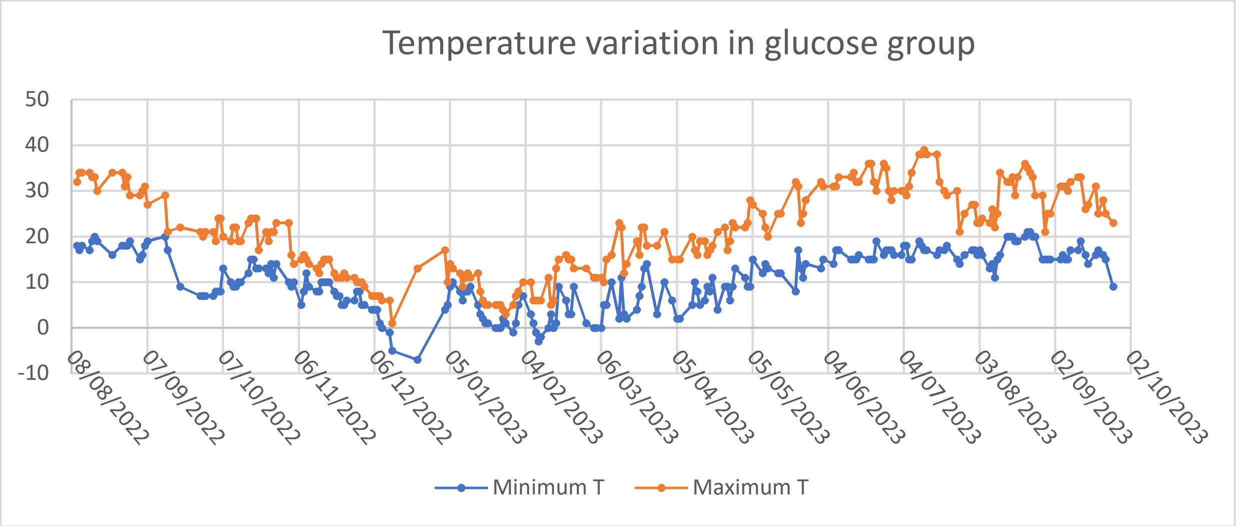


**Figure 2.** Minimum and maximum temperature variation along the year of the experiment in glucose supplementation group aviary.

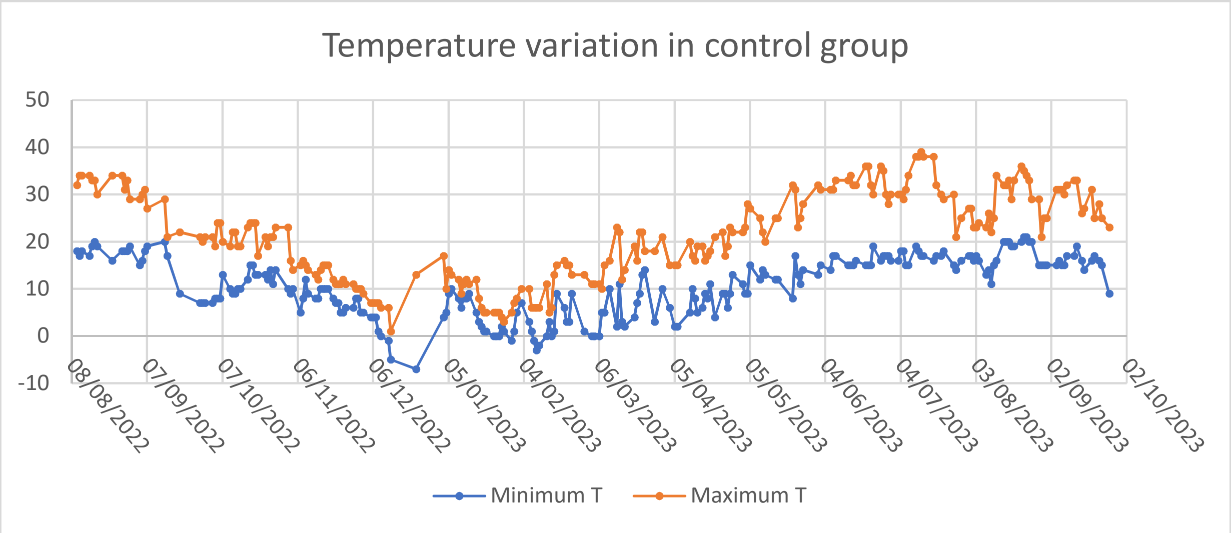


**Figure 3.** Minimum and maximum temperature variation along the year of the experiment in control group aviary.
